# *Atxn2*-CAG100-KnockIn mouse spinal cord shows progressive TDP43 pathology associated with cholesterol biosynthesis suppression

**DOI:** 10.1101/838177

**Authors:** Júlia Canet-Pons, Nesli-Ece Sen, Aleksandar Arsovic, Luis-Enrique Almaguer-Mederos, Melanie V. Halbach, Jana Key, Claudia Döring, Anja Kerksiek, Gina Picchiarelli, Raphaelle Cassel, Frédérique René, Stéphane Dieterlé, Nina Hein-Fuchs, Renate König, Luc Dupuis, Dieter Lütjohann, Suzana Gispert, Georg Auburger

## Abstract

Large polyglutamine expansions in Ataxin-2 (ATXN2) cause multi-system nervous atrophy in Spinocerebellar Ataxia type 2 (SCA2). Intermediate size expansions carry a risk for selective motor neuron degeneration, known as Amyotrophic Lateral Sclerosis (ALS). Conversely, the depletion of ATXN2 prevents disease progression in ALS. Although ATXN2 interacts directly with RNA, and in ALS pathogenesis there is a crucial role of RNA toxicity, the affected functional pathways remain ill defined. Here, we examined an authentic SCA2 mouse model with *Atxn2*-CAG100-KnockIn for a first definition of molecular mechanisms in spinal cord pathology. Neurophysiology of lower limbs detected sensory neuropathy rather than motor denervation. Triple immunofluorescence demonstrated cytosolic ATXN2 aggregates sequestrating TDP43 and TIA1 from the nucleus. In immunoblots, this was accompanied by elevated CASP3, RIPK1 and PQBP1 abundance. RT-qPCR showed increase of *Grn*, *Tlr7* and *Rnaset2* mRNA versus *Eif5a2*, *Dcp2, Uhmk1* and *Kif5a* decrease. These SCA2 findings overlap well with known ALS features. Similar to other ataxias and dystonias, decreased mRNA levels for *Unc80*, *Tacr1*, *Gnal*, *Ano3*, *Kcna2*, *Elovl5* and *Cdr1* contrasted with *Gpnmb* increase. Preterminal stage tissue showed strongly activated microglia containing ATXN2 aggregates, with parallel astrogliosis. Global transcriptome profiles from stages of incipient motor deficit versus preterminal age identified molecules with progressive downregulation, where a cluster of cholesterol biosynthesis enzymes including *Dhcr24*, *Msmo1*, *Idi1* and *Hmgcs1* was prominent. Gas chromatography demonstrated a massive loss of crucial cholesterol precursor metabolites. Overall, the ATXN2 protein aggregation process affects diverse subcellular compartments, in particular stress granules, endoplasmic reticulum and receptor tyrosine kinase signaling. These findings identify new targets and potential biomarkers for neuroprotective therapies.

## Introduction

Within the dynamic research field into neurodegenerative disorders, the rare monogenic variant SCA2 (Spinocerebellar Ataxia type 2) gained much attention over the past decade, since its pathogenesis is intertwined with the motor neuron diseases ALS/FTLD (Amyotrophic Lateral Sclerosis / Fronto-Temporal Lobar Dementia) and with extrapyramidal syndromes such as Parkinson/PSP (Progressive Supranuclear Palsy). Particularly in the past year, it received massive interest since antisense-oligonucleotides were shown to effectively prevent the disease in mouse models, with clinical trials now being imminent. Still, there is an urgent unmet need to define the molecular events of pathogenesis and to identify biomarkers of progression in SCA2 patients, which reflect therapeutic benefits much more rapidly than clinical rating scales, brain imaging volumetry or neurophysiology measures.

SCA2 was first described as separate entity in Indian patients, based on the characteristic early slowing of eye saccadic movements [222]. A molecularly homogeneous founder population of ∼1,000 patients in Cuba made detailed clinical, neurophysiological and pathological analyses possible [5, 8, 45, 137, 161, 167, 218]. The autosomal dominant inheritance was shown to be caused by unstable polyglutamine (polyQ) expansions in the Ataxin-2 gene (*ATXN2*) [153], a stress-response factor conserved from yeast to plants and mammals [7]. The expansion size correlates with younger onset age and faster progression of disease, which manifests with weight loss (later enhanced by dysphagia), muscle cramps and deficient motor coordination [3, 14, 160, 165, 180, 212]. Organisms lacking ATXN2 do not show immediate adverse effects, and are protected from several neurodegeneration variants with tauopathy, such as ALS/FTLD [2, 12, 44, 163, 189]. However, in view of the association of ATXN2-deficiency with progressive obesity, hyperlipidemia and diabetes mellitus [6, 95], it must be asked if such insidiously appearing metabolic excess signs would accompany any neuroprotective treatment by ATXN2 depletion.

PolyQ expansions in different proteins always cause progressive neurodegeneration, via the formation of protein aggregates. However, each polyQ disease shows a different pattern of pathology in the brain [145, 177], so features of each disease protein around the polyQ expansion determine which neuron populations are preferentially vulnerable. ATXN2 is strongly expressed in large neurons and normally localized at the rough endoplasmic reticulum [46, 208] or at plasma membrane sites of receptor tyrosine kinase endocytosis [41, 133, 156], due to protein interactions with PABPC1 and SRC, respectively. Possibly via its influence on endocytosis, an inhibition of mTOR signals, fat stores and cell size was shown for ATXN2 orthologs in yeast, worms and mice [10, 37, 96]. Its role for the ribosomal translation machinery is unclear, since its absence does not change the polysome profile markedly [96]. During periods of cell damage, ATXN2 relocalizes to cytosolic stress granules where RNA quality control is performed, via recruitment of several nuclear proteins such as TIA1 [155]. ATXN2 promotes stress granule formation and decreases P-body size [132]. The physiological interactions of ATXN2 with proteins and RNAs depend on its C-terminal PAM2 motif, its central Lsm and LsmAD domain, and its alternatively spliced proline-rich domain [35, 97, 175, 232], whereas its pathological aggregation is determined by the N-terminal polyQ domain. Currently it is unclear if ATXN2 aggregation occurs within stress granules, to what degree its physiological interactomes are disturbed in SCA2, and which molecular and functional deficits underlie the progressive neurodegeneration, as putative targets of neuroprotective therapies.

In model organisms such as yeast, flies and mice it was shown that ATXN2 depletion by genetic knockout or mRNA-knockdown protects against the neurotoxicity of TDP43, which causes motor neuron degeneration in ALS and in FTLD. In addition, patient studies showed genetic ATXN2 variants to contribute to the risk of ALS/FTLD [12, 44, 58, 94, 99, 170]. Although the molecular mechanisms of ALS/FTLD are not understood, the crucial role of RNA toxicity in its pathogenesis became plain upon the identification of various ALS/FTLD disease genes, such as the RNA binding proteins TDP43 (encoded by the *TARDBP* gene), FUS, HNRNPA2B1, the RNA-particle transporting KIF5A, the RNA toxicity sensor RIPK1, and other stress response factors such as GRN and SOD1 [200].

We recently generated a new SCA2 mouse model via *Atxn2*-CAG100-KnockIn (KIN) [185] and are using it here to obtain initial insights into spinal cord pathology at the molecular and cellular level. Since human autopsy material is very scarce and usually of insufficient quality for RNA studies, these animals provided the first opportunity to elucidate affected pathways and subcellular compartments, without confounding overexpression artifacts. Given that ATXN2 is an RNA-binding protein and that RNA toxicity is central for ALS pathogenesis, an unbiased global transcriptome survey at two different ages was performed.

Overall, our study defines molecular changes that underlie afferent sensory and efferent motor pathology in spinal cord neurons, and provide mechanistic insights into glial changes. At global transcriptome level we identify the most dramatic progression events. Crucially, we demonstrate a severe deficit in steroidogenesis, which will eventually affect all cell types in nervous tissue. A clear overlap with the known pathomechanisms of ALS, as well as other ataxias and dystonias was observed.

## Materials and methods

### Animal breeding

Generation, housing, genotyping and dissection of the *Atxn2*-CAG100-Knock-in mouse [185] and *Atxn2*-Knock-out mice [96] maintained in C57BL/6J background was previously described. The study was ethically assessed by the Regierungspräsidium Darmstadt, with approval number V54-19c20/15-FK/1083.

### Nerve conduction studies, electromyography and quantitative reverse-transcriptase PCR of lower limbs

Recordings were made with a standard electroneuromyograph apparatus (AlpineBiomed ApS, Denmark) in accordance with the guidelines of the American Association of Electrodiagnostic Medicine. Mice at the age of 9-10 months (*n* = 6 for mutants and *n* = 7 for controls) were anaesthetized with 100 mg/kg of ketamine chlorhydrate (Imalgene 1000; Merial, France) and 12 mg/kg of xylazine (Rompun 2%; Bayer HealthCare, Loos, France).

For the tail sensory nerve response, stimulating electrodes placed distally and recording electrodes placed proximally were inserted in the tail 3 cm apart. Sensory nerve responses were elicited by supramaximal square pulses of 0.2 ms duration. Amplitudes (µV) from the sensory nerve-evoked responses were measured and averaged, resulting in one averaged amplitude per animal, which was used for statistical analysis. The latency was measured as the time from the given electrical stimulus to the appearance of a sensory nerve response corresponding to the initial deflection from the baseline.

For the nerve conduction studies, compound muscle action potentials (CMAP) were recorded in *gastrocnemius* muscle as described previously [136]. Briefly, CMAPs were elicited by supramaximal square pulses, of 0.2 ms duration, delivered with a monopolar needle electrode to the sciatic nerve at the sciatic notch level. CMAPs were measured by two monopolar needle electrodes inserted in the *gastrocnemius*, and the system was grounded by subcutaneously inserted monopolar needle electrode in the back of the animal. Amplitudes (mV) from the muscle-evoked responses were measured and averaged, resulting in one average CMAP amplitude per animal, which was used for statistical analysis. The latency was measured as the time from the given electrical stimulus to the appearance of a muscle response—the initial CMAP deflection from the baseline.

For electromyography recording, a monopolar needle electrode (diameter, 0.3 mm; 9013R0313, Alpine Biomed ApS, Denmark) was inserted into the tail of the mouse to ground the system. Recordings were made in the *gastrocnemius* with a concentric needle electrode (diameter, 0.3 mm; 9013S0012, AlpineBiomed ApS, Denmark). Electrical activity was monitored at least 2 min in three different sites. Spontaneous activity was differentiated from voluntary activity by visual inspection.

The RT-qPCR analysis was done in *Tibialis Anterior* and *Soleus* muscles from 4 mice per genotype. Muscles were harvested, rapidly frozen in liquid nitrogen and stored at −80°C until use. Frozen tissues were placed into tubes containing a 5 mm stainless steel bead (Qiagen, Courtaboeuf, France) and 1 ml of Trizol reagent (Life Technologies), and homogenized using a TissueLyser (Qiagen). RNA was prepared from tissue homogenenates following Trizol manufacturer’s instructions. One microgram of total RNA was used to synthesize cDNA using Iscript reverse transcriptase (iscriptTM Reverse Transcription Supermix for RT-qPCR, Bio-Rad) as specified by the manufacturer. Quantitative PCR was performed on a CFX96 Real-time System (Bio-Rad) using iQ SYBR Green supermix (Bio Rad). Three standard genes (H2AC: F-CAACGACGAGGAGCTCAACAAG, R-GAAGTTTCCGCAGATTCTGTTGC/H2AX: F-TCCTGCCCAACATCCAGG, R-TCAGTACTCCTGAGAGGCCTGC/H1H2BC: F-AACAAGCGCTCGACCATCA, R-GAATTCGCTACGGAGGCTTACT) were used to compute a normalization factor using Genorm software v3.5. The following primer sequences were used to assess muscle denervation: AchRalpha (F-CCACAGACTCAGGGGAGAAG, R-AACGGTGGTGTGTGTTGATG), AchRbeta (F-GGCAAGTTCCTGCTTTTCGG, R-CGTCCGGAAGTGGATGTTCA), AchRdelta (F-CGCTGCTTCTGCTTCTAGGG, R-ATCAGTTGGCCTTCGGCTT) and AchRepsilon (F-CAATGCCAATCCAGACACTG, R-CCCTGCTTCTCCTGACACTC).

### Triple immunofluorescence

Mice were anesthetized (Ketaset 300 mg/kg and Domitor 3 mg/kg by i.p. injection), intracardially perfused and tissues were fixed overnight (O/N) in 4% PFA at 4 °C. Samples were frozen, cut with a cryostat and kept in cryoprotection solution at −20 °C until used.

All immunohistochemistry was done in free-floating 30 μm sections. Sections were washed three times for 10 min each in PBS, blocked (5% goat serum in 0.3% Triton X-100/PBS) at RT for 1 h, and incubated with primary antibodies O/N at 4 °C. Samples were washed three times in PBS and incubated with corresponding secondary antibodies and DAPI at RT for 1 h. After three 10 min washing steps in PBS, samples were mounted on SuperFrost Plus slides with Mountant PermaFluor mounting medium and stored at 4 °C.

Immunocytochemistry was performed as described [185], blocking with 5% BSA for 30 min at RT. Primary antibodies were incubated O/N at 4 °C. Images from the different stainings were done with a Nikon Eclipse TE200-E confocal microscope. In all cases, Z-stacks were processed with ImageJ software [178]. Photoshop CS5.1 was used to generate figures.

Primary antibodies used were: ATXN2 (BD Transduction 611378, 1:50), IBA1 (Wako 019-19741, 1:1000), PABPC1 (Abcam ab21060, 1:100); TDP43 (Abcam ab41881, 1:100), TIA1 (Santa Cruz sc-1751, 1:100). Secondary antibodies used were: Alexa Fluor 488 goat anti-mouse IgG A11029, Alexa Fluor 488 goat anti-rabbit IgG A11034, Alexa Fluor 565 rabbit anti-goat IgG A11079, Alexa Fluor 568 donkey anti-sheep IgG A21099, Alexa Fluor 568 goat anti-rabbit IgG A11036 (all Invitrogen, 1:1000).

### Quantitative immunoblots

Protein extraction from tissues and cells was performed with RIPA buffer as described previously [185]. Subsequently, the pellet was re-suspended in SDS lysis buffer (137 mM Tris-HCl pH 6.8, 4% SDS, 20% Glycerol, Proteinase inhibitor, Roche), sonicated and quantified using the Pierce BCA Assay Kit (Thermo Fisher Scientific). The following antibodies were used: ACTB (Sigma A5441, 1:10000), ATXN2 (Proteintech 21776-1-AP, 1:500), CASP3 (Cell Signaling 9665, 1:1000), GFAP (Dako ZO334, 1:2000), GPNMB (Biotechne AF 2330, 1:500), IBA1 (Wako 019-19741, 1:2000), NeuN (Millipore ABN78, 1:1000), PGRN (Biotechne AF 2557, 1:250), PQBP1 (Biomol A302-802A-M, 1:500), RIPK1 (Cell Signaling 3493S, 1:500), TDP43 (Abcam ab41881, 1:1000). The secondary antibodies were: IRDye 680RD goat anti-mouse 926-32220, IRDye 800CW donkey anti-goat 926-32214, IRDye 800CW goat anti-mouse 926-32210, IRDye 800CW goat anti-rabbit 926-32211 (all LI-COR, 1:10000).

### Quantitative reverse-transcriptase PCR in spinal cord

Total RNA isolation from mouse tissues (4 WT vs. 4 KO; 6 WT vs. 5 KIN at 3 months; 7 WT vs. 8 KIN at 14 months) and RT-qPCR were done as described previously [185]. *Tbp* was used as the housekeeping gene. The data were analyzed using the 2^-Ct^ method [179]. TaqMan assays from Thermo Scientific were used for *Aif1* Mm00479862_g1, *Ano3* Mm01270409_m1, *Atxn2* Mm01199902_m1, *C1qa* Mm00432142_m1, *C1qb* Mm01179619_m1, *C1qc* Mm00776126_m1, *C3* Mm01232779_m1, *Casp3* Mm01195085_m1, *Cdr1* Mm04178856_s1, *Cirbp* Mm00483336_g1, *Cybb* Mm01287743_m1, *Cyp46a1* Mm00487306_m1, *Cyp51a1* Mm00490968_m1, *Dcp1a* Mm00460131_m1, *Dcp1b* Mm01183995-m1, *Dcp2* Mm01264061_m1, *Dcps* Mm00510029_m1, *Ddx1* Mm00506205_m1, *Ddx6* Mm00619326_m1, *Dhcr24* Mm00519071_m1, *Dhx15* Mm00492113_m1, *Eif5a2* Mm00812570_g1, *Fmr1* Mm01339582_m1, *Gfap* Mm01253033_m1, *Gpnmb* Mm01328587_m1, *Grn* Mm00433848-m1, *Hmgcs1* Mm01304569_m1, *Hnrnpa2b1* Mm01332941_m1, *Hnrnpd* Mm01201314_m1, *Irak4* Mm00459443_m1, *Kcna1* Mm00439977_s1, *Kcna2* Mm00434584_s1, *Kif5a* Mm00515265_m1, *Lsm1* Mm01600253_m1, *Msmo1* Mm00499390_m1, *Pcbp1* Mm00478712_s1, *Pcbp2* Mm01296174_g1, *Pcbp3* Mm01149750_m1, *Pcbp4* Mm00451991_g1, *Prpf19* Mm01208295_m1, *Ptbp1* Mm01731480_gH, *Pura* Mm01158049_s1, *Puf60* Mm00505017_m1, *Rbfox3* Mm01248771_m1, *Ripk1* Mm00436354_m1, *Rnaset2a/b* Mm02601904_m1, *Scn4b* Mm01175562_m1, *Srrm2* Mm00613771_m1, *Tacr1* Mm00436892_m1, *Tardbp* Mm00523866_m1, *Tbp* Mm00446973_m1, *Tlr3* Mm01207404_m1, *Tlr7* Mm00446590_m1, *Tlr9* Mm00446193_m1, *Trem2* Mm00451744_m1, *Ttbk2* Mm00453709_m1, *Tyrobp* Mm00449152_m1, *Unc80* Mm00615703_m1, *Ybx1* Mm00850878_g1.

### BV2 microglia cell line culture

Murine microglia cell line BV2 [17] was cultured in DMEM supplemented with 10% FBS, 1x L-glutamine and 1x Pen/Strep. BV2 cells were seeded at 5×10^4^ cells/well on poly-D-lysine (0.1 mg/mL) coated glass slides in a 12-well plate O/N and then were stressed with 0.25 mM Sodium Arsenite (NaArs, S7400-100G Sigma-Aldrich) for 15 min. Cells were fixed with 4% PFA at RT for 20 min, permeabilized with 0.1% Triton X-100/PBS for 20 min and washed three times with DPBS before staining.

### Global transcriptomics by Clariom D microarrays

As recommended, 1 μg of RNA was pre-treated with DNase amplification grade (Invitrogen). The GeneChip WT PLUS Reagent Kit (Applied Biosystems) was used to generate single-stranded cDNA (ss-cDNA) following the manufacturer’s instructions. The ss-cDNA was fragmented and labeled right before the hybridization to a Clariom D Array (Thermo Fisher). The arrays were scanned with the Affymetrix GeneChip Scanner and the data were processed with the Transcriptome Analysis Console (TAC) 4.0.1 (Applied Biosystems) using default algorithm parameters.

### STRING protein-protein interaction bioinformatics

The web-server https://string-db.org/ was used in 2019 with version 11.0.

Quantification of cholesterol pathway intermediates by gas chromatography mass spectrometry using selective ion detection methodology:

The procedures were carried out as previously described [194].

### Statistics and graphical visualization

Statistical tests were performed as unpaired Student’s t-test with Welch’s correction using GraphPad Prism software version 7.02. Figures display the mean and standard error of the mean (SEM) values. Significance was assumed at p<0.05 and highlighted with asterisks: p<0.05 *, p<0.01 **, p<0.001 ***, p<0.0001 ****.

## Results

### Neurophysiology of lower limbs finds mainly axonal sensory neuropathy as earliest deficit

The initial characterization of our novel *Atxn2*-CAG100-KnockIn (KIN) mouse mutants had demonstrated a peripheral motor weakness upon forelimb grip strength analysis in comparison to sex-/age-matched wildtype (WT) controls from the age of 11 months onwards [185]. To elucidate if motor neuron denervation and sensory neuropathy are early features within the locomotor phenotype of this SCA2 model as in patients, neurophysiological analyses by hindlimb electroneurography (ENG) were now performed at the age of 9-10 months, when a weight reduction is already evident in males [185]. Sensory potentials showed significant reductions of amplitude (to ∼60%), conduction velocity (SNCV, to ∼80%), and, for males, minimal intensity threshold (to ∼70%) (Figure 1). In contrast, for the compound motor action potentials (CMAP) there was no significant change of velocity, amplitude, minimal and maximal intensity threshold at this disease stage. Upon electromyography (EMG) routines, no muscle denervation potentials such as fasciculations or fibrillations were detected (Suppl. Figure S1A). The reverse-transcriptase quantitative real-time polymerase chain reaction (RT-qPCR) analysis of Acetylcholine Receptor (AchR) alpha / beta / delta / epsilon subunit mRNA expression in the *Tibialis Anterior* and *Soleus* muscle homogenates showed no significant reduction (Suppl. Figure S1B). These observations indicate a loss of some fast-conducting axons and of myelin wrapping that reduce the perception of slight stimuli as initial dysfunction in peripheral nerves, before the advent of later motor deficits. In SCA2 patients, somatosensory denervation of lower limbs is indeed an early phenomenon [169, 212], while the motor neuron pathology at disease onset affects primarily the face, neck and arms [45, 121]. Overall, these data demonstrate sensory neuropathy, but not yet motor denervation of KIN hindlimbs at the age of 9-10 months.

**Figure 1:**
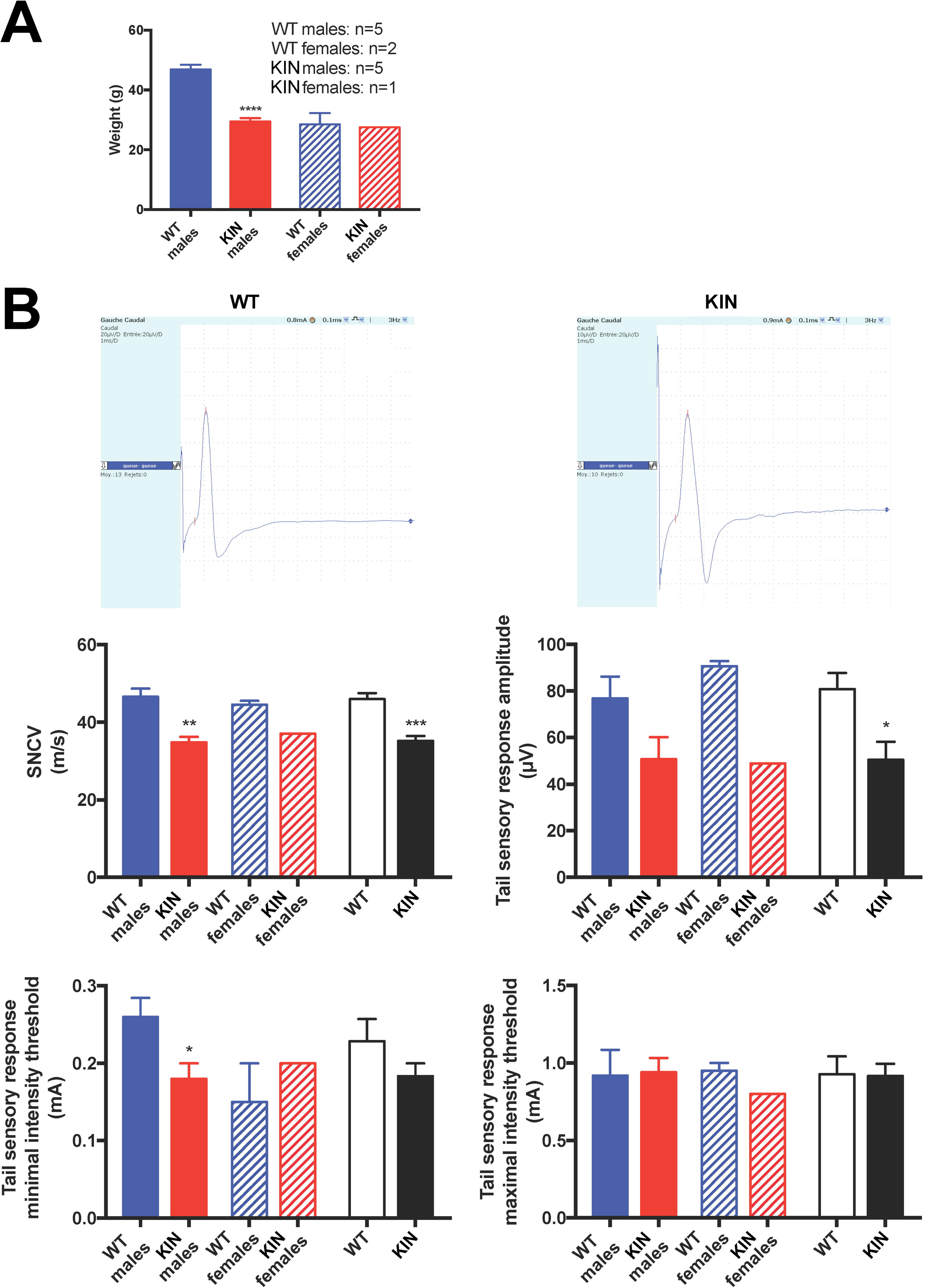
Functional assessment of hindlimb nerve function detects early sensory, predominantly axonal neuropathy. Assessment of 5 male and 1 female *Atxn2*-CAG100-KnockIn (KIN) mice with age-/sex-matched wildtype controls at the age of 9-10 months showed **(A)** a weight reduction with significance for the male mutants. **(B)** Analyses of neurophysiology demonstrated sensory nerve conduction amplitudes and velocity (SNCV) from tail to be reduced and the perception of minimal intensity stimuli to be affected.

### Progressive sequestration of PABPC1 together with nuclear TDP43 and TIA1 proteins into cytosolic ATXN2 aggregates in spinal motor neurons

To further investigate the reduced grip strength in the *Atxn2*-CAG100-KIN mice from the age of 11 months onwards, the spinal cord pathology was examined by histology. Previous analyses of cerebellar Purkinje neurons in 2-year-old mice with KnockIn of a *Atxn2*-CAG42-expansion, a frequent size among SCA2 patients that triggers clinical onset at ages around 30 years, demonstrated a pathological aggregation process where cytosolic ATXN2 sequestrates its direct interactor protein PABPC1 [poly(A)-binding-protein-cytoplasmic-1] into insolubility [35, 185]. In an effort to test this observation in our new *Atxn2*-CAG100-KIN mice for spinal cord motor neurons, triple immunofluorescence was employed to study animals at the early adult age of 3 months, at the age of initial weight deficits around 6 months, and the prefinal age of 14 months. As shown in Figure 2, the yellow-stained colocalization of green ATXN2 signals and red PABPC1 signals within cytosolic clumps were clearly visible at 6 months and became more numerous by 14 months. Additional detection of nuclear TDP43, which is known to interact with ATXN2 indirectly via RNA-association within short-lived stress granules [44, 236], also demonstrated colocalization within the cytosolic ATXN2 aggregates, albeit only at later age to a massive degree (Figure 2). This observation provides evidence of progressive spinal motor neuron affection leading to a motor neuropathy in this SCA2 mouse model over time. A similar pathological relocalization from a nuclear position to the cytosolic aggregates was also observed for the RNA-binding protein TIA1 (Suppl. Figure S2), an established marker of stress granules [47]. As a neuropathological diagnostic hallmark of spinal motor neuron affection in ALS, the nuclear depletion and cytoplasmic accumulation of TDP43, TIA1 and other RNA-binding proteins is probably reflecting a crucial disease mechanism [31, 88, 103]. The TDP43 mislocalization to the cytosol was already reported for SCA2 patients in lower and upper motor neurons, cerebellar Purkinje cells, brainstem neurons and other neuron populations [11, 44, 206], so the animal model provides a faithful reflection of the temporal development and the spatial distribution of the human pathology.

**Figure 2:**
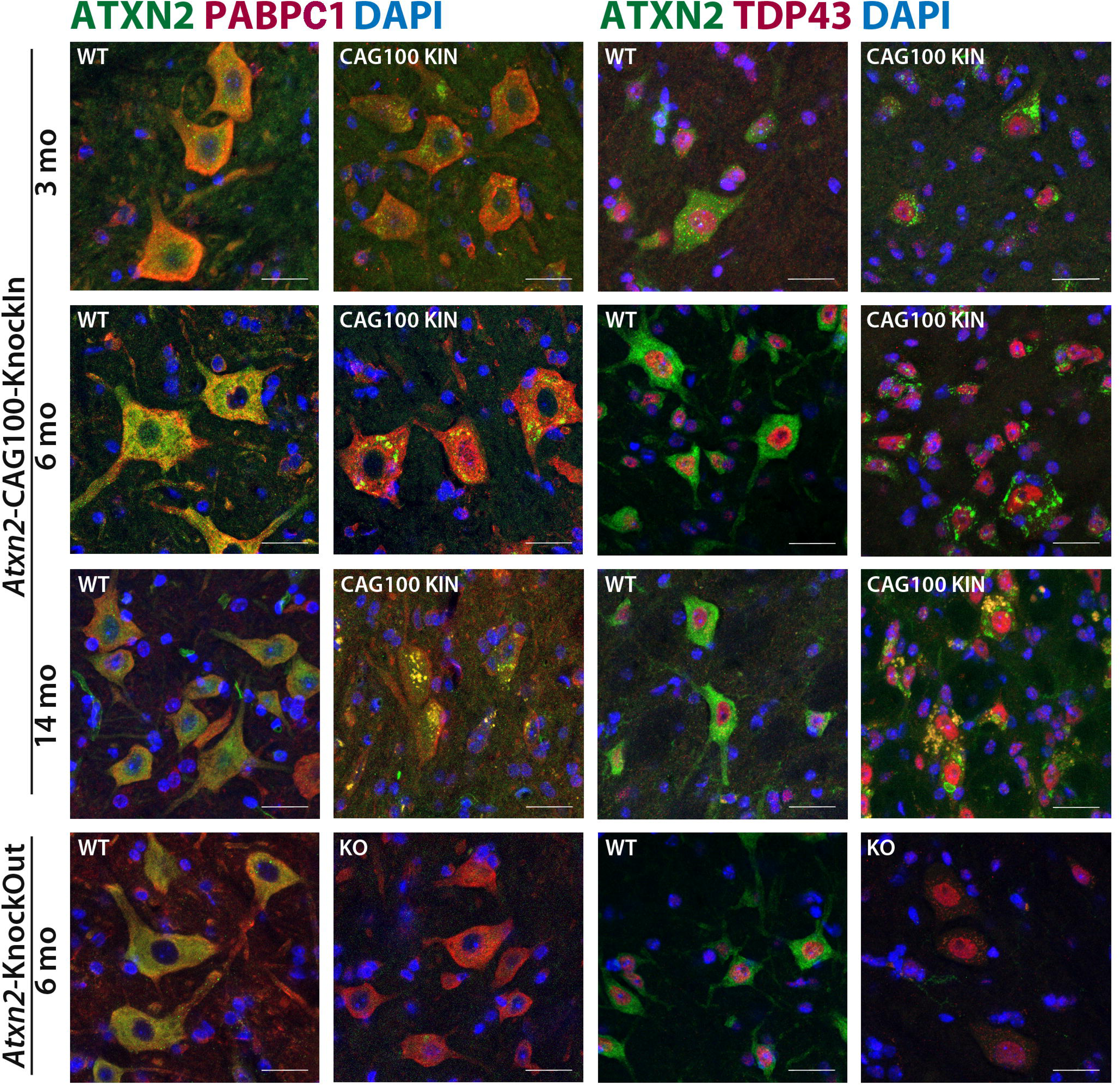
ATXN2 protein aggregates in spinal motor neurons progressively sequestrate PABPC1 and TDP43. Triple co-immunofluorescent staining of ATXN2 (green) versus PABPC1 or TDP43 (red) in *Atxn2*-CAG100-KIN (at 3, 6 or 14 months of age) versus *Atxn2*-KO mice (at 6 months) versus age-/sex-matched WT controls. The sequestration process co-localizes both proteins in cytosolic foci and produces a yellow signal in the merged panels. Nuclei were detected by DAPI (blue color), scale bar reflects 25 µm.

### Cascade of molecular events upstream from TDP43 mislocalization is mirrored in spinal cord

The importance of TDP43 mislocalization for ALS and FTLD has prompted numerous studies into the underlying molecular events, so we investigated whether the pathomechanisms upstream from TDP43 aggregation can also be found in the *Atxn2*-CAG100-KIN spinal cord homogenates. It was reported that ATXN2 expansions trigger stress-dependent activation of caspase-3 (CASP3), which is responsible for the proteolytic C-terminal cleavage and cytoplasmic retention of TDP43 [67]. To test whether this pathogenesis feature is present, and if it is dependent (i) on the physiological function of ATXN2, (ii) on the polyQ expansion or (iii) on disease progression, we studied spinal cord from mice with *Atxn2*-KO versus *Atxn2*-CAG100-KIN at the presymptomatic age of 3 months versus the preterminal age of 14 months. Indeed, quantitative immunoblots demonstrated an elevated abundance of TDP43 (1.6-fold) and CASP3 protein (1.8-fold) only in KIN mice at the old age, while their mRNA retained normal expression (Figure 3A/B). Thus, the elevated abundance of both proteins occurs only in polyQ expansion tissue during the progressive aggregation process, in good agreement with our immunohistochemical demonstration of TDP43 deposits in the cytosol. At the age of 14 months the neuronal mass appeared already reduced below >0.6-fold (as estimated via quantitative immunoblots of the marker NeuN, Figure 3A), in parallel to a similar 2-fold increase of astrogliosis (marker GFAP, Figure 3A and 3B) and together with substantial microgliosis (marker IBA1, encoded by *Aif1* mRNA, Figure 3B).

**Figure 3:**
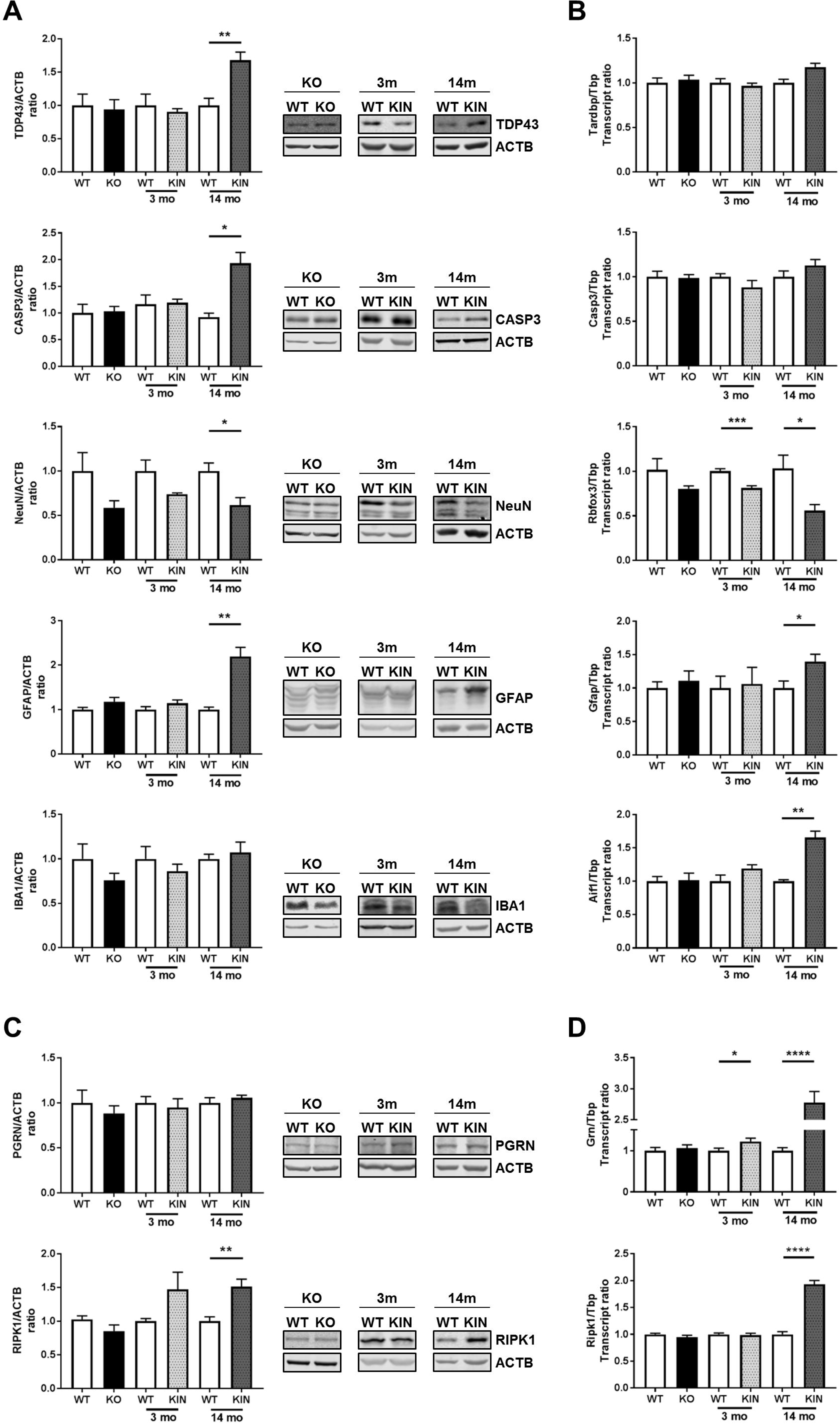
Validation of protein and mRNA level dysregulations. **(A)** Quantitative immunoblots confirmed neuronal loss (marker NeuN), astrogliosis (marker GFAP) and microgliosis (marker IBA1) to occur in *Atxn2*-CAG100-KIN spinal cord at the preterminal stage of 14 months age, but not at the early KIN stage of 3 months age and in the *Atxn2*-KO at 6 months. Significantly increased levels in KIN at 14 months were also shown for TDP43 and the factor responsible for its cleavage, CASP3. **(B)** Quantitative RT-PCR analyses showed a significant deficit of NeuN transcript (*Rbfox3*) already at incipient disease stage in 3-month-old KIN, whereas astrogliosis (marker *Gfap*) and microgliosis (marker IBA1 transcript *Aif1*) became significant at late state. Protein abundance **(C)** and transcript levels **(D)** were also documented for PGRN (encoded by *Grn* mRNA) as molecular marker of lysosomal activation and atrophy, as well as RIPK1 as molecular marker of RNA-toxicity and necroptosis. Again, a significant elevation of *Grn* mRNA at the age of 3 months suggested atrophy and lysosomal breakdown to occur in parallel with first locomotor deficits, predating necroptotic cell death.

Furthermore, mutations of progranulin (PGRN, encoded by *Grn* mRNA) were demonstrated in FTLD individuals to cause cytoplasmic TDP43 mislocalization [154]. The depletion of progranulin also leads to accumulation of TDP43 fragments, in a caspase-dependent manner [239]. Intracellular PGRN gives rise to processed secreted isoforms. They stabilize the protease CTSD in lysosomes and extracellularly to stimulate axonal outgrowth [13]. The absence of PGRN conversely acts as trigger of synaptic pruning by microglia via the complement membrane attack complex [111]. It was therefore interesting to observe significantly increased *Grn* mRNA levels in 14-month-old KIN spinal cord (Figure 3C/D) as a quite specific response to the TDP43 pathology, possibly representing a compensatory effort.

PGRN abundance is modulated in response to RIPK1-mediated phosphorylation [118], whose activation can sense viral or toxic nucleotides via the toll-like receptor TLR3 [93, 129], drives pro-inflammatory necroptosis [234], and is implicated in ALS [81, 230]. Consistent with these reports, enhanced RIPK1 protein levels were observed in the spinal cord of *Atxn2*-CAG100-KIN mice, possibly with a 1.4-fold abundance already at 3 months, but achieving significance only for the 1.5-fold increase at the age of 14 months (Figure 3C/D). The *Ripk1* mRNA levels showed a clearly significant increase to almost 2-fold at the age of 14 months.

Toll-like-receptor (TLR)-triggered activation of microglial cells, as well as inflammatory lysis of synapses via neuronal deposition of complement C1q and C3 factors on their membranes, were carefully documented in ALS spinal cord tissue [29, 143]. Again, in the 14-month-old spinal cord of *Atxn2*-CAG100-KIN mice there were strong transcriptional inductions for *Tlr3*, *Tlr7*, *Tlr9*, *C1qa*, *C1qb, C1qc* and *C3* (Suppl. Figure S3A). The particularly strong induction of *Tlr7* as sensor (at endosomes or lysosomes) for toxic single-stranded RNA and miRNA (3.5-fold), in comparison to *Tlr9* as sensor of single-stranded DNA sequences (2.3-fold) and *Tlr3* as sensor of double-stranded RNA (1.3-fold) underlined and elucidated the RNA-toxicity [68, 74, 100] present in the spinal cord tissue of our SCA2 model. It is interesting to note that Ataxin-2 domains were reported to play a role in the miRNA-mediated mRNA degradation [90] and to interact with the microRNA effector DDX6 to mediate quality control of mRNAs [4, 120, 132, 198].

Finally, it is known that non-expanded polyQ-domain containing proteins are being recruited into the aggregates formed by expanded ATXN2 or expanded ATXN3 [149, 207]. Other proteins also bind specifically to expanded polyQ domains, e.g. PQBP1 (Polyglutamine binding protein 1) as a sensor of viral or toxic nucleotides within the innate immunity pathway of brain cells [224, 231]. Indeed, in 14-month-old *Atxn2*-CAG100-KIN spinal cord a significant accumulation was documented (Suppl. Figure S3B). Overexpression of PQBP1 in transgenic mice results in a loss of spinal motor neurons as well as a loss of cerebellar Purkinje neurons [135]. Deletion mutations in PQBP1 affect the turnover of FMRP (fragile-X mental retardation protein) in neuronal RNA granules and trigger synaptic dysfunction [238].

Jointly, these analyses of protein abundance or mRNA expression demonstrate a cascade of abnormal molecular events, where interactions of toxic RNAs affect endosomal TLRs and cytosolic PQBP1, which act as pathology sensors together with RIPK1, affecting PGRN-dependent neurite growth and CASP3-modulated TDP43 aggregation. This molecular cascade was defined for the motor neuron pathology in ALS and it is observed also in the spinal cord of our SCA2 model at advanced stages.

### Activated microglia contains ATXN2 aggregates and is mainly pro-inflammatory, possibly also by cell-autonomous affection

Upon close inspection of spinal cord immunohistochemistry, in the 14-month-old *Atxn2*-CAG100-KIN mice some ATXN2 aggregates appeared positioned outside motor neurons. Triple immunofluorescence staining showed them to colocalize with the microglia marker IBA1 and these microglia cells showed pronounced activation with larger cell size and thicker branches (Figure 4A). Within their cytoplasm, beyond the ATXN2 aggregates also a diffuse ATXN2 staining was visible, compatible with the known stress-dependent induction of Ataxin-2 expression [96]. Indeed, publically available RNAseq findings in diverse brain cell populations from murine and human samples (http://brainrnaseq.org/) documented Ataxin-2 transcript expression to be similar in neurons and astrocytes, while Ataxin-2 transcript expression in oligodendrocytic, microglial and endothelial cells ranges at 30-50% in comparison to neurons. To assess this issue at the protein level, the microglial cell line BV2 was immunocytochemically stained for ATXN2 and its direct interactor protein PABPC1, while exposure to oxidative stress via NaArs administration was used to trigger the formation of characteristic stress granules, with the relocalization of ATXN2 with PABPC1 there. The immunofluorescent signals confirmed the typical diffuse cytosolic pattern for both proteins that changes to multiple granules with the expected delay (Figure 4B). These data indicate that microglia would not only phagocytose ATXN2 aggregates that were produced by neurons, but may also suffer in cell-autonomous manner from the RNA toxicity triggered by the ATXN2 mutation. To address the question whether the activated microglia cells in old KIN spinal cord are anti-inflammatory protective (M2 differentiation) or rather show the pro-inflammatory toxic properties (M1 differentiation) that were documented in ALS [56], expression of the phagocytosis factor TREM2 and its adaptor DAP12 (encoded by *Tyrobp*) as M2 markers, as well as the common downstream TLR signaling element IRAK4 and the respiratory burst oxidase subunit NOX2 (encoded by *Cybb* mRNA) as M1 marker [25, 85, 147, 240] was quantified by RT-qPCR. While *Trem2* and *Tyrobp* showed about 3-fold induction and *Irak4* exhibited 1.5-fold induction, the superoxide-production enzyme *Cybb* was elevated 7-fold (Suppl. Figure S4), indicating a massive pro-inflammatory activity of microglial cells.

**Figure 4:**
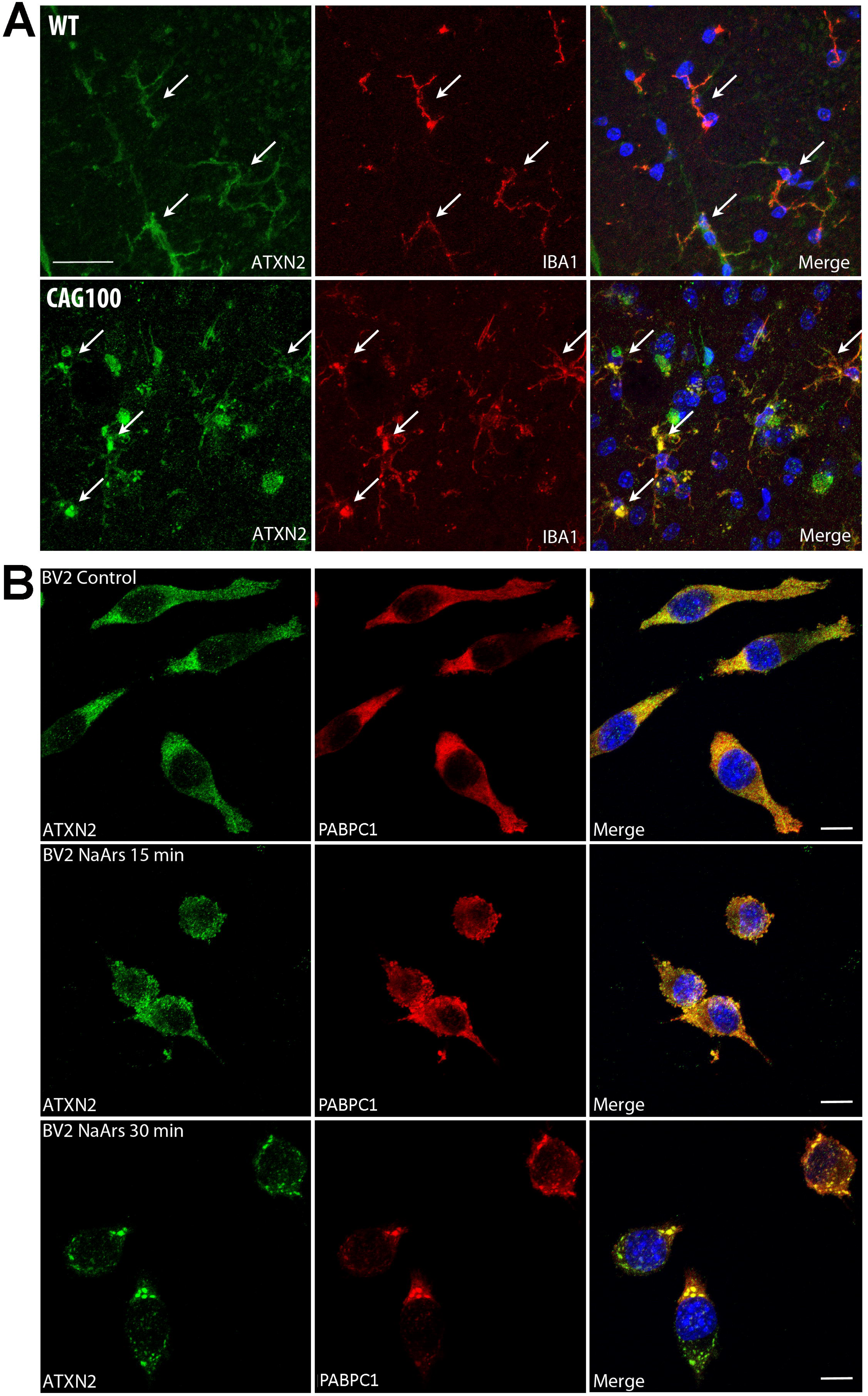
Microglia is affected by ATXN2 aggregates in spinal cord and by cell-autonomous expression of ATXN2 *in vitro*. Triple immunofluorescence of ATXN2 (green color) and IBA1 (red) in **(A)** spinal cord sections of 14-month-old *Atxn2*-CAG100-KIN versus sex-/age-matched shows anti-ATXN2 positive foci colocalizing with IBA1 signals of microglia cells. Scale bar represents 25 µm. In the microglial BV2 cell line **(B)**, diffuse anti-ATXN2 immunoreactivity in the cytosol showed progressive condensation into separate foci after 30 min NaArs treatment, in colocalization with PABPC1 signals, as typical stress granule features. In the merged panels, nuclei are visualized stained with DAPI (blue). Scale bar represents 50 µm.

Altogether, the strong microglial activation in spinal cord at prefinal stages does not only serve neuroprotective functions via ATXN2 aggregate phagocytosis, but shows prominent toxic pro-inflammatory features, possibly via cell-autonomous sensitization of microglia to RNA toxicity caused by the stress-augmented expression of expanded ATXN2.

### Expression profiles of spinal cord at incipient versus preterminal disease stage consistently show increases for RNA-binding proteins and bioenergetics markers, decreases for tyrosine kinase signaling factors, synapse/axon components, and cholesterol biosynthesis enzymes

In view of the crucial role of ATXN2 and stress granules for RNA quality control, we surveyed the global transcriptome with Clariom D microarrays containing >214,000 spotted oligonucleotides, which represent almost every exon from coding mRNAs, microRNAs and long non-coding RNAs. This was first done in spinal cord tissue from animals at the age of 10 weeks when initial deficits of weight and spontaneous motor activity become apparent [185], and then again at the preterminal stage of 14 months when maximal expression dysregulations are expected but may already be confounded by altered cellular composition. Figure 5 illustrates the general approach, filtering criteria and a total overview of significant anomalies. It is important to state that we filtered and assessed all 1.2-fold expression dysregulations with nominal significance, if they were components of a pathway that showed enrichment after correction for multiple testing. The reasoning behind the screening of so subtle changes comes from research into Parkinson’s disease (PD), where 2-fold dosage increase of the disease protein alpha-synuclein was shown to trigger disease onset at ages of 30 years, 1.5-fold increase causes onset around 50 years, and 1.3-fold increase starts disease after 70 years of age [19, 34, 191]. The survey of 1.2-fold global transcriptome dysregulations in a PD mouse model correctly discovered neuroinflammatory changes as initial molecular pathology, which were later found to be crucial for a successful rescue [193, 205]. Given that ATXN2 mutations also trigger mitochondrial dysfunction as in PD, and that SCA2 may manifest with a Parkinsonian phenotype [142, 181, 186, 187, 223], we assessed here if distortion of such pathways becomes relevant by subtle dysregulations at several points.

**Figure 5:**
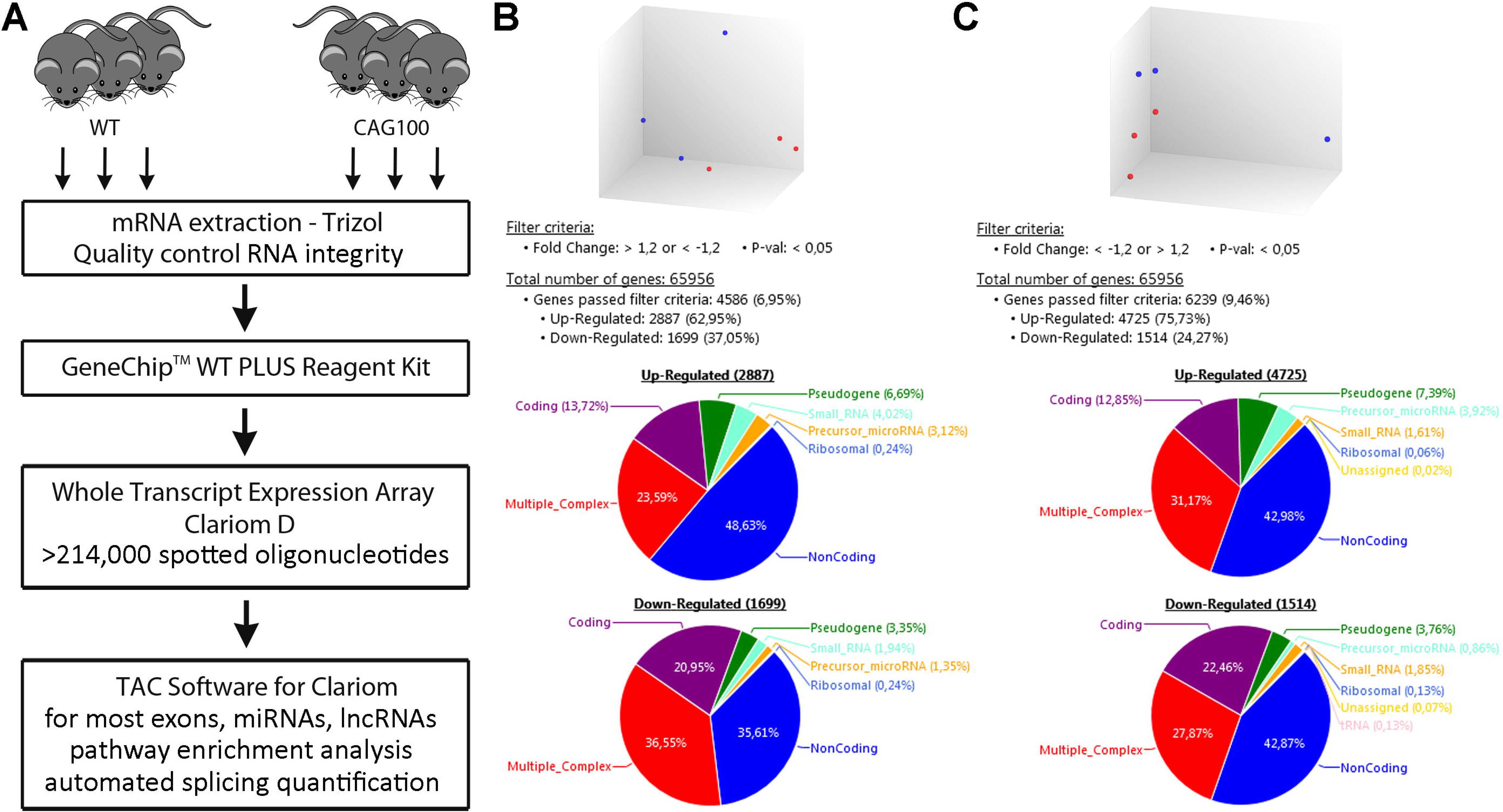
Global transcriptome profiles of spinal cord at presymptomatic versus prefinal stage. **(A)** This scheme depicts the general approach to survey molecular impact of polyQ-expansion in the RNA-binding protein ATXN2 at the global transcriptome level in unbiased manner. Spinal cord tissue from 3 homozygous *Atxn2*-CAG100-KIN versus 3 age-/sex-matched WT mice was used to extract high-integrity RNA. Hybridization signals were quantified from >214,000 oligonucleotides representing practically each exon from coding mRNAs, as well as microRNAs and lncRNAs. Automated statistics and pathway enrichment analysis was performed by the Clariom Transcriptome Analysis Console (TAC). **(B)** Results at the age of 10 weeks showed a clear separation of WT and mutant profiles upon principal component analysis (above, WT red dots, KIN blue dots) and significant expression dysregulation by >20% for 6.95% of all genes. **(C)** Results at the age of 14 months also showed a clear separation of WT and mutant profiles upon PCA. Expression dysregulation affected 9.46% of all genes, with a strongly increased number of upregulations.

At incipient disease stage of the *Atxn2*-CAG100-KIN mouse, a 1.2-fold expression change was observed for 1,699 downregulations versus 2,887 upregulations; at preterminal age, the number of 1,514 downregulations appeared quite similar, but an increased number of 4,725 upregulations reflected the disease progression. The individual factors in their interaction clusters were visualized in STRING diagrams (Suppl. Fig. S5). Initially the upregulations were prominent (Suppl. Fig. S5A) for the ATXN2 interactor *Pabpc1* mRNA, together with various other translation initiation and spliceosomal factors, in parallel to upregulations of acetylation and mitochondrial factors. This was accompanied by initial downregulations (Fig. S5B) for receptor tyrosine kinases that signal via the ATXN2-interactors SRC and GRB2, together with depletions for their downstream effectors, various MAP kinases and CAM kinases. Downregulations included pathway enrichments for potassium channels, synapse/axon factors, and cholesterol biosynthesis enzymes (statistics in Suppl. Table S1).

**Table 1:**
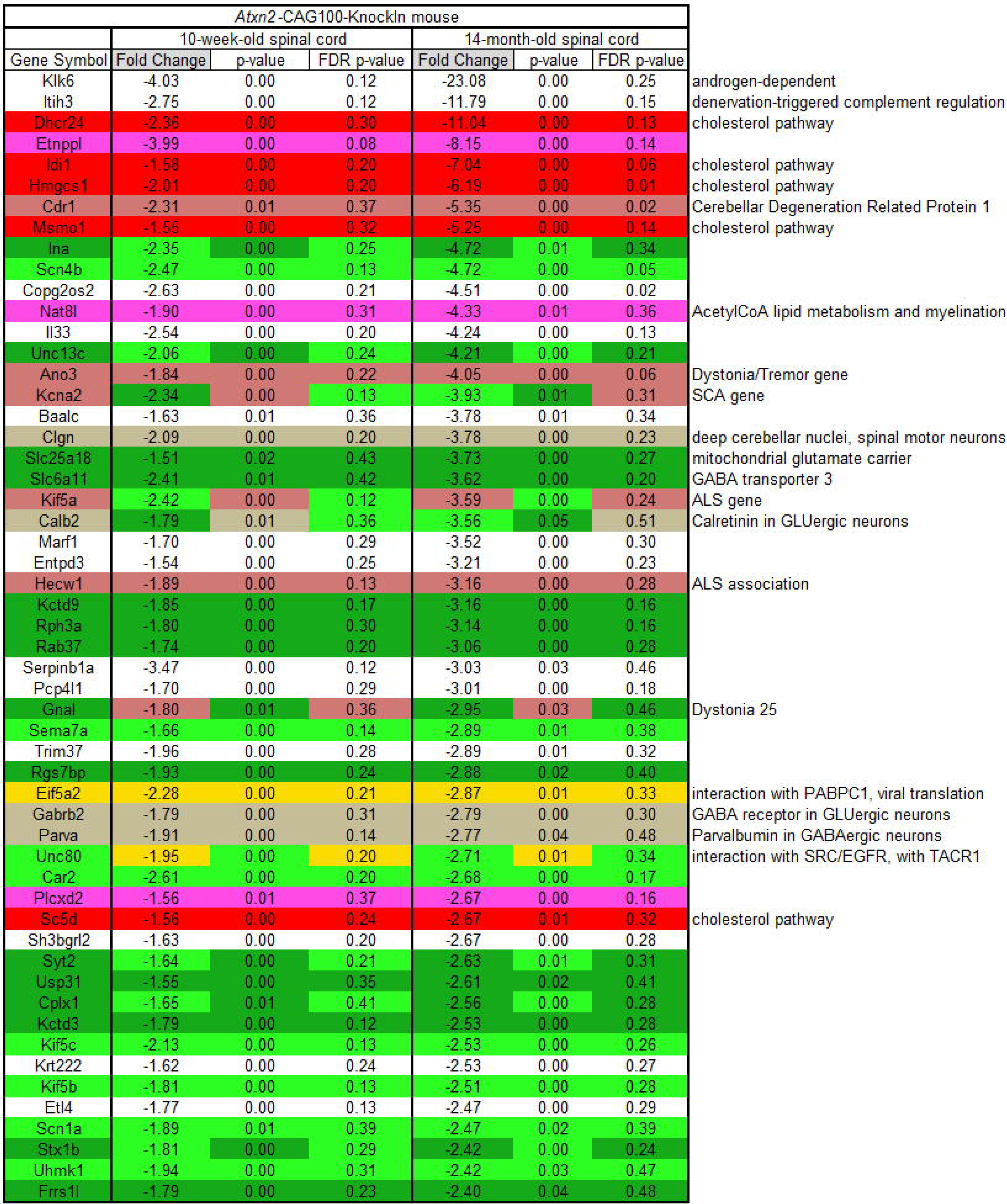
List of coding transcripts with dysregulation effect doubling over KIN lifespan. . Candidate progression markers were selected from global transcriptome profiles upon >1.2-fold expression change at 10 weeks versus ≥2.4-fold change at 14 months of age in *Atxn2*-CAG100-KIN spinal cord. All such factors were downregulated, as illustrated by negative fold change effects. Significances shown were calculated by microarray Clariom D Transcriptome Analysis Console. Pathway component clusters are highlighted by colors, using red for cholesterol biosynthesis, purple for lipid metabolism, light green for axon, dark green for presynapse, gold for members of the ATXN2 interactome, rose for factors responsible for neurodegenerative disorders, beige for markers of distinct neuron populations. Authors’ comments on pathway functions of disease implications are provided at the right margin.

To reduce complexity at preterminal stage, only 2-fold expression changes were evaluated by STRING. The strong upregulations included immune defense and lysosome components, beyond the RNA-binding and bioenergetics proteins (Suppl. Fig. S5C). The strong downregulations again reflected deficits of potassium channels, synapse/axon factors, and cholesterol biosynthesis enzymes (Suppl. Fig. S5D; statistics in Suppl. Table S2).

### Pathway enrichments in the Transcriptome Analysis Console relate to ATXN2 interactome

Bioinformatics via the Affymetrix Clariom Transcriptome Analysis Console (TAC) recognized enrichments for apparently diverse pathways at incipient stage (Suppl. Figure S6) and preterminal stage of disease (Suppl. Figure S7). As common denominator, all these pathways reflect progressive expression adjustments at the diverse sites of physiological ATXN2 localization in young unstressed tissue: ATXN2 was previously shown to interact with the EGF / insulin receptor and downstream AKT/MAPK signaling [41, 95, 133], to inhibit the mTORC1 growth complex in three species [10, 37, 96], to reside at the endoplasmic reticulum (ER) [36, 208] where cholesterol biosynthesis and calcium storage occur [62, 65, 107, 209], to modulate ribosomal mRNA translation [46, 138, 175], to interact with actin/actinin [105, 174] and to modulate adipogenesis [95, 185]. Thus, both transcriptome profiles are compatible with the concept that expanded ATXN2 irritates its various subcellular interactomes and triggers many gradual expression adaptations that may compensate most, but not all dysfunctions. Apart from RNA toxicity, the disruptions of cholesterol biogenesis and calcium dynamics at the ER have been proposed as crucial disease mechanisms in ALS [15, 210].

### Expression profiles of spinal cord at preterminal disease stage reveal affected RNA toxicity factors, ALS-ataxia-dystonia genes, as well as cholesterol biogenesis repression

Pathway analyses did not highlight and expression anomalies for the typical stress granule factors, even in the presence of old age and inflammation stress. Despite several publications implicating ATXN2 in micro/non-coding RNAs and implicating TDP43 in splicing of RNAs [120, 203], no strong progression effects were observed for non-coding or alternatively spliced RNAs at preterminal age. Thus, efforts were made to survey individual factors that are prominent, due to their massive dysregulation or through their known roles for disease pathogenesis.

The strongest single effect at incipient stage was on *Etnppl* mRNA (−3.9 fold, false discovery rate FDR q-value 0.0795), which encodes an enzyme responsible for the breakdown of membrane phospholipids to acetaldehyde. The strongest effects at preterminal stage included increases for microglial (*Gpnmb,* 261-fold, q=9e-05) and lysosomal activation (*Atp6v0d2,* 9-fold, q=0.01), versus reductions for the cholesterol biosynthesis enzyme *Hmgcs1* (−6.2-fold, q=0.01), the ataxia disease gene *Cdr1* (−5.3-fold, q=0.01), the ER-Golgi traffic coatomer factor *Copg2os2* (−4.5-fold, q=0.01), and the action potential modulator *Scn4b* (−4.7-fold, q=0.05).

In additional bioinformatics efforts to elucidate molecular underpinnings of prefinal pathology, volcano plots were generated that illustrate the significance and fold-change for prominent factors. Key factors in RNA toxicity such as *Rnaset2* and *Tlr7* were upregulated (Suppl. Fig. S8A); ALS disease genes like *Grn* and *Hnrnpa2b1* were also upregulated, while *Kif5a* showed a downregulation (Suppl. Fig. S8B); among the ataxia disease genes, two enzymes responsible for very-long-chain fatty acid elongation (*Elovl4/5*) and three potassium channels (*Kcnc3*, *Kcnj10*, *Kcna1*) were downregulated (Suppl. Fig. S8C); we have previously reported the deficit of *Elovl4/5* expression and of its metabolic product C24 sphingomyelin in the CAG100-KIN mouse, as well as the loss of C24 sphingomyelin in the cerebellum of a SCA2 patient [184]. Interestingly, in the CAG100-KIN spinal cord expression profile (Suppl. Fig. S8C), significance was just achieved for the downregulation of *Ttbk2*, as a factor with causal role in tauopathies and TDP43 phosphorylation [201].

### Strongest progression of dysregulation for cholesterol biosynthesis and synapse/axon factors

Next, we attempted to identify molecular biomarkers of disease progression by selecting the ≥1.2-fold expression changes in 10-week-old spinal cord, which evolved into ≥2.4-fold dysregulations in the 14-month-old tissue. Table 1 shows the 51 factors that were defined by this approach. They were studied in great detail with respect to pathway enrichment, partially validated at RT-qPCR and immunoblot level, and assessed regarding enzyme metabolic effects. We prioritized them in view of the high importance of progression markers for the understanding of pathogenesis and the evaluation of neuroprotective therapies.

Table 1 uses red color to highlight the novel and crucial observation that factors of cholesterol metabolism are clustering among the strongest repression effects. This was evident upon automated STRING bioinformatics (Suppl. Fig. S5E; Suppl. Table S3). In addition, the androgen- (a cholesterol-derived hormone) responsive *Klk6* expression was downregulated [233]. Similarly, the *Etnppl* mRNA was decreased, and its activity is triggered by corticosteroids, which are derived from cholesterol [48]. It is noteworthy that the mRNA encoding the enzyme *Nat8l* is downregulated as well, reflecting the insidious depletion of the abundant brain metabolite N-acetyl-aspartate (generated in neuronal mitochondria to shuttle metabolic substrates for oligodendroglia myelination), as an established imaging biomarker of advanced disease in SCA2 patients and in our mouse model [185]. Together, these findings suggest progressive and prominent deficits in acetyl-CoA supply, membrane phospholipid metabolism, myelin lipid synthesis, cholesterol biogenesis and hormone homeostasis.

### Quantification of cholesterol biogenesis pathway intermediate metabolites confirms significant deficits that cannot be compensated

We wondered whether the cholesterol pathway dysregulation is a primary event that contributes to neurodegeneration or whether it is secondary, a consequence of membrane breakdown. The transcriptional downregulations of various enzymes in cholesterol biogenesis and turnover were confirmed by RT-qPCR (Suppl. Fig. S9). Are the mRNA levels of these enzymes repressed, because excessive amounts of free cholesterol are available in the aftermath of synapse and axon loss? Or is this transcript deficit responsible for cholesterol depletion? To assess this question, the intermediary metabolites of cholesterol metabolism were quantified by gas chromatography-mass spectrometry using selective ion detection methodology in spinal cord tissue at the age of 14 months. Conversely to the significant increase of cholesterol in *Atxn2*-KO mouse blood [95], a significant 0.76-fold decrease of cholesterol was documented (Suppl. Table S4). Also its degradation products 24OH-cholesterol and 27OH-cholesterol showed significant 0.80-fold and 0.60-fold deficits, respectively (Suppl. Table S4). Other known deficits of cholesterol biogenesis such as the Smith-Lemli-Opitz syndrome or desmosterolosis have a phenotype that includes mental retardation due to brain atrophy, with the membrane cholesterol deficit being partially compensated by the integration of precursor metabolites [98, 151, 176, 225]. In contrast, in our SCA2 mouse model the precursor metabolites showed massive deficits at multiple steps in both the Bloch pathway and the Kandutsch-Russell pathway, with 0.08-fold reduction for lanosterol at the beginning of post-squalene synthesis, and 0.09-fold reduction for lathosterol towards the end of the biosynthesis cascade (Figure 6). Comparisons between the cholesterol enzyme expression deficits listed Table 1 and these metabolite quantifications demonstrate an excellent correlation of cholesterol precursor metabolite deficits (Figure 6). Furthermore, the significant gradual downregulations of the biosynthesis enzymes *Sqle*, *Sc5d*, *Nat8l*, *Etnppl* and *Plcxd2* indicate that cholesterol synthesis is affected already in the pre-squalene pathway and that the basal homeostasis of Acetyl-CoA and membrane phospho-lipids is also altered. These findings are in good agreement with global proteome profiles of *Atxn2*-null organisms, which demonstrated a prominent alteration of the breakdown of fatty acids and amino acids to Acetyl-CoA within mitochondria of mice, and demonstrated a prominent affection of the citric acid cycle in yeast [122, 182], jointly emphasizing a metabolic role for ATXN2. Altogether, expression profiles at initial and late disease stages are corroborated by lipid quantification studies, pointing to significant deficiencies of cholesterol and its precursors in the nervous tissue as a primary event of pathogenesis, which occurs so early and progresses so strongly that this pathway can serve as a sensitive and specific read-out in therapeutic trials.

**Figure 6:**
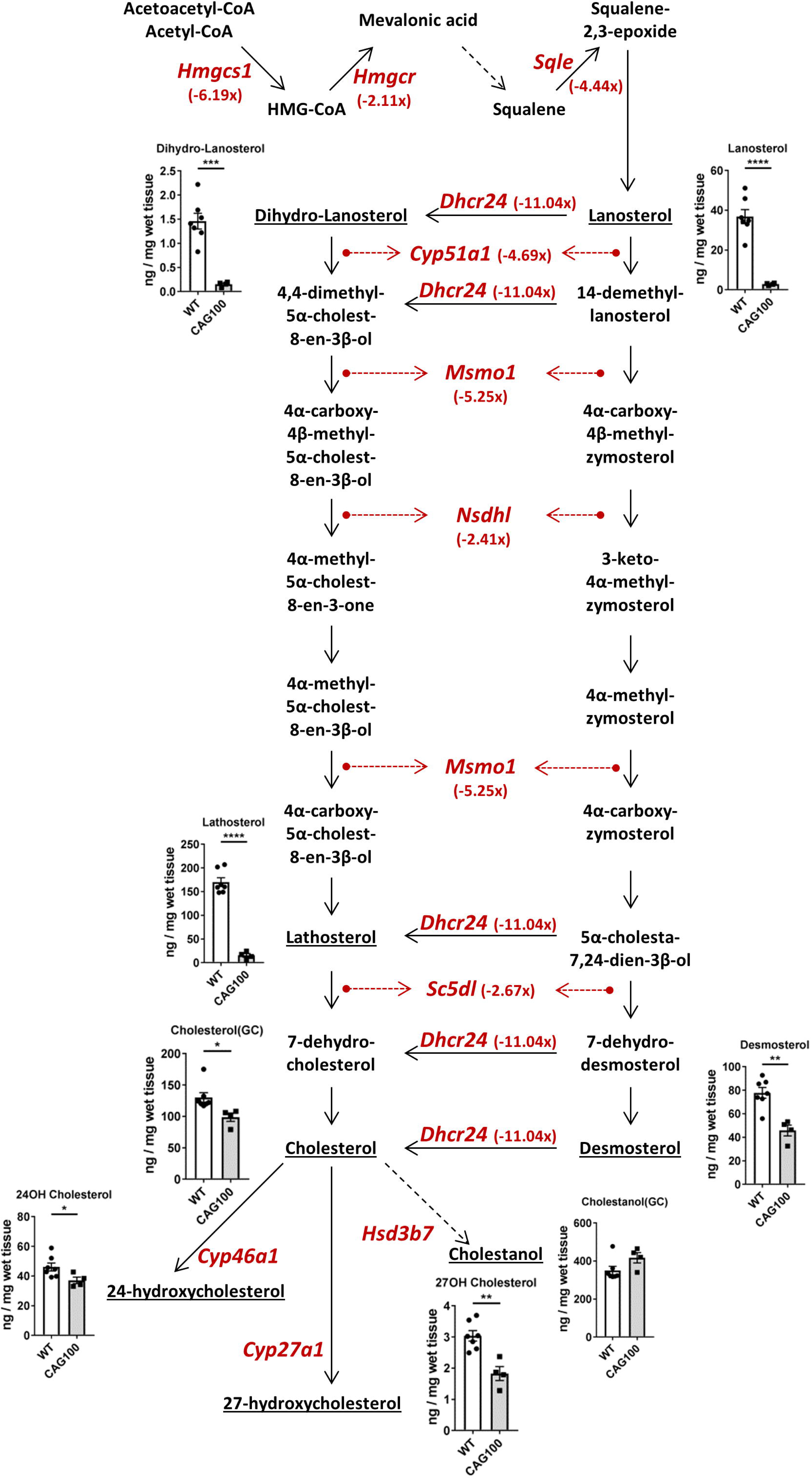
Cholesterol biosynthesis pathways show suppressed expression of enzymes and deficient metabolism intermediates in KIN spinal cord. In this schematic presentation of cholesterol biosynthesis via lathosterol and via desmosterol, significant transcriptional downregulations of various enzymes are highlighted in red letters for each gene symbol and each fold-change. The quantification of several intermediate metabolites is illustrated by bar graphs. The gas chromatography analysis was performed in 12-month-old spinal cord from 7 WT versus 4 KIN mice, see Suppl. Table S5.

### Progressive downregulation of further factors that mirror specific neuron pathways

Several calcium-binding factors with selective expression in different neuron populations seemed relevant in Table 1 and were highlighted in beige color: The progressive decrease of *Calb2* mRNA encoding Calretinin in cerebellum mirrors specifically the GLUergic parallel fibers, whereas the depletion of Parvalbumin (encoded by the *Parva* gene) is a marker of GABAergic Purkinje neurons. Both are lost from ALS spinal cord tissue [26, 71, 117, 228]. The parallel reduction of *Clgn* mRNA encoding the endoplasmic reticulum chaperone Calmegin represents cerebellar deep neurons and spinal motor neurons, according to the Allen mouse spinal cord *in-situ* hybridization data. These observations suggest that the degeneration affects several neural projections in parallel rather than in a hierarchical time-course.

The strongest progression marker in Table 1 was the decreased expression of the *Klk6* gene, which contains a purine repeat similar to the Friedreich ataxia gene and shows dysregulated expression during the neurodegenerative process of Alzheimer’s disease [190]. Similarly, the dysregulation of *Ina* expression and of *Il33* in Table 1 for the CAG100-KIN were similarly reported for ALS [70, 106]. The progressive decrease of *Unc13c* mRNA levels in Table 1 reflects the specific motor deficits of these mice, in view of its key role in cerebellar parallel fibers for fast reflexes and motor learning [9, 128]; the depletion of *Slc6a11* encoding the GABA-transporter-3, and of *Gabrb2* encoding the GABA-A-receptor-beta-2, in CAG100-KIN clearly mirrors the dysfunction of GABA-ergic signaling in the spinal dorsal horn and in cerebellar Purkinje cells [63, 86]; the deficiency of *Itih3*-encoded protein complexes was documented in Table 1 and occurs similarly after denervation, leading to diminished exploration and anxiety-like behavior in mice [21, 61, 134].

### Gradual expression anomalies for neurodegenerative disease genes Cdr1, Kcna2, Kif5a and Ano3 are prominent

As novel insights into the progressive spinal cord pathology in this SCA2 model, Table 1 also highlights progressively reduced expression for several known neurodegeneration genes: firstly, *Cdr1* (encoding Cerebellar Degeneration Related Protein 1, aka CDR62 or CDR34 or Yo-antigen) downregulation relates to its well-known autoimmune depletion as a cause of paraneoplastic ataxia [18] - it is interesting to note that *Cdr1* expression is induced by the myelination factor Prion protein [172]; secondly, the Spinocerebellar Ataxia gene and voltage-gated potassium channel *Kcna2*, which is preferentially expressed in afferent synapses onto the degenerating neurons [73, 229]; it is interesting to note that the parallel reduction of *Kctd3* and *Kctd9* affects two factors with potassium channel tetramerization domains; thirdly, the ALS disease gene *Kif5a* [20, 130] and its interactors *Kif5b* and *Kif5c*, which encode factors of axonal transport; *Kif5a* clusters with the progressive dysregulations of *Uhmk1* (aka Ser/Thr-Protein kinase KIS) and *Ina* (aka internexin neuronal intermediate filament) in Table 1, since these factors relate to ribonucleoprotein and stress granule transport [24, 52, 108]. In the same context, the progressive expression downregulation of *Hecw1* seems relevant, since this ubiquitination enzyme is responsible for the degradation of the ALS disease protein SOD1, is sequestrated into the cytosolic aggregates in ALS neurons, and its mutation leads to ALS-like phenotypes in mouse [123, 237]; fourth, the downregulation of *Ano3* is important in view of its impact on tremor and dystonia [30, 196]; similarly, the reduced expression of *Gnal* encoding the G-protein G(olf) alpha, and of *Rgs7bp* encoding R7bp as general regulator of G-protein signaling appears relevant, in view of *Gnal* mutations triggering dystonia type 25 and the key role of R7bp in spinal afferents [51, 104, 141]; fifth, the insidious reduction of *Scn4b* mRNA seems relevant, given that *Scn4b*-null mice show motor coordination and balance deficits [158], that *Scn4b* expression depends on GABA-A signaling [150] and that *Scn4b* depletion was also observed in the striatum affected by polyglutamine-neurotoxicity due to Huntington’s disease mutation [139]. In view of the importance of some of these factors, validation of their dysregulation expression was done by RT-qPCR (Suppl. Fig. S9).

### Insidious downregulation of Eif5a2 and Unc80 as factors within the interactome of ATXN2

It was particularly interesting to observe the depletion of *Eif5a2* transcripts encoding a translation initiation factor, given that ATXN2 is associated to PABPC1 in the ribosomal translation complex [35, 46, 96, 208], but its exact role was never understood. The EIF5A protein is the only factor containing the unusual conserved amino acid hypusine, was reported to aid translation of polyproline-motifs, modulates co-translational ER translocation, promotes stress granule assembly and mRNA decapping, is key for pancreatic beta-cell inflammation in diabetes mellitus, and mediates the effect of polyamines on neuronal process extension as well as survival via control of autophagy-controlling factors TFEB and ATG3. EIF5A also regulates mitochondrial respiration via initiation from alternative start codons. In addition, EIF5A restricts RNA virus infections [22, 55, 64, 78, 101, 110, 112, 113, 125, 148, 152, 164, 235]. It is important to note in this context that ATXN2 is cleaved by Coxsackie-virus and Polio-virus, during their optimized efforts to diminish host cell defenses against invading viral RNA [84]. Therefore, we used RT-qPCR to systematically assess expression dysregulation triggered by ATXN2 mutations in cerebellum as an independent validation effort, examining various factors that are involved in the RNA translation and shuttling, versus degradation of toxic viral RNA (Suppl. Table S5). Indeed, this approach confirmed that independent from polyQ expansions, altered ATXN2 functions lead to significant expression downregulations for *Eif5a2*. A similar downregulation was observed for the RNA decapping enzyme *Dcp2* that co-localizes with the PABPC1 interactor TOB1 in P-bodies, but unlike ATXN2 and PABPC1 is not sequestrated to viral production factories around lipid droplets [4, 188]. The concept of excessive exposure to toxic RNA was further substantiated, when RT-qPCR experiments validated microarray data in 14-month-old *Atxn2*-CAG100-KIN spinal cord on the 1.6-fold transcriptional upregulation of *Rnaset2* (Suppl. Fig. S9). This ribonuclease with localization in lysosomes is responsible for the degradation of mitochondrial RNA and mitochondria-associated ribosomal RNA [69, 77]. Mutations in *Rnaset2* cause neuroinflammation in a syndrome with cystic leukoencephalopathy [75]. The age-progressive release of toxic RNA and DNA from dysfunctional mitochondria was recently shown to constitute a key stress for innate immune defenses and for the neuroinflammation underlying Parkinsonian brain atrophy [38, 193, 226].

Another factor at a physiological site of ATXN2 localization that showed steady expression downregulation during the neurodegenerative process was UNC80, which is activated by the tachykinin receptor 1 (TACR1) via SRC as an interactor of ATXN2 [41, 109, 133]. TACR1 binds substance-P selectively for the sensory perception of itch, pain and inflammation. It is abundant in the spinal cord dorsal horn neurons but is also a component of microglia sensing [28, 241]. Tachykinin levels in the cerebrospinal fluid of ALS patients have been found elevated [119]. Furthermore, TACR1 accumulation was documented in the SOD1 mouse model of ALS and its pharmacological inhibition was found to be neuroprotective [23, 102, 192]. In the *Atxn2*-CAG100-KIN spinal cord, a significant decrease of *Tacr1* mRNA levels was evident already at the exceptionally early age of 3 months, and remained similarly strong until the age of 14 months (Suppl. Fig. S9). This observation provides a possible molecular correlate for our phenotypic finding that sensory pathology precedes motor pathology in SCA2. Furthermore, the strongest upregulation among all transcripts (∼45-fold upon RT-qPCR, Suppl. Figure S9) concerned the Glycoprotein NMB (aka osteoactivin or Hematopoietic Growth Factor Inducible Neurokinin-1 = HGFIN), encoded by *Gpnmb*. This neural-expressed glycoprotein interacts with substance-P, it activates SRC signaling, its depletion reduces neuropathic pain, and it can be induced by lysosomal stress also in microglia [53, 76, 114, 157]. It is highly relevant to note that GPNMB has a neuroprotective role for TDP43 toxicity or ALS [127, 199]. GPNMB was also described as neurodegeneration biomarker in Alzheimer’s, Parkinson’s, Gaucher’s disease and PLOSL leukodystrophy [79, 82, 124, 126, 173].

## Discussion

We have previously generated the novel *Atxn2*-CAG100-KnockIn mouse and shown for the cerebellum that (i) the temporal evolution of locomotor deficits and progressive atrophy, (ii) the spatial distribution of its pathology, and (iii) the neurochemical anomalies upon brain imaging faithfully reflect the known features of SCA2 [185]. For the spinal cord pathology as well, the current study confirms that this mouse mutant represents an authentic SCA2 model: Neurophysiologically, the distinctive features of SCA2 patients around clinical manifestation include an early sensory neuropathy by predominantly axonal lesion with signs of myelin damage [16, 146, 166, 168, 169, 211, 212], before the degeneration of lower and upper motor neurons starts with cranio-cervical preference [214-216, 219, 220]. Similarly, the *Atxn2*-CAG100-KIN showed sensory neuropathy as earliest peripheral manifestation of disease at the age of 9-10 months. As a pathological hallmark of spinal motor neuron affection, the progressive aggregation of TDP43, ATXN2 and other stress granule proteins was observed in the nervous tissue of human SCA2 [44, 89, 167, 183], and is also now documented in this mouse model. In view of the rarity of SCA2 autopsies, no human expression profiles became available until now that could elucidate the pathological mechanisms in the affected tissue. Thus, the *Atxn2*-CAG100-KIN mouse provided a unique opportunity to understand the spinal pathology of SCA2, and the molecular insights documented here are completely novel.

Overall, the pathway enrichments demonstrated a decreased expression for synaptic and axonal factors as well as cholesterol enzymes. This loss occurs in parallel to astrogliosis, as expected. But interestingly, our data represent the first report that enhanced expression of lysosomal factors and microgliosis are prominent at late disease stages. Mechanistically, the activated microglia may be partially due to the presence of ATXN2 aggregates, which are presumably extruded from neurons and internalized by phagocytosis, but this may also be exacerbated by the robust expression of expanded ATXN2 in stressed microglia cells, as a cell-autonomous trigger.

From the expression profiles in the KIN spinal cord, two molecular cascades can be assembled as plausible scenarios, both culminating in microgliosis. Firstly, as a putative correlate of the sensory neuropathy the transcript expression of several key factors in afferent signaling changed. The downregulation of the itch/pain-related Substance-P receptor *Tacr1* reached significance particularly early at the age of 3 months. The *Tacr1*-dependent neuronal excitability factor *Unc80*, which encodes a scaffold for the ATXN2-interactor SRC, was identified as one of the best biomarkers for disease progression. The afferent excitability problem seems to involve altered potassium homeostasis, in view of the conspicuous expression dysregulation of several presynaptic K^+^ channels (Ataxia disease genes *Kcnj10*, *Kcnc3*, *Kcna1* in Suppl. Fig. S7C; *Kcna2* with K^+^-channel tetramerization factors *Kctd9* and *Kctd3* in Table 1). This observation is in good agreement with a previous report that impaired excitability of Purkinje neurons in a SCA2 mouse model was rescued by modulation of K^+^ dependent hyperpolarization [43]. Also the expression dysregulation of *Ano3* and *Gnal*, two disease genes responsible for dystonia and tremor, may be caused within the same sensory pathway. In this context, it is important to consider that SCA2 mutations always trigger intention tremor and may have a clinical presentation mainly with tremor or dystonia [32, 50, 92, 116]. At late stage, the strongest upregulation among all transcripts concerned the Substance-P interactor *Gpnmb*, which can be strongly induced in microglia cells by lysosomal stress and exerts a neuroprotective role in TDP43 pathology.

Secondly, a molecular cascade was documented as correlate of motor neuron affection by RNA toxicity, in excellent overlap with the pathogenesis of ALS. The downregulation of *Eif5a2* may be a primary consequence of ATXN2 expansion, given that both factors colocalize in the ribosomal translation complex and that both act in the defense against viral RNAs. Also the accumulation of the toxic DNA/RNA sensor and stress granule assembly factor PQBP1 [91] may be a direct consequence of its sequestration by polyQ-expanded ATXN2. The downregulation of *Dcp2* as RNA degrading enzyme in the P-body may reflect the pathological retention of RNAs in stress granules affected by ATXN2 aggregation. In contrast, the upregulation of *Rnaset2* is probably a compensatory effort, since this enzyme has the ability to degrade inappropriately methylated RNAs, which are released from dysfunctional mitochondria in ever higher quantities during the ageing process. Altogether, the accumulation of toxic RNAs seems to be reflected by the transcriptional induction of sensors such as *Trl7*, of the signaling factor *Ripk1*, the particularly early elevation of lysosomal PGRN protein and the increase of cytosolic CASP3 protein levels, which promote the aggregation of TDP43 at stress granules. A known correlate of this cascade in SCA2 pathology is probably the accumulation of Staufen1 as a marker of neuronal RNA granules, which is also recruited into ATXN2 aggregates [144]. Given that KIF5 is crucial for anterograde transport of virus particles [42], the progressive expression downregulation of all three KIF5 subunits in the KIN spinal cord might constitute a protective antiviral response within neurons. The progressive downregulation of *Hecw1* expression is known to modulate the vulnerability of motor neurons by reduced degradation of misfolded SOD1, which was shown to affect stress granule dynamics via interaction with G3BP1 [54]. It is important to note that the permanent cytosolic sequestration of such nucleotide processing factors by ATXN2 aggregates would lead not only to impaired RNA quality control, but also to altered defenses of motor neurons e.g. against poliovirus RNA infections [40, 115, 202, 227]. Of course, the hierarchical sequence of events in both scenarios may be more complex, but these dysregulations clearly stood out in the global transcriptome profiles of presymptomatic and preterminal KIN spinal cord.

The comparison of both stages identified 54 genes with progressively changed expression during the disease course, including conspicuously strong downregulations for several enzymes of cholesterol biogenesis. It is crucial to note that also in a mouse model of SCA3 (Machado-Joseph disease), recent global transcriptome profiling demonstrated a significant enrichment of cholesterol biogenesis dysregulations in brainstem, but increased levels of ceramides, di- and triglycerides in blood [204]. Furthermore, for SCA3 it was shown that restoration of brain cholesterol homeostasis has therapeutic potential [131]. Our observation of reduced cholesterol biogenesis in the *Atxn2*-CAG100-KIn mouse is easily related with the documented depletion of peripheral fat tissues in SCA2 [121], and with findings of early demyelination in this CAG100-KIN mouse and in SCA2 patients [57, 140, 168]. The quantitative analysis of several intermediate steps of cholesterol metabolism in the KIN spinal cord confirmed a significant deficit of cholesterol and massive deficiencies for several precursors. Even stronger depletion of cholesterol and also of very-long-chain sphingomyelins (products of the ataxia disease gene *Elovl4*, also downregulated in the CAG100-KIN, see Suppl. Figure S7C) were recently documented in a SCA2 patient cerebellum [184]. Cholesterol is a requirement for the synthesis of sex hormones and of corticosteroid stress hormones, so this might explain also the gradually declining expression of androgen-dependent factor *Klk6* as the most dramatic downregulation effect. The progressive loss of *Cdr1* expression might be explained in a similar context, since it is known to be regulated by the myelination factor *Prnp* (prion protein). Our study is not the first report of a connection between RNA-binding proteins such as ATXN2, TDP43 or TIA1 on the one hand and obesity or cholesterol on the other hand. Transgenic overexpression or KnockIn of TDP43 in mouse triggered weight loss and increased fat deposition with elevated HDL cholesterol in blood [195, 197]; LXRbeta^-/-^ mice showed TDP43 aggregation together with higher brain cholesterol [87]; conversely, the KnockOut of TDP43 results in dramatic loss of body fat [33]; similarly, the KnockOut of TIA1 triggers upregulation of fat storage factors [72] and the KnockOut of ATXN2 triggers hypercholesterolemia [95]. In ALS, the spinal cord ventral horn of patients showed a significant decrease of cholesterol [66], whereas in blood an increase of cholesterol was documented [60]. Controversial reports exist whether the use of cholesterol-lowering medication such as statins enhances or reduces ALS risk [49, 59]. In SCA2, the lipid metabolism seems to be dysregulated at a more basic level than cholesterol generation, given that expression of *Nat8l* as the enzyme responsible for N-acetylaspartate (NAA) synthesis was gradually declining in *Atxn2*-CAG100-KIN spinal cord. Indeed, spinal NAA decreases during the disease course of ALS patients [27, 80, 171].

Overall, the gradual expression reduction of several cholesterol enzymes is prominent and strong, so it may become useful for the future validation of progression biomarkers in SCA2 patient samples. However, cholesterol depletion is hardly a SCA2-specific feature. It is therefore important to state that the progressive expression reduction of *Unc80* and *Eif5a2* concerns factors within the ATXN2 interactome and might thus represent primary and specific effects of SCA2 pathology. At present, clinical trials in SCA2 depend on the quantification of clinical, neurophysiological and imaging features with documented progression, such as the clinical SARA score [39, 83], quantification of sensory neuropathy [213], saccade slowing [162, 217, 218], periodic leg movements during sleep [161, 221] and brain volumetry [1, 159]. Since any improvement in these disease features occurs over extended periods of time, there is an unmet need to characterize molecular biomarkers that mirror therapeutic benefits very rapidly.

In conclusion, the spinal cord of our new *Atxn2*-CAG100-KIN mouse mutant revealed first insights into the molecular pathogenesis of SCA2. Sensory and motor affection involves prominently a loss of axonal and presynaptic factors, in parallel to depletion of cholesterol and phospholipid enzymes. Various molecular pathways are irritated by the sequestration of ATXN2 interactor molecules into aggregates, principally the role of stress granules in the defense against toxic RNAs. Several other altered pathways such as ribosomal translation, calcium homeostasis and cholesterol biogenesis occur at the endoplasmic reticulum, where ATXN2 was shown to play a crucial role for structure and dynamics according to studies in *C. elegans* and *D. melanogaster* [36]. SCA2 pathogenesis has considerable overlap with the mechanisms documented in other ataxias, dystonias, and with ALS-associated features such as RNA toxicity and endoplasmic reticulum dysfunction [200].

## Acknowledgements

For their technical assistance, we are grateful to Birgitt Meseck-Selchow and Gabriele Köpf in Frankfurt, to Jérome Sinniger in Strasbourg, and to the staff of the ZFE animal facility at the Goethe University in Frankfurt. For financial support, we thank the Deutsche Forschungsgemeinschaft (grants AU96/11-1 and 11-3 to GA).

---

**Supplementary Table S1: 10-week-old KIN spinal cord global transcriptome dysregulations >1.2-fold, assessed for pathway enrichments by STRING statistics**. Different datasheets are provided to document enrichment scores and individual factors involved, among GO terms Biological Process, Molecular Function, Cellular Component, PubMedID of Reference Publications, KEGG pathways, Reactome pathways, UniProt keywords, PFAM protein domains, InterPro protein domains and features, and SMART domains. Colors are used to highlight pathways that are prominent in STRING diagrams of Suppl. Figure S5A-B.

**Supplementary Table S2: 14-month-old KIN spinal cord global transcriptome dysregulations beyond 2-fold, assessed for pathway enrichments by STRING statistics**. Different datasheets are provided to document enrichment scores and individual factors involved, among GO terms Biological Process, Molecular Function, Cellular Component, PubMedID of Reference Publications, KEGG pathways, Reactome pathways, UniProt keywords, PFAM protein domains, InterPro protein domains and features, and SMART domains. Colors are used to highlight pathways that are prominent in STRING diagrams of Suppl. Figure S5C-D.

**Supplementary Table S3: KIN spinal cord global transcriptome dysregulations that doubled fold-change over lifespan, assessed for pathway enrichments by STRING statistics**. Different datasheets are provided to document enrichment scores and individual factors involved, among GO terms Biological Process, Molecular Function, Cellular Component, PubMedID of Reference Publications, KEGG pathways, Reactome pathways, UniProt keywords, PFAM protein domains, InterPro protein domains and features, and SMART domains. Colors are used to highlight pathways that are prominent in STRING diagrams of Suppl. Figure S5E.

**Supplementary Table S4: Gas chromatographic quantification of cholesterol biosynthesis intermediate metabolites in 12-month-old *Atxn2*-CAG100-KIN spinal cord.** The datasheet shows genotype, animal identity numbers, gender, age, dry sample weight, absolute quantities of various precursor metabolites (black digits), cholesterol (blue digits) and derivative oxysterols (red digits), as well as ratios that were derived either from normalization versus mg dry weight, or versus µg cholesterol (blue field colors), or between two metabolites (grey fields). Average values (AVG) are shown below, together with significance p-values calculated by Student’s t-test.

**Supplementary Table S5: RT-qPCR survey of expression changes among RNA-binding factors upon ATXN2 mutations.** Diverse RNA-binding factors with roles in viral transcript repression, nuclear transcription and splicing, export and cytosolic trafficking, RNA quality control and decay were tested both in *Atxn2*-KO and CAG100-KIN cerebellum at advanced age. The table details the assays employed, gene symbols, nominal significance level for Student’s t-test, number of animals, mean and standard error of the mean values. Red cell background highlights significant upregulations, blue color illustrates downregulations, bold letters emphasize two factors with significant dysregulation in both mutants.

**Supplementary Figure S1: Functional assessment of hindlimb nerve function cannot demonstrate motor dysfunction at the age of 9-10 months. (A)** Neurophysiological assessment of 5 male and 1 female mutants with age-/sex-matched wildtype controls failed to detect motor neuropathy for sciatic nerve in lower limbs. Upon electromyography (EMG), no muscle denervation potentials were observed in hindlimb muscles. **(B)** RT-qPCR analyses of acetylcholine receptor isoforms in tibialis anterior and soleus muscle tissue as markers of denervation showed no major dysregulation in CAG100-KIN animals.

**Supplementary Figure S2: ATXN2 protein aggregates in spinal motor neurons sequestrate TIA1**. Triple co-immunofluorescent staining of ATXN2 (green) versus TIA1 (red) in *Atxn2*-CAG100-KIN mice (at 14 months of age) versus age-/sex-matched WT controls. Nuclei were detected by DAPI (blue color), scale bar reflects 25 µm.

**Supplementary Figure S3: Activation of immune pathways is observed in KIN spinal cord at the preterminal stage. (A)** RT-qPCR analyses of microglial Toll-like receptor isoforms *Tlr3*, *Tlr7* and *Tlr9*, together with complement factors *C1qa*, *C1qb*, *C1qc* and *C3* showed significant increases at the age of 14 months. **(B)** Quantitative immunoblot of polyglutamine-binding-protein 1 (PQBP1), as a sensor of toxic RNA/DNA and immune activator, showed increased protein levels at 14 months.

**Supplementary Figure S4: Activation of microglia in KIN spinal cord has pro-inflammatory toxic features.** In order to dissect the mode of microglial activation (whether it has anti- or pro-inflammatory nature), RT-qPCR analyses were performed to measure the transcript levels of *Trem2* and *Tyrobp* as markers of protective anti-inflammatory efforts, as well as *Irak4* and *Cybb* as markers of toxic pro-inflammatory activation. All transcripts were found upregulated at the preterminal stage of 14 months, with the highest induction of *Cybb* indicating a stronger pro-inflammatory state of microglia.

**Supplementary Figure S5: STRING bioinformatics plots of protein-protein-interaction clusters among KIN spinal cord transcriptome dysregulations**. For incipient disease stages at the age of 10 weeks, 1.2-fold dysregulations were analysed in the first two images. **(A)** Upregulations included *Pabpc1*, other translation initiation factors (e.g. *Eif1*), RNA helicases (e.g. *Ddx17*), spliceosomal factors (e.g. *Srsf3, Snrpc*), (manually placed below and right from the center image), RNA degradation enzymes (e.g. *Cnot6*), ribonucleoprotein components (e.g. *Tia1*, *Hnrnpa2b1*), ribosome subunits (e.g. *Rps2*), histone components (e.g. *Hist1h2af*), among many “nucleic acid binding factors” (highlighted as blue bullets). Upregulations also included factors with “acetylation” (yellow) and in the “inner mitochondrial membrane protein complex”. **(B)** Downregulations included two translation initiation factors (*Eif5a2, Eif2ak1*), several “protein tyrosine kinase” members (e.g. *Ntrk2*) with their downstream effectors such as MAP kinases (e.g. *Mapk9*) and CAM kinases (e.g. *Camkv*) (manually placed below and left from the center image, as light blue bullets). Downregulations also involved axonal (light green) and synaptic (dark green) factors, in particular many potassium channels (e.g. *Kcnj10*, *Kctd3*, in image center). Moreover, they concerned “cholesterol biosynthesis” enzymes (red bullets, e.g. *Sqle*) and “metabolism of lipids” (purple, e.g. *Acsl6*, *Stard1*). Given that too many factors showed 1.2-fold dysregulation, STRING bioinformatics at the preterminal stage was performed for 2-fold dysregulations. **(C)** Upregulations in spinal cord at 14 months included many “immune system process” components (blue bullets, particularly microglia factors, e.g. *Trem2* below center) and “lysosome” factors (yellow). **(D)** Downregulations at 14 months again concerned axonal (light green) and synaptic (dark green) factors, in particular many potassium channels (orange bullets, manually clustered in center). In addition, “sterol biosynthetic process” components (red color) were again affected, and the “metabolism of lipids” (purple). **(E)** As candidate markers of disease progression, any transcriptome dysregulation that progressed from 1.2-fold at 10 weeks to 2.4-fold at 14 months was assessed for significant pathway enrichments. The same colors as previously visualize the factors of “sterol biosynthetic process” (red), “lipid metabolism” (purple), “axon” (light green), “presynapse” (dark green), “potassium channel tetramerisation-type BTB domain” (orange). In addition, “neurotransmitter transport” (light blue), “endoplasmic reticulum subcompartment” (yellow), “plasma membrane” (dark violet), “cytoplasmic ribonucleoprotein granule” (brown) and “cell junction” (dark blue) were highlighted because of their enrichments. The significance levels and less prominent pathways are detailed in Suppl. Tables S1/2/3.

**Supplementary Figure S6: Pathway enrichment survey by Affymetrix Clariom D Transcriptome Analysis Console in 10-week-old KIN spinal cord.** Significant pathways are shown in predefined schemes, highlighting upregulations by red background and downregulation by green background, with color intensity reflecting the effect size. The pathways at incipient disease stage included “Calcium regulation in the cardiac cell” (significance 5.0), “Cytoplasmic ribosomal proteins” (0.1), “Cholesterol metabolism” (2.7), “Regulation of actin cytoskeleton” (7.7), “Mapk signaling” (4.2), “Insulin signaling” (4.2), “Focal adhesion-Pi3k-Akt-mTOR-signaling” (1.2), “Egfr1 signaling pathway” (2.0), “mRNA processing” (3.9), “Adipogenesis genes” (0.2), and “Spinal cord injury” (significance 0.1).

**Supplementary Figure S7: Pathway enrichment survey by Affymetrix Clariom D Transcriptome Analysis Console in 14-month-old KIN spinal cord.** Significant pathways are shown in predefined schemes, highlighting upregulations by red background and downregulation by green background, with color intensity reflecting the effect size. The pathways at preterminal stage included microglia (e.g. “Tyrobp causal network” with significance 10, “MicrogliaPathogenPhagocytosis” with significance 7.3) and inflammation (“B cell receptor signaling” with significance 9.3). Similar to incipient age, also “Calcium regulation in the cardiac cell” (significance 8.7), “Cytoplasmic ribosomal proteins” (6.1), “Cholesterol metabolism“ (5.3), “Regulation of actin cytoskeleton” (3.4), “Mapk signaling” (3.3), “Insulin signaling” (2.6), “Focal adhesion-Pi3k-Akt-mTOR-signaling” (2.6), “Egfr1 signaling” (1.9), mRNA processing (1.5), “Adipogenesis genes” (1.3), and “Spinal cord injury” (3.0) were detected.

**Supplementary Figure S8: Global transcriptome profile of 14-month-old KIN spinal cord presented as volcano plot, highlighting key factors of pathogenesis, (A)** concerning RNA toxicity, **(B)** concerning proteins responsible for ALS variants**, (C)** concerning proteins responsible for ataxia variants. The fold change of protein abundance is shown on the X-axis (filtering cut-off 1.2-fold), the negative logarithm of the nominal p-value is shown on the Y-axis (significance cut-off at 1.3 corresponds to p=0.05). Upregulated factors are shown in red, downregulations in blue color.

**Supplementary Figure S9: Additional validations of the high-troughput transcriptome data confirm the dysregulations in inflammation, sensory, motor and cholesterol pathways.** Quantitative RT-PCR analyses were performed to assess the expression levels of *Rnaset2* and *Gpnmb* in the RNA toxicity and inflammation pathway, *Unc80* and *Tacr1* within sensory neuropathy, *Ano3*, *Cdr1*, *Kif5a*, *Ttbk2*, *Kcna1*, *Kcna2* and *Scn4b* within dystonia-ataxia-ALS-tauopathy-axonopathy pathogenesis, *Cyp46a1* as a component of cholesterol turnover, and *Hmgcs1*, *Dhcr24*, *Msmo1* and *Cyp51a1* as enzymes for cholesterol biosynthesis.

