## Supplementary figures and images for "*Atxn2*-CAG100-KnockIn mouse spinal cord shows progressive TDP43 pathology associated with cholesterol biosynthesis suppression"

### Supplemental Figure S1

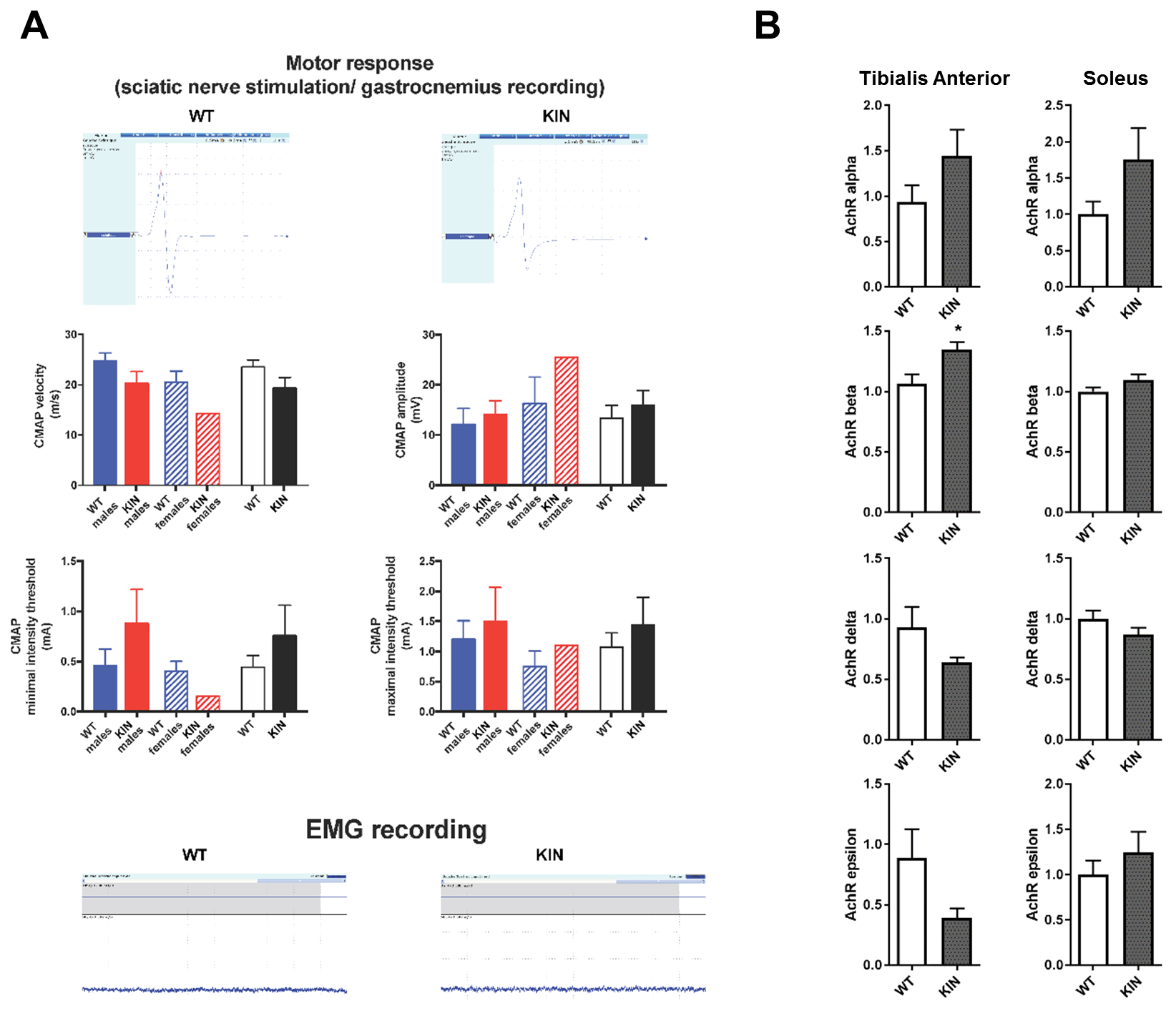

### Supplemental Figure S2

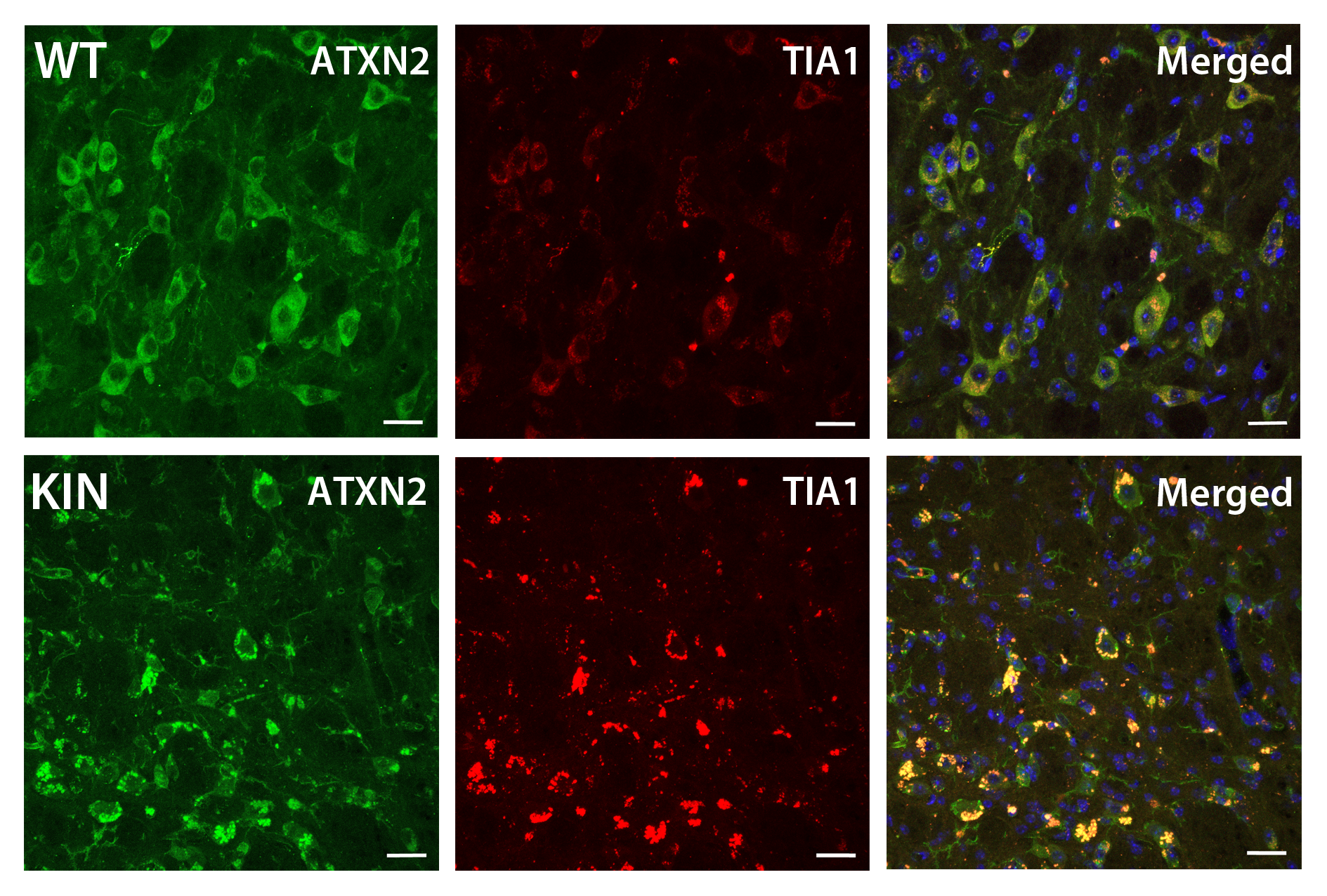

### Supplemental Figure S3

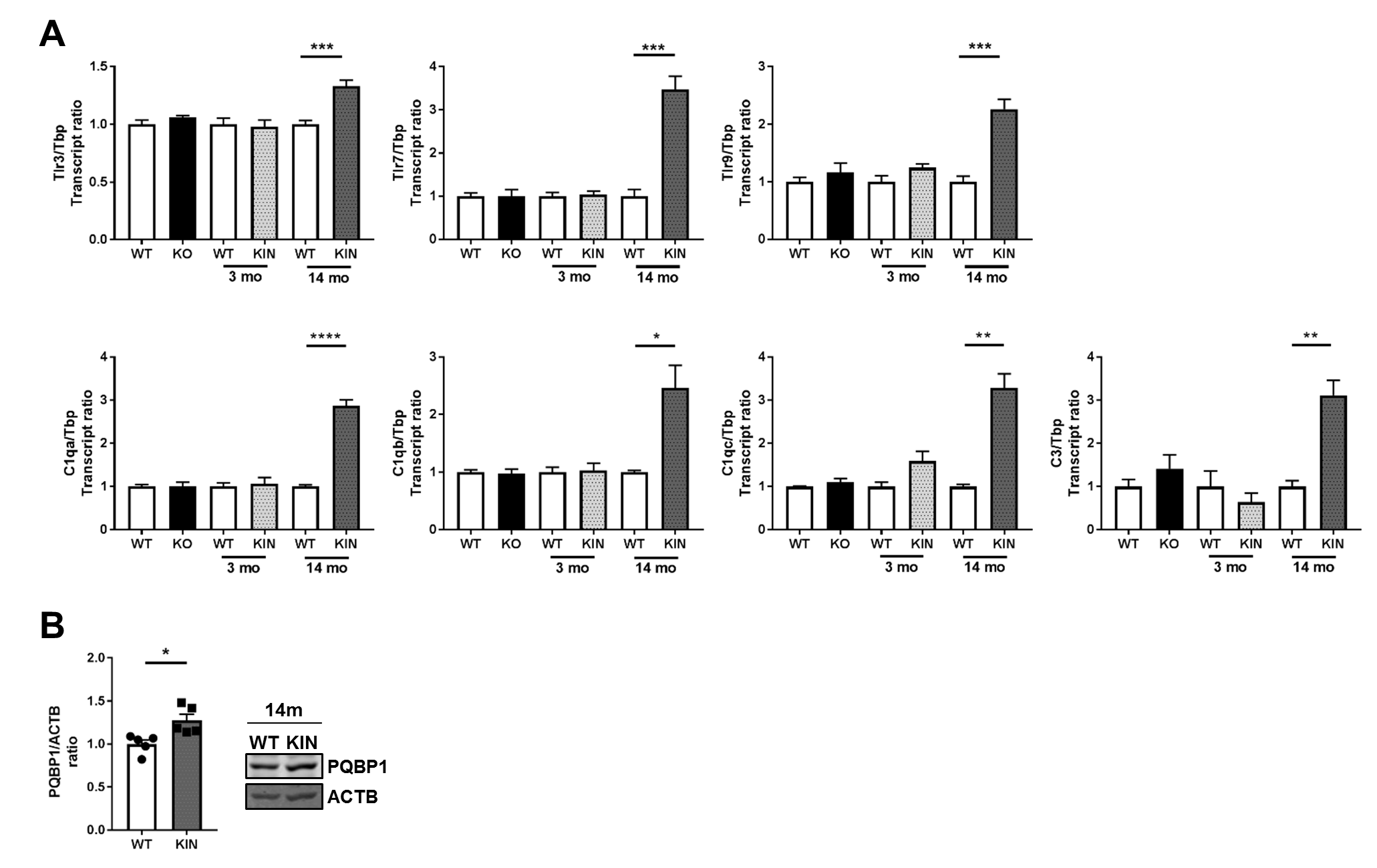

### Supplemental Figure S4

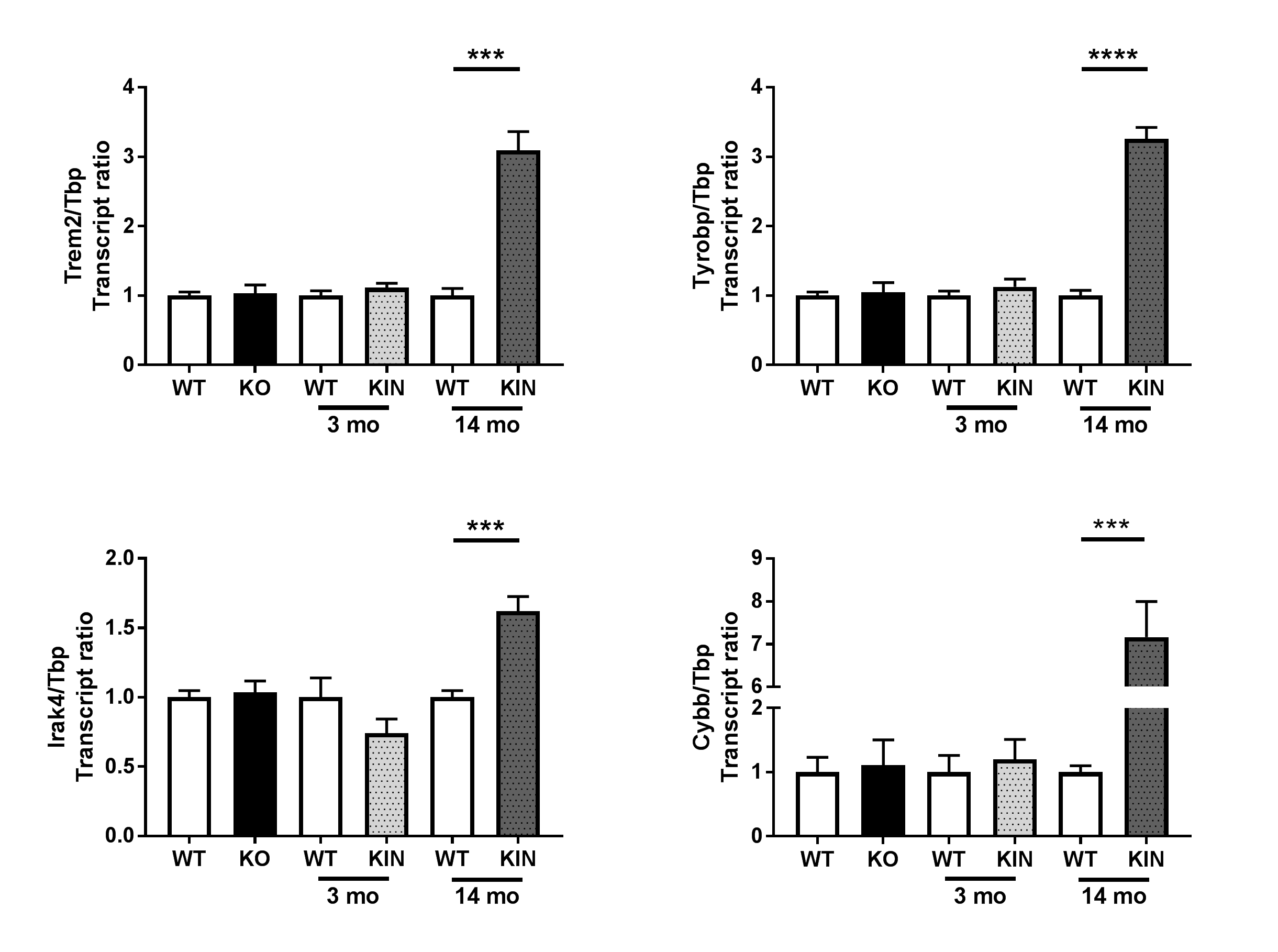

### Supplemental Figure S5

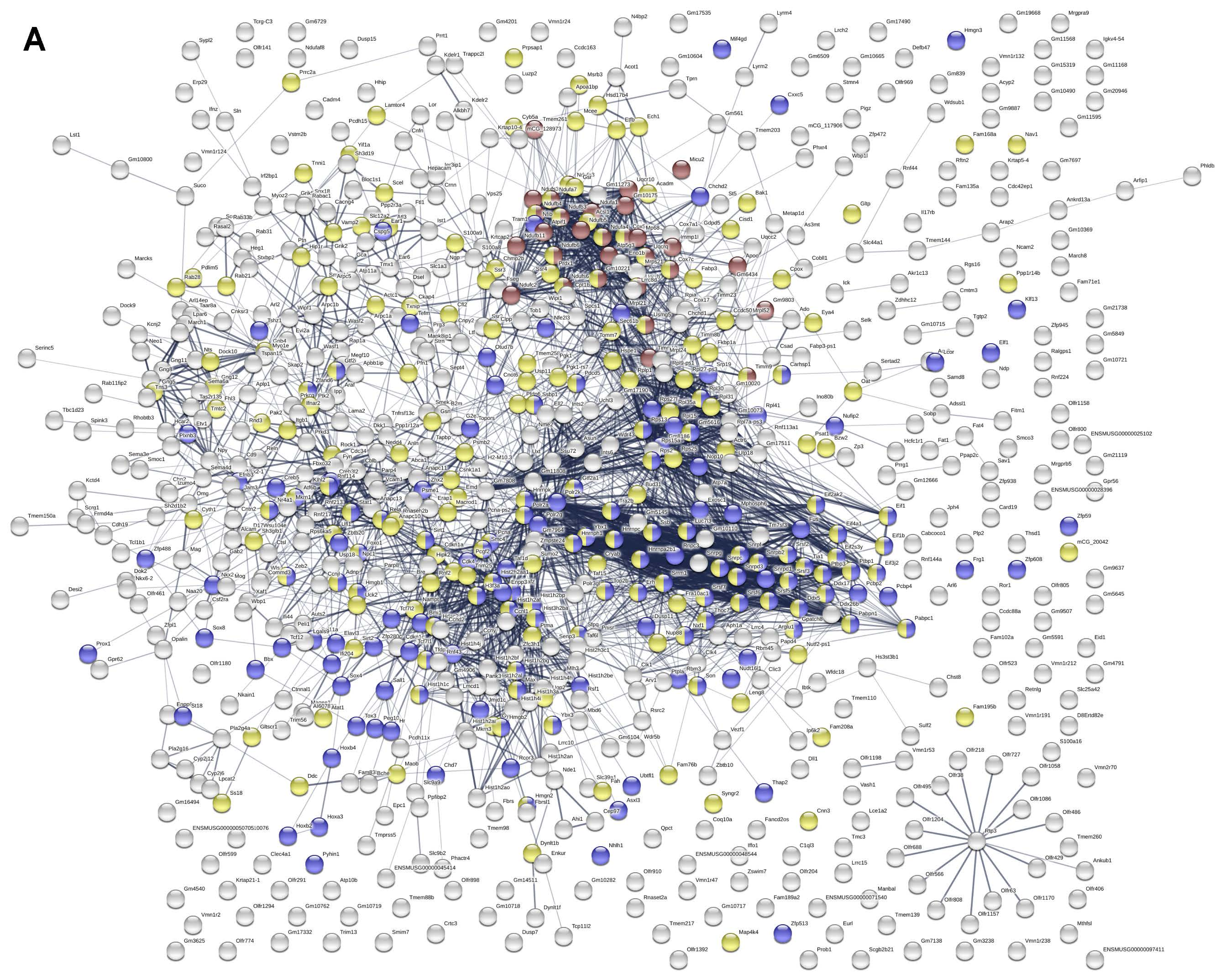

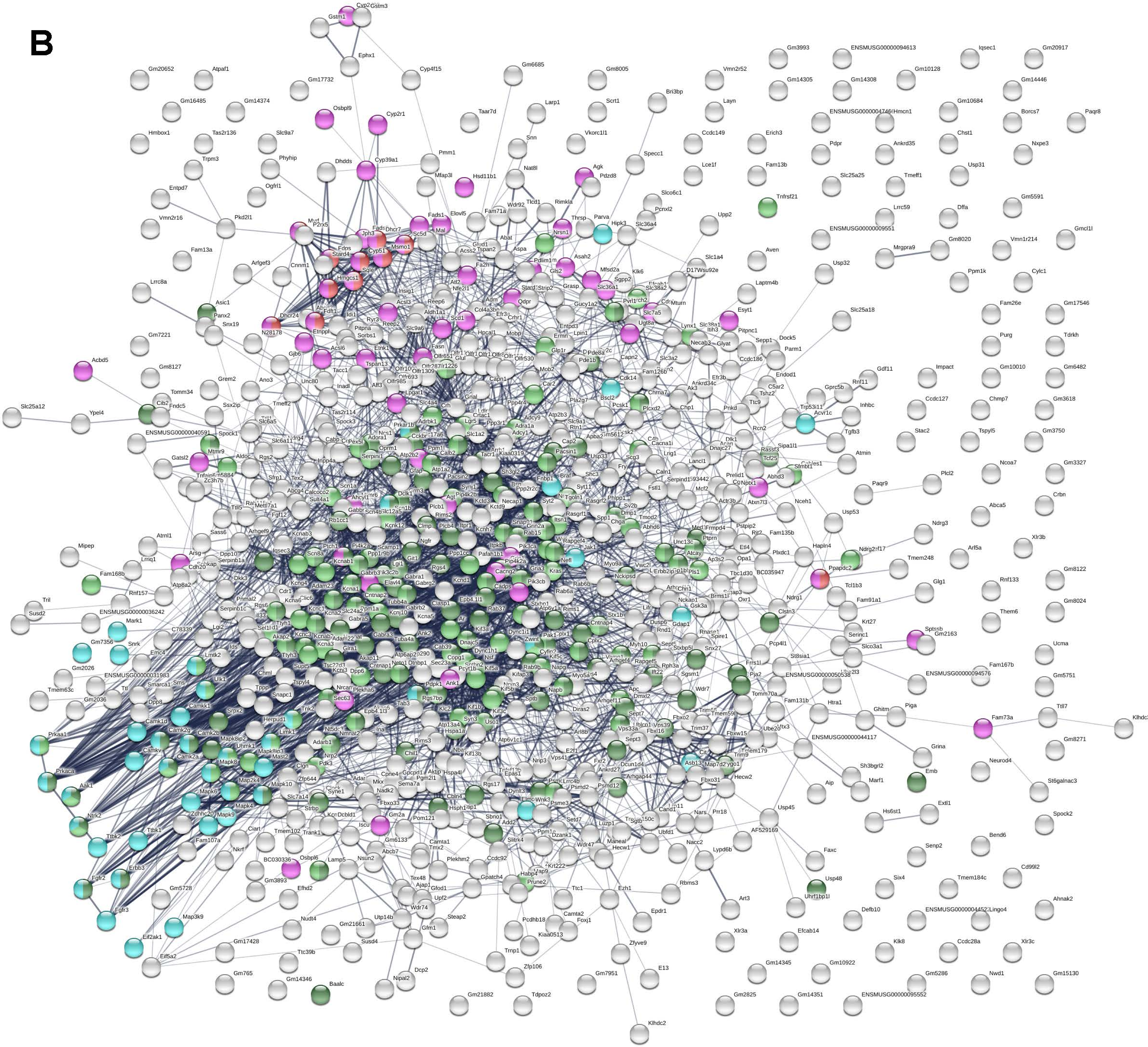

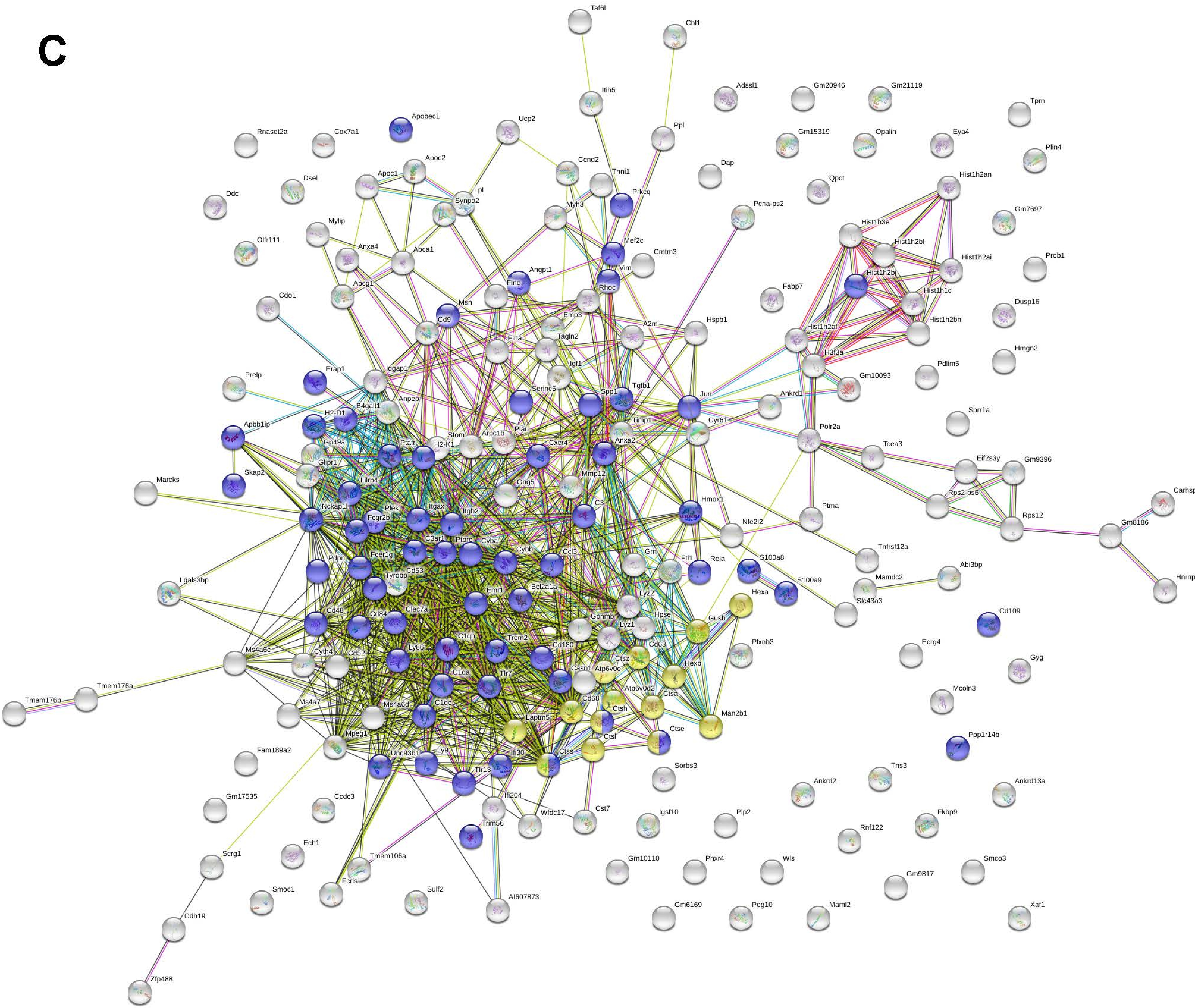

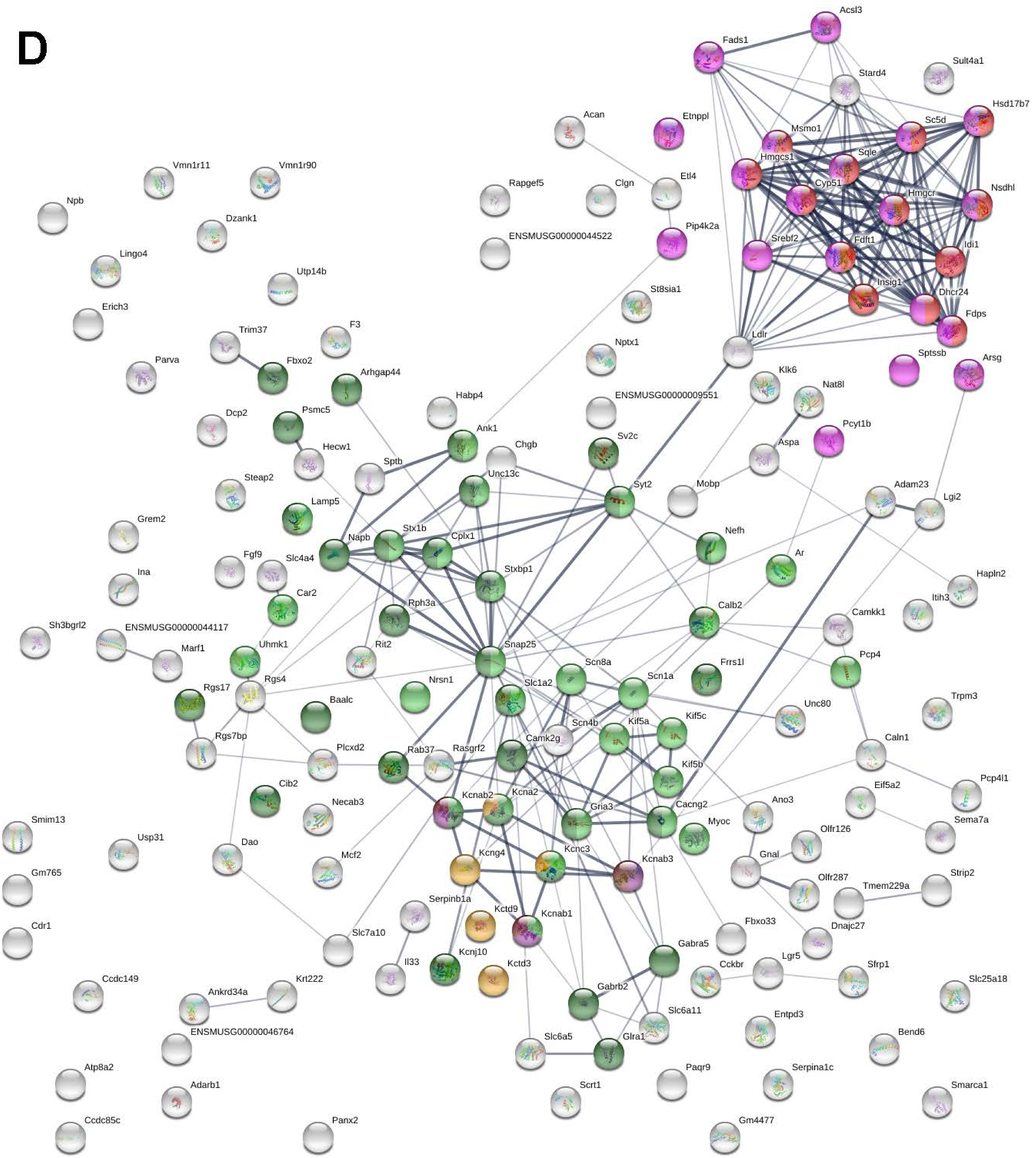

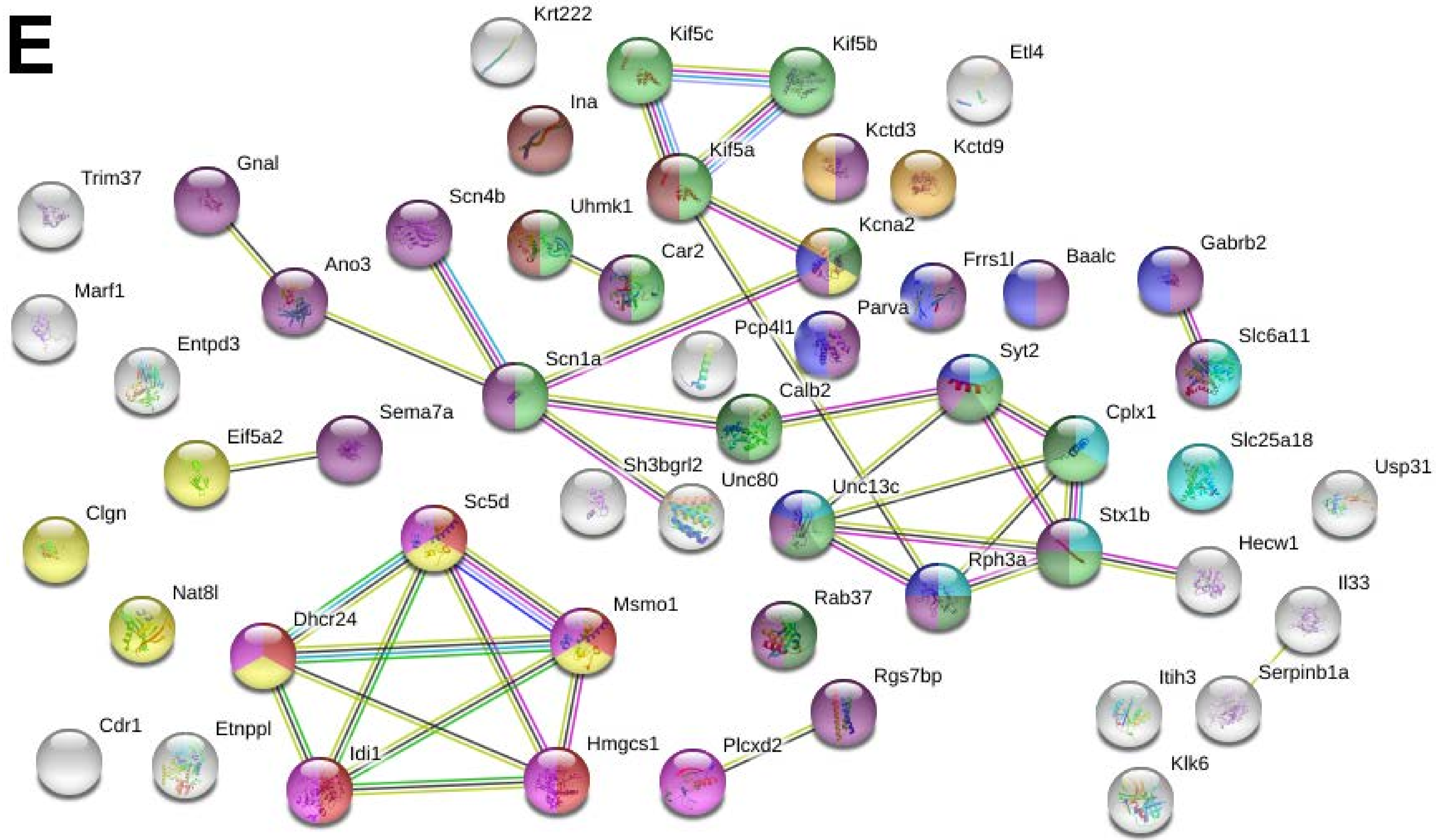

### Supplemental Figure S8

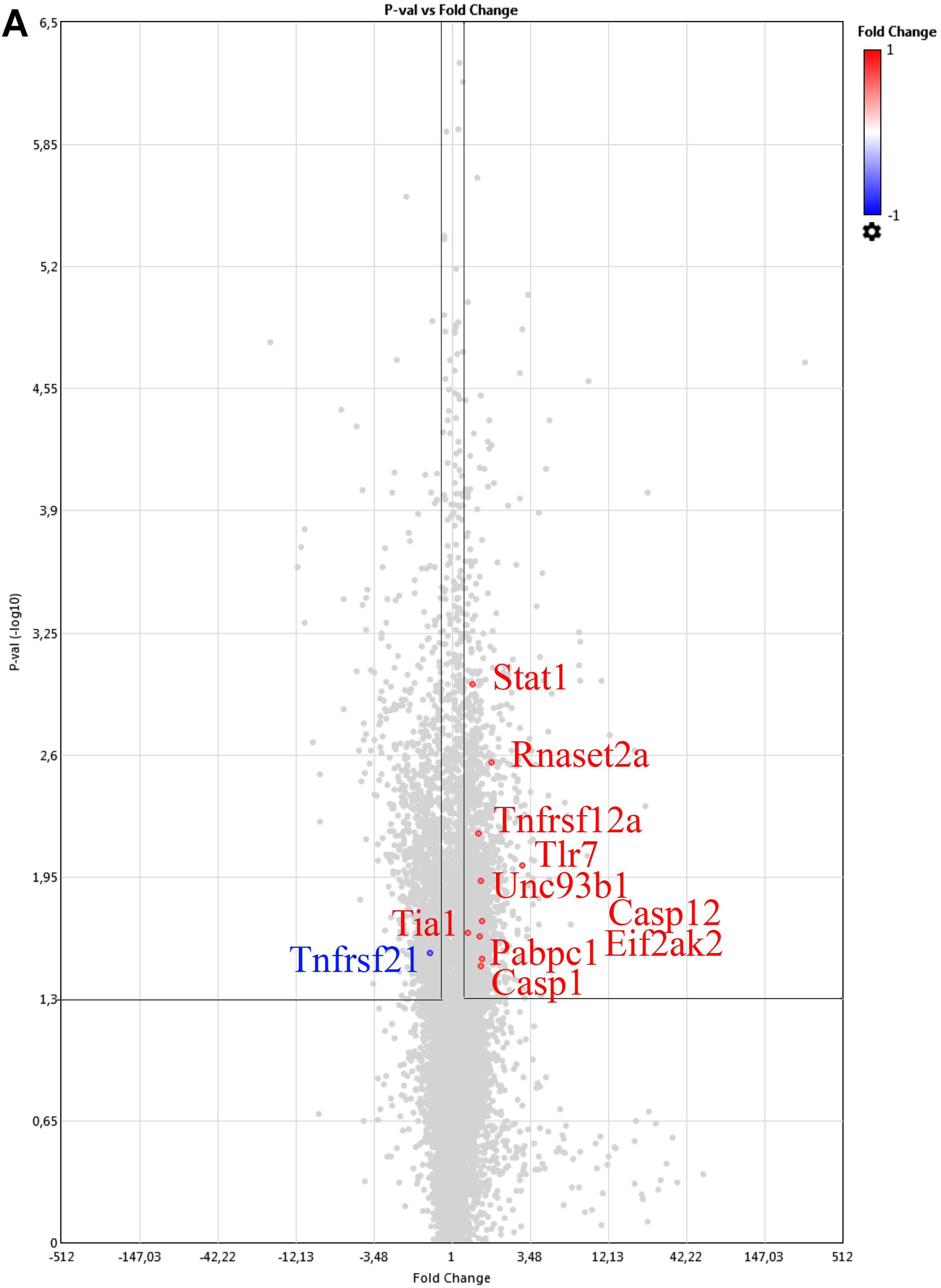

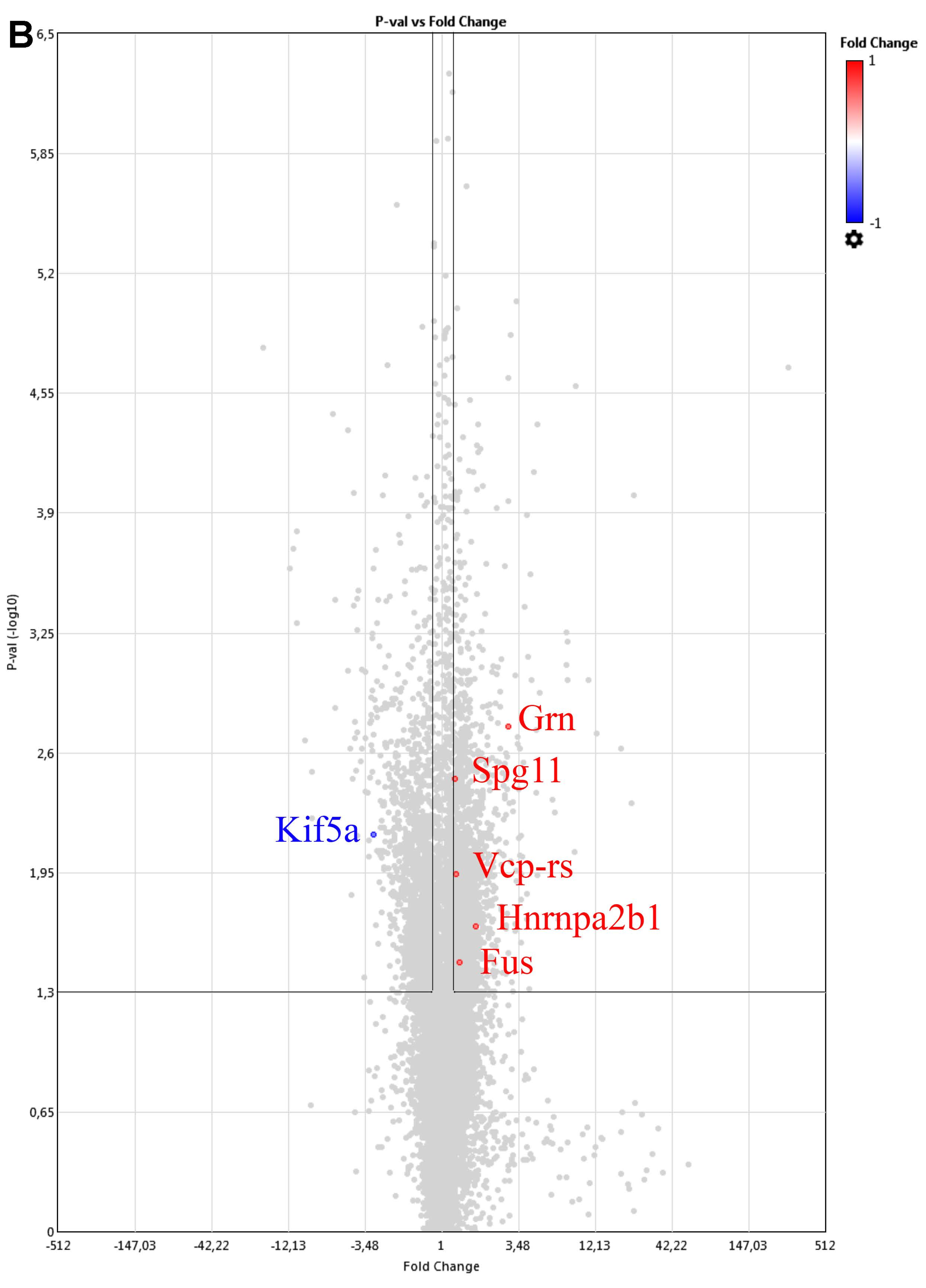

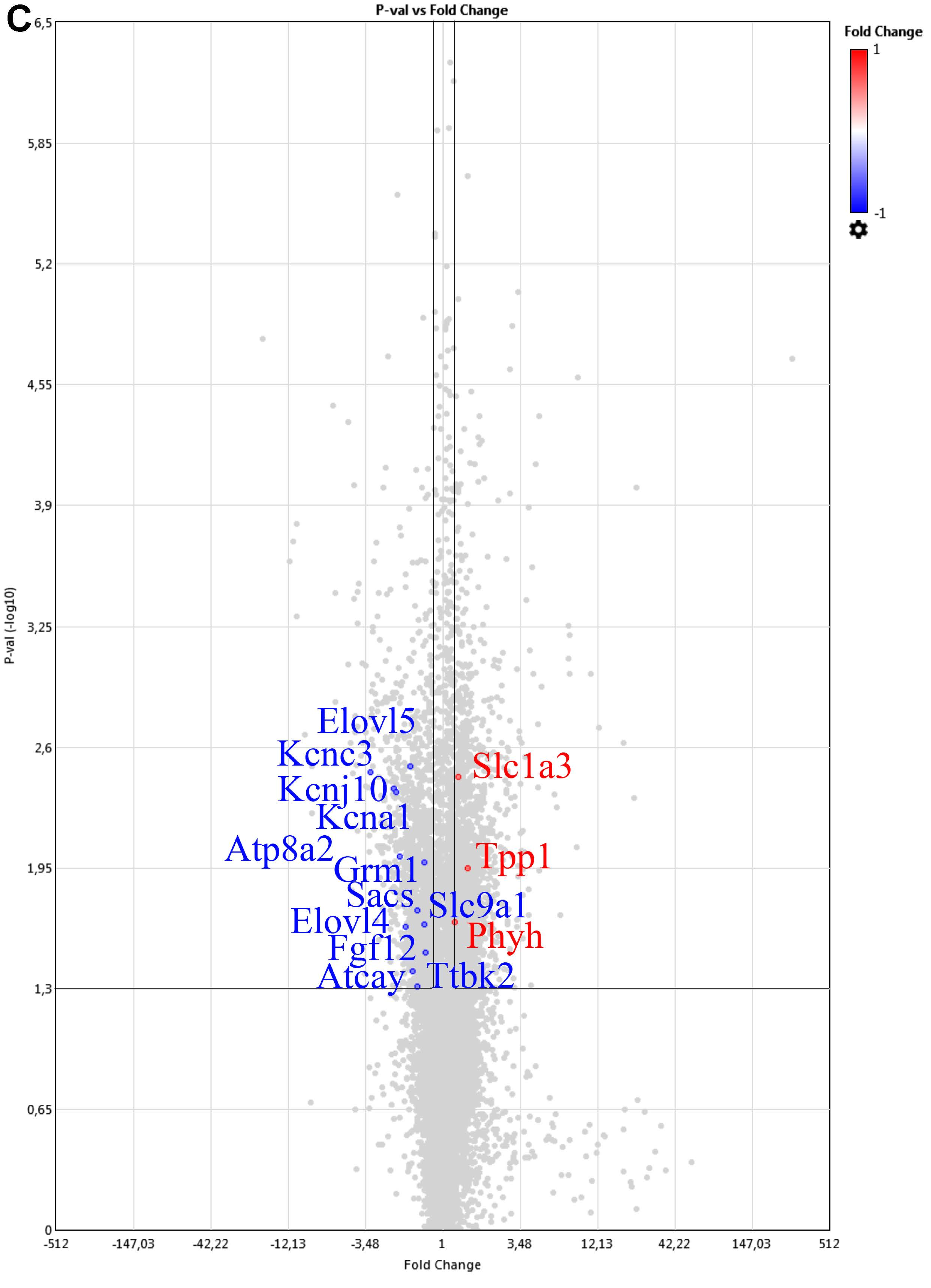

### Supplemental Figure S9

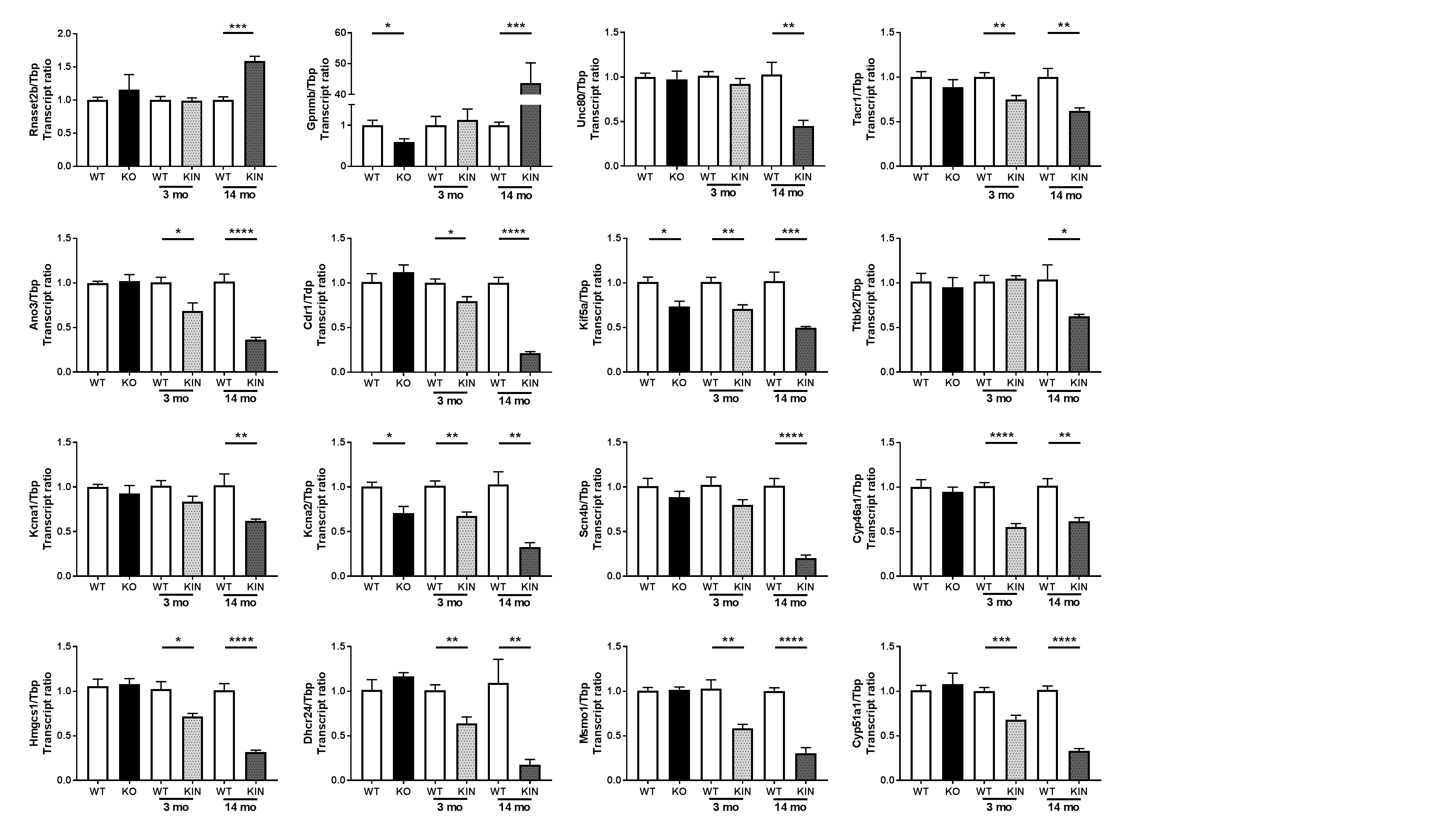
