## Supplemental Figure S7 for "*Atxn2*-CAG100-KnockIn mouse spinal cord shows progressive TDP43 pathology associated with cholesterol biosynthesis suppression"

### TYROBP Causal Network

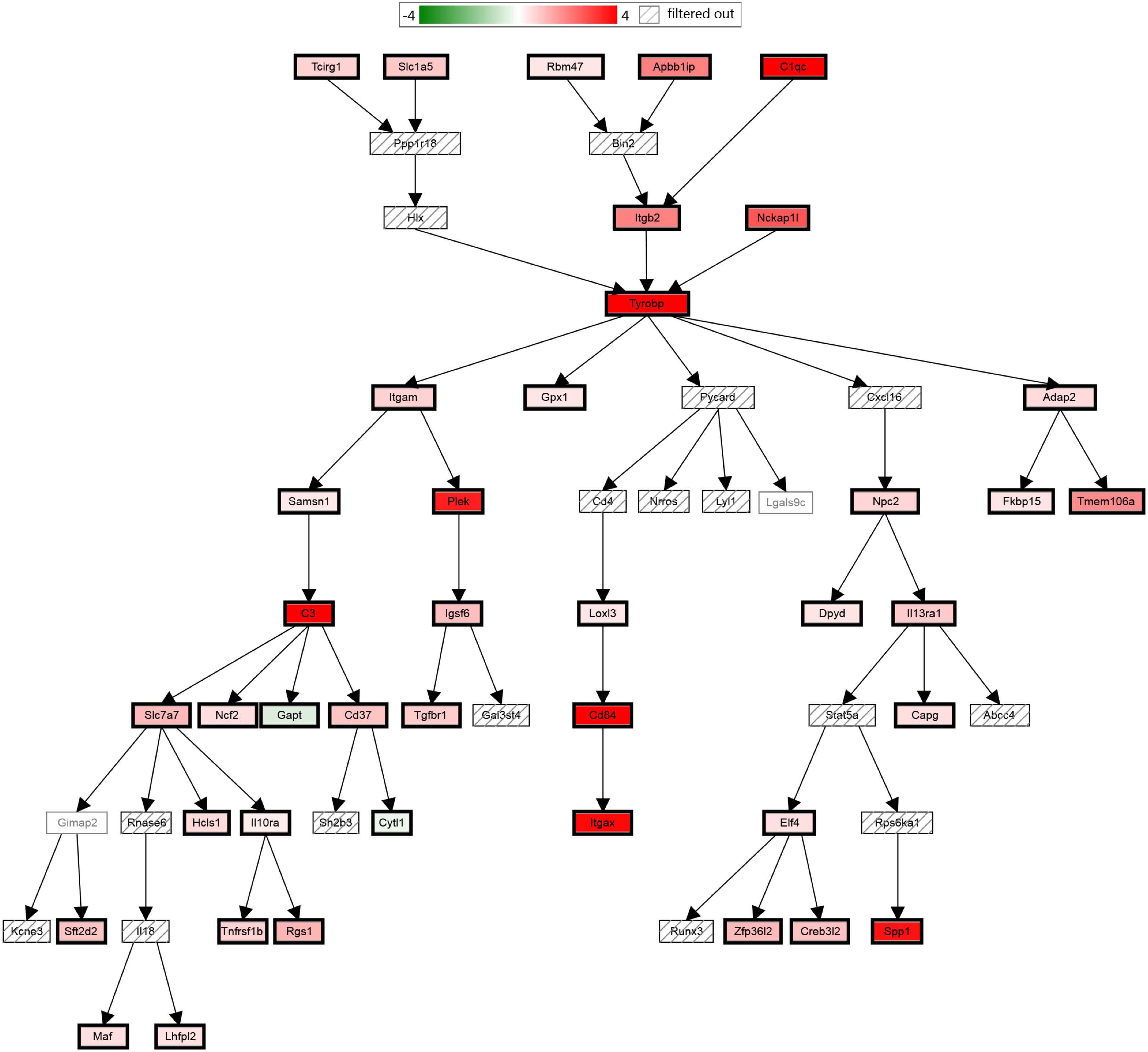

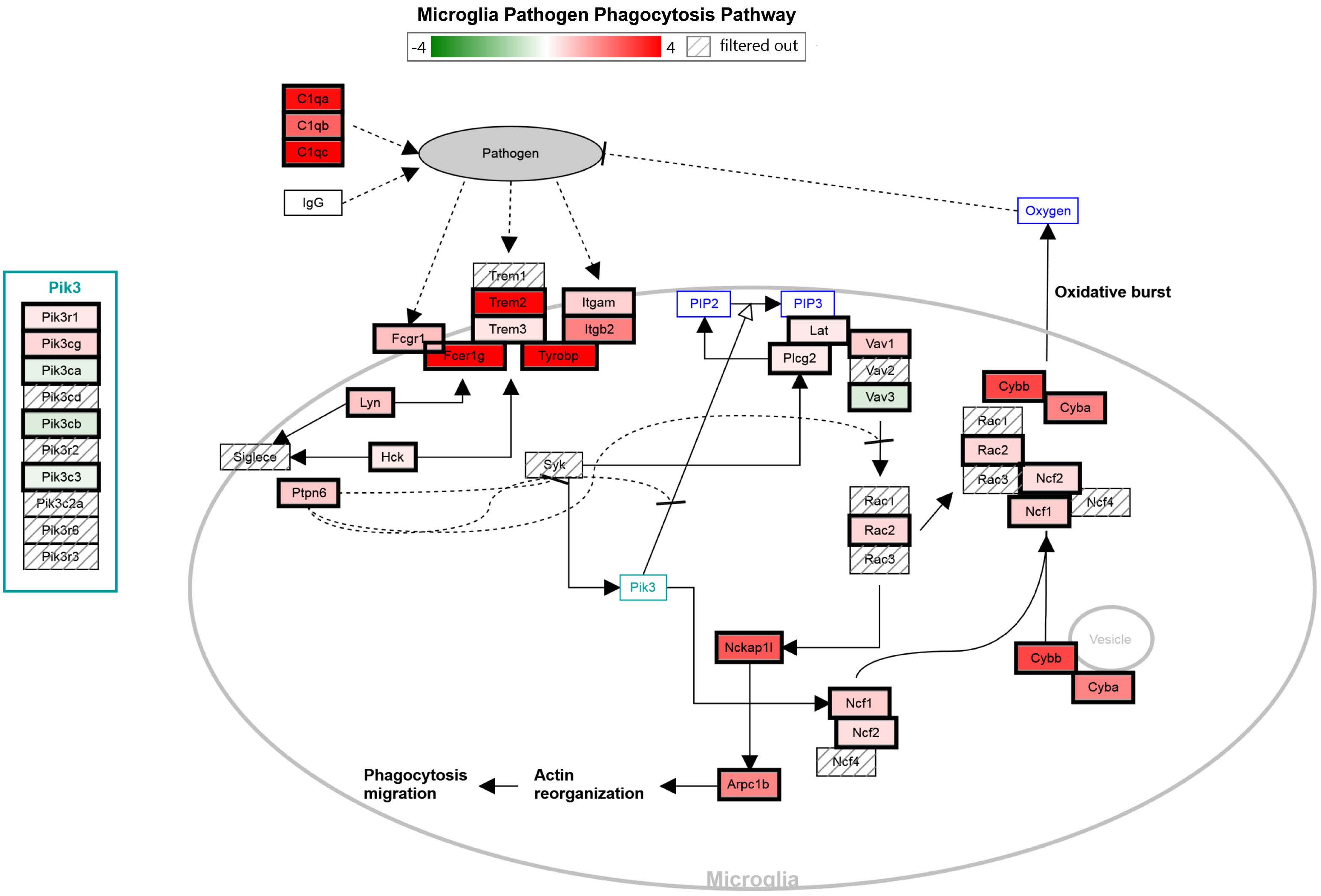
Microglia Pathogen Phagocytosis Pathway
-4
4
filtered out
C1qa
C1qb
C1qc
IgG
Pathogen
Oxygen
Oxidative burst
Pik3
Pik3r1
Pik3cg
Pik3ca
Pik3cd
Pik3cb
Pik3r2
Pik3c3
Pik3c2a
Pik3r6
Pik3r3
Trem1
Trem2
Trem3
Itgam
Itgb2
Fcgr1
Fcer1g
Tyrobp
PIP2
PIP3
Lat
Plcg2
Vav1
Vav2
Vav3
Cybb
Cyba
Rac1
Rac2
Rac3
Ncf2
Ncf1
Ncf4
Lyn
Hck
Siglece
Ptpn6
Syk
Rac1
Rac2
Rac3
Pik3
Nckap1l
Ncf1
Ncf2
Ncf4
Vesicle
Cybb
Cyba
Arpc1b
Phagocytosis
migration
Actin
reorganization
Microglia

### B Cell Receptor Signaling Pathway

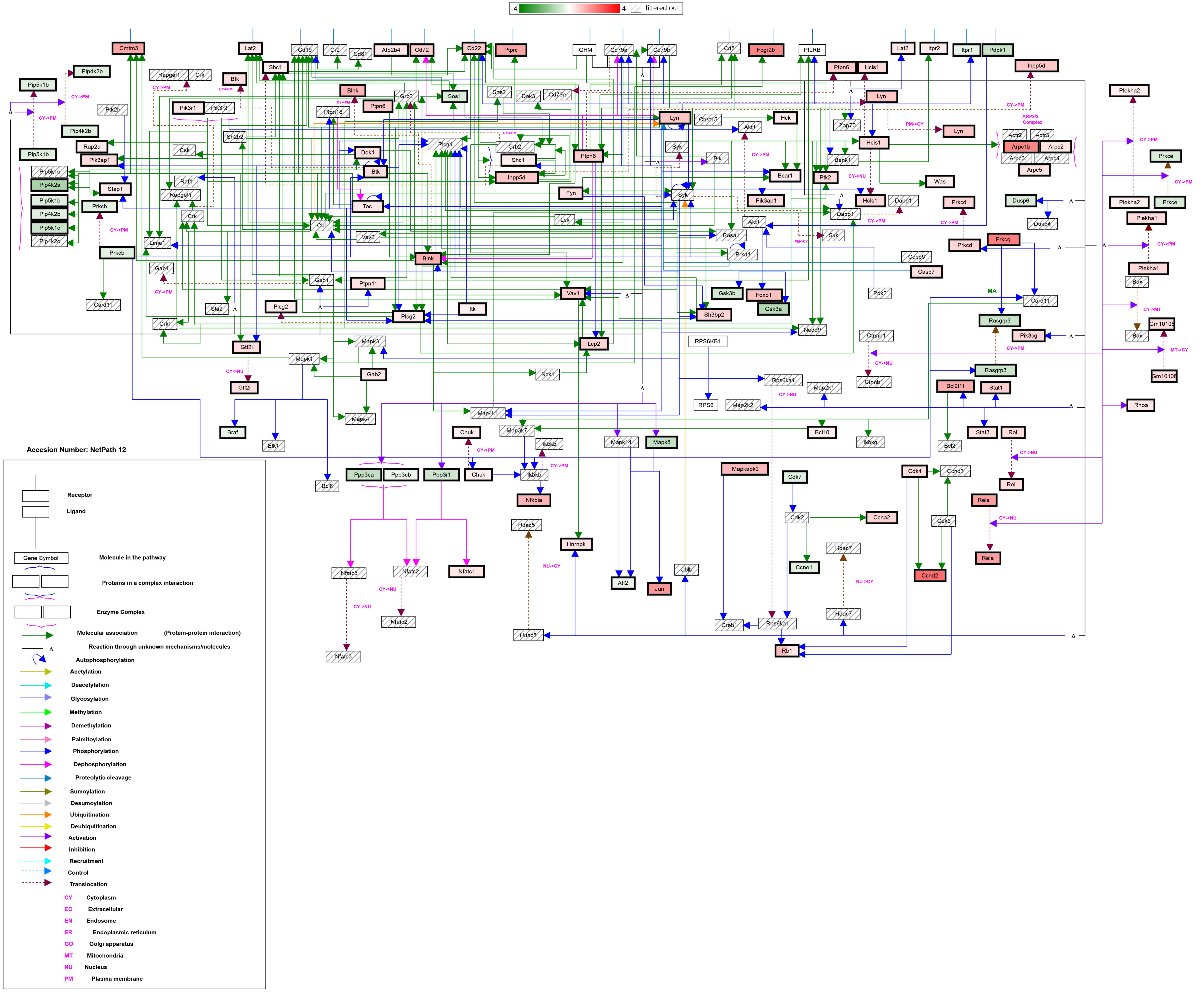

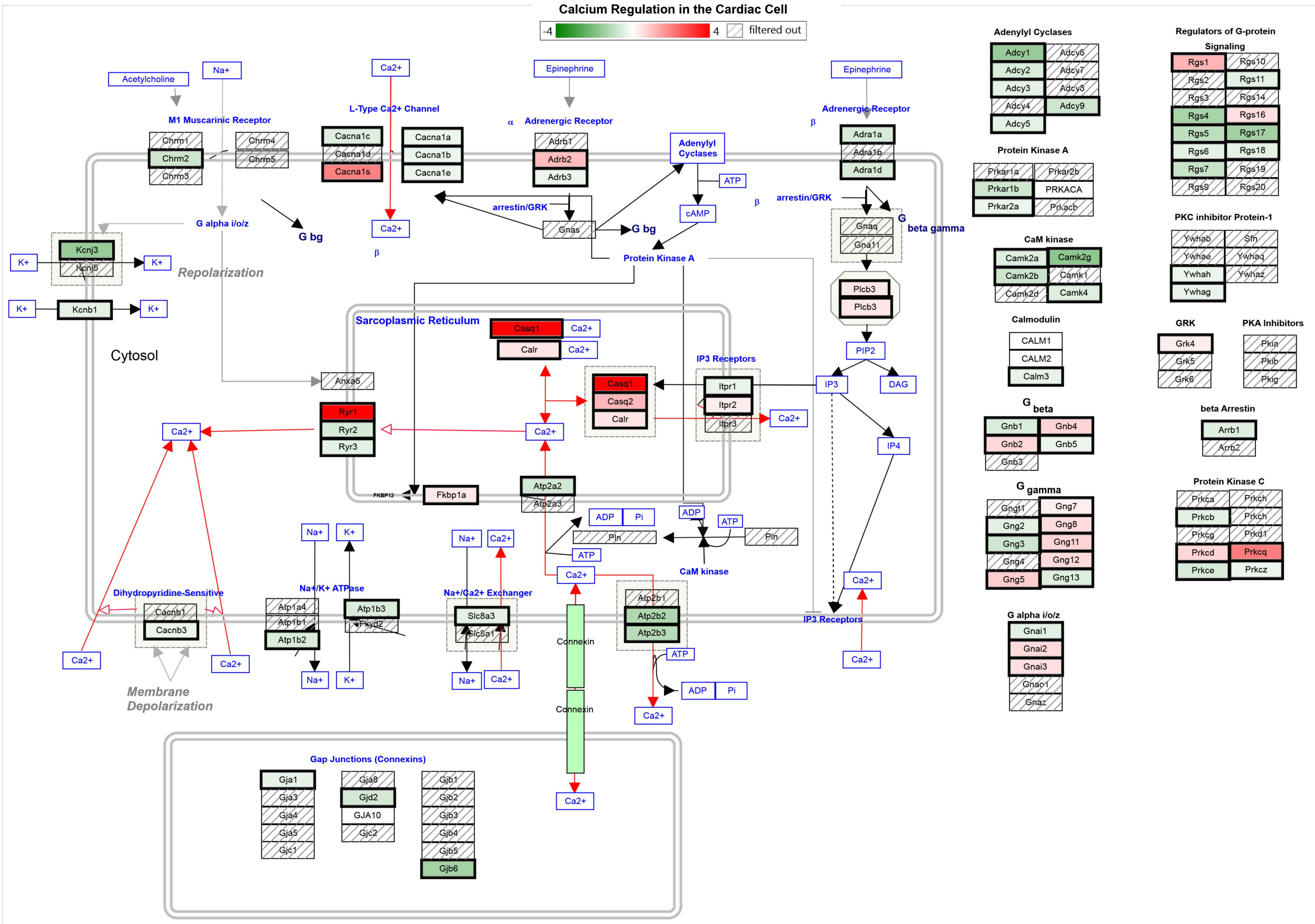
Calcium Regulation in the Cardiac Cell
-4
4
filtered out
Acetylcholine
Na+
M1 Muscarinic Receptor
Chrm1
Chrm2
Chrm3
Chrm4
Chrm5
Ca2+
L-Type Ca2+ Channel
Cacna1c
Cacna1d
Cacna1s
Cacna1a
Cacna1b
Cacna1e
Epinephrine
α Adrenergic Receptor
Adrb1
Adrb2
Adrb3
Adenylyl Cyclases
ATP
cAMP
Epinephrine
Adrenergic Receptor
β
Adra1a
Adra1b
Adra1d
arrestin/GRK
Gnas
G bg
Protein Kinase A
β arrestin/GRK
Gnaq
Gna11
G beta gamma
Plcb3
Plcb3
PIP2
IP3
DAG
IP4
G alpha i/o/z
G bg
K+
Kcnj3
Kcnj5
K+
Repolarization
K+
Kcnb1
K+
Cytosol
Sarcoplasmic Reticulum
Casq1
Ca2+
Calr
Ca2+
IP3 Receptors
Itpr1
Itpr2
Itpr3
Casq1
Casq2
Calr
Ca2+
Anxa6
Ryr1
Ryr2
Ryr3
Ca2+
Ca2+
Atp2a2
Atp2a3
FKBP12
Fkbp1a
ADP
Pi
ADP
ATP
Pln
Pln
ATP
CaM kinase
Ca2+
Na+
K+
Na+
Ca2+
Na+/K+ ATPase
Atp1a4
Atp1b1
Atp1b2
Atp1b3
Fxyd2
Na+/Ca2+ Exchanger
Slc8a3
Slc8a1
Dihydropyridine-Sensitive
Cacnb1
Cacnb3
Ca2+
Ca2+
Membrane Depolarization
Na+
K+
Na+
Ca2+
Connexin
Connexin
Atp2b1
Atp2b2
Atp2b3
ATP
ADP
Pi
Ca2+
Ca2+
IP3 Receptors
Ca2+
Ca2+
Gap Junctions (Connexins)
Gja1
Gja3
Gja4
Gja5
Gjc1
Gja8
Gjd2
GJA10
Gjc2
Gjb1
Gjb2
Gjb3
Gjb4
Gjb5
Gjb6
Adenylyl Cyclases
Adcy1
Adcy2
Adcy3
Adcy4
Adcy5
Adcy6
Adcy7
Adcy8
Adcy9
Protein Kinase A
Prkar1a
Prkar1b
Prkar2a
Prkar2b
PRKACA
Prkacb
CaM kinase
Camk2a
Camk2b
Camk2d
Camk2g
Camk1
Camk4
Calmodulin
CALM1
CALM2
Calm3
G beta
Gnb1
Gnb2
Gnb3
Gnb4
Gnb5
G gamma
Gngt1
Gng2
Gng3
Gng4
Gng5
Gng7
Gng8
Gng11
Gng12
Gng13
G alpha i/o/z
Gnai1
Gnai2
Gnai3
Gnao1
Gnaz
Regulators of G-protein Signaling
Rgs1
Rgs2
Rgs3
Rgs4
Rgs5
Rgs6
Rgs7
Rgs9
Rgs10
Rgs11
Rgs14
Rgs16
Rgs17
Rgs18
Rgs19
Rgs20
PKC inhibitor Protein-1
Ywhab
Ywhae
Ywhah
Ywhag
Sfn
Ywhaq
Ywhaz
GRK
Grk4
Grk5
Grk6
PKA Inhibitors
Pkia
Pkib
Pkig
beta Arrestin
Arrb1
Arrb2
Protein Kinase C
Prkca
Prkcb
Prkcg
Prkcd
Prkce
Prkch
Prkch
Prkd1
Prkcq
Prkcz

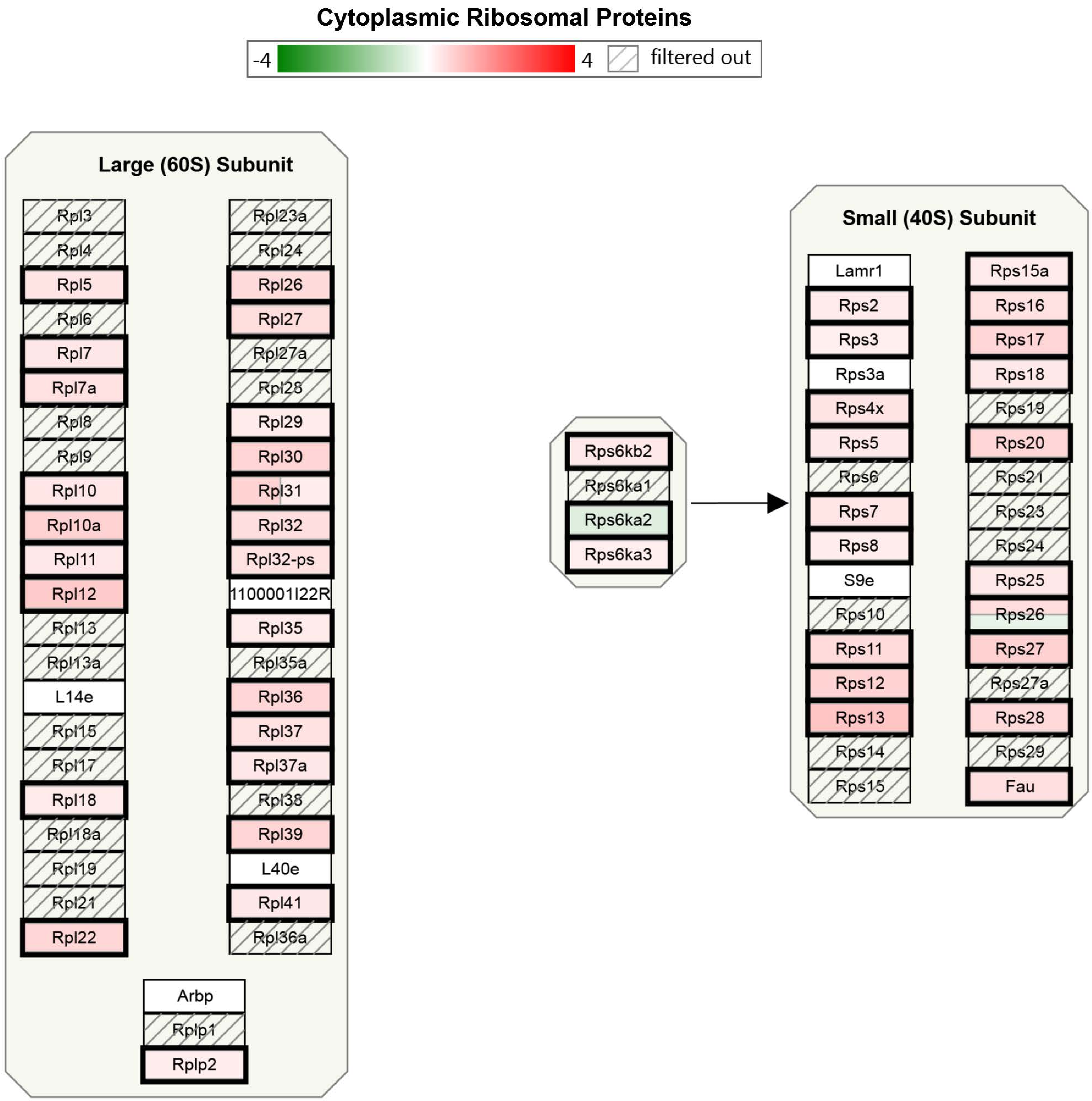
Cytoplasmic Ribosomal Proteins
-4
4
filtered out
Large (60S) Subunit
Rpl3
Rpl4
Rpl5
Rpl6
Rpl7
Rpl7a
Rpl8
Rpl9
Rpl10
Rpl10a
Rpl11
Rpl12
Rpl13
Rpl13a
L14e
Rpl15
Rpl17
Rpl18
Rpl18a
Rpl19
Rpl21
Rpl22
Rpl23a
Rpl24
Rpl26
Rpl27
Rpl27a
Rpl28
Rpl29
Rpl30
Rpl31
Rpl32
Rpl32-ps
1100001I22R
Rpl35
Rpl35a
Rpl36
Rpl37
Rpl37a
Rpl38
Rpl39
L40e
Rpl41
Rpl36a
Arbp
Rplp1
Rplp2
Rps6kb2
Rps6ka1
Rps6ka2
Rps6ka3
Small (40S) Subunit
Lamr1
Rps2
Rps3
Rps3a
Rps4x
Rps5
Rps6
Rps7
Rps8
S9e
Rps10
Rps11
Rps12
Rps13
Rps14
Rps15
Rps15a
Rps16
Rps17
Rps18
Rps19
Rps20
Rps21
Rps23
Rps24
Rps25
Rps26
Rps27
Rps27a
Rps28
Rps29
Fau

#### Cholesterol metabolism (includes both Bloch and Kandutsch-Russell pathways)

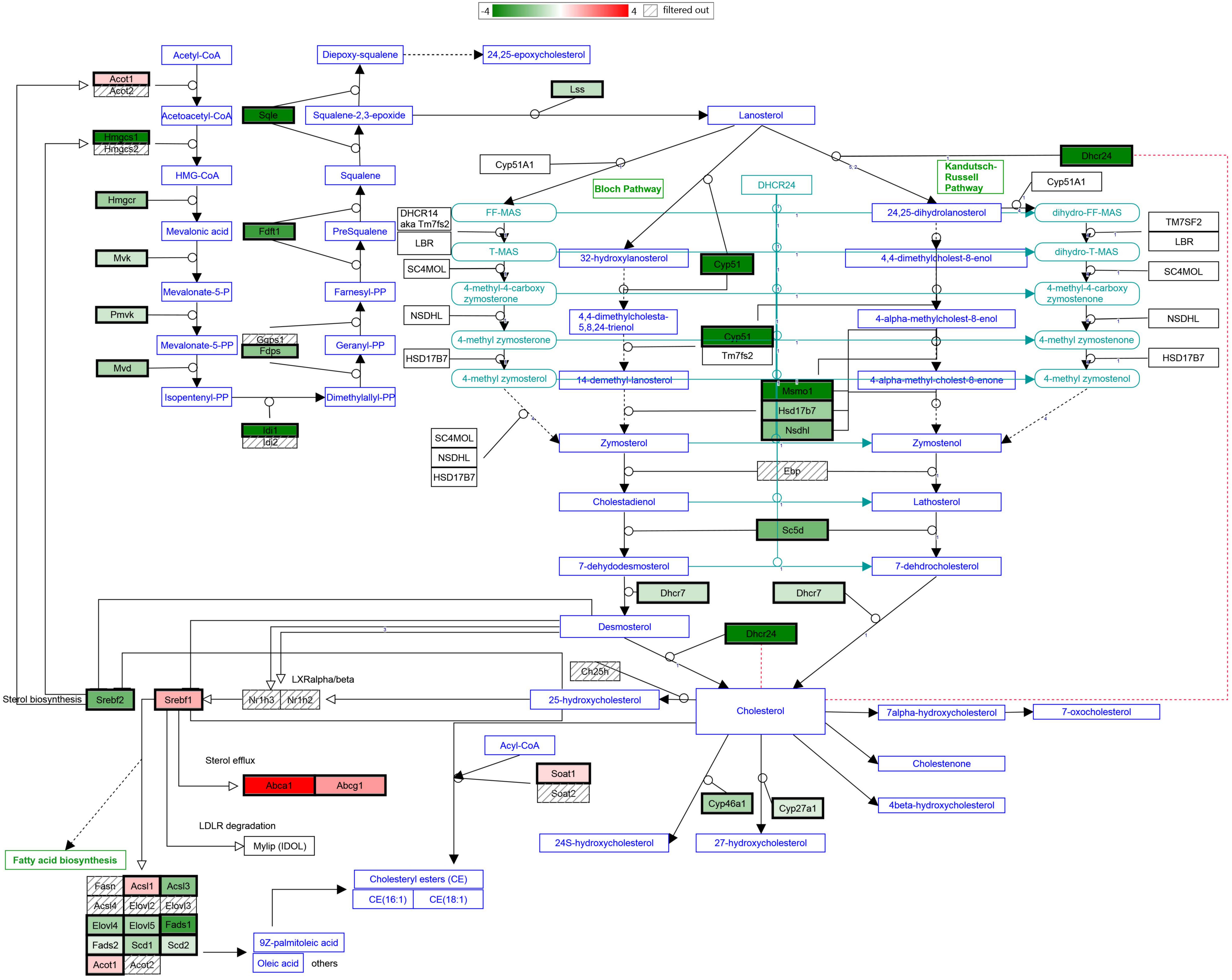

#### Regulation of Actin Cytoskeleton

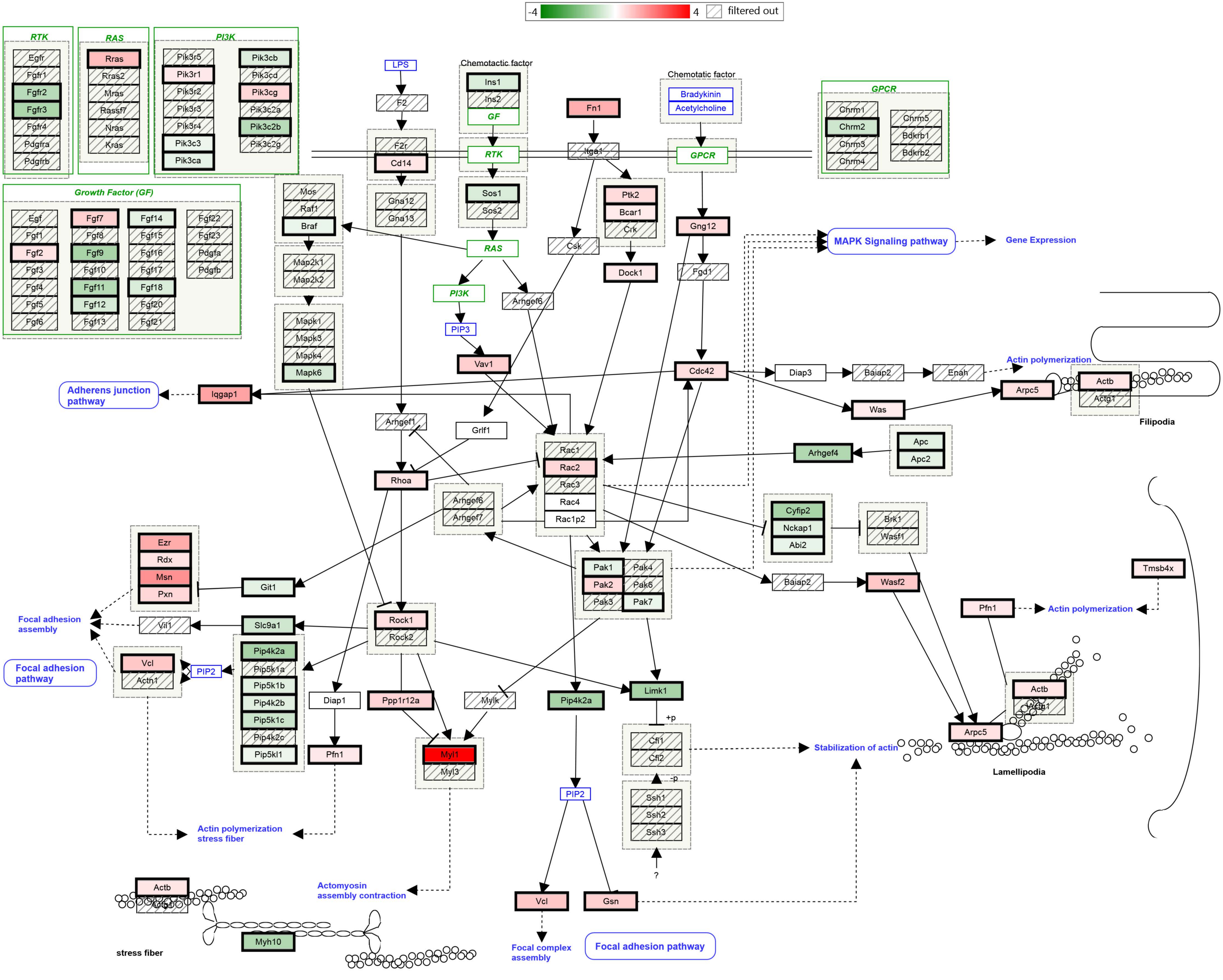

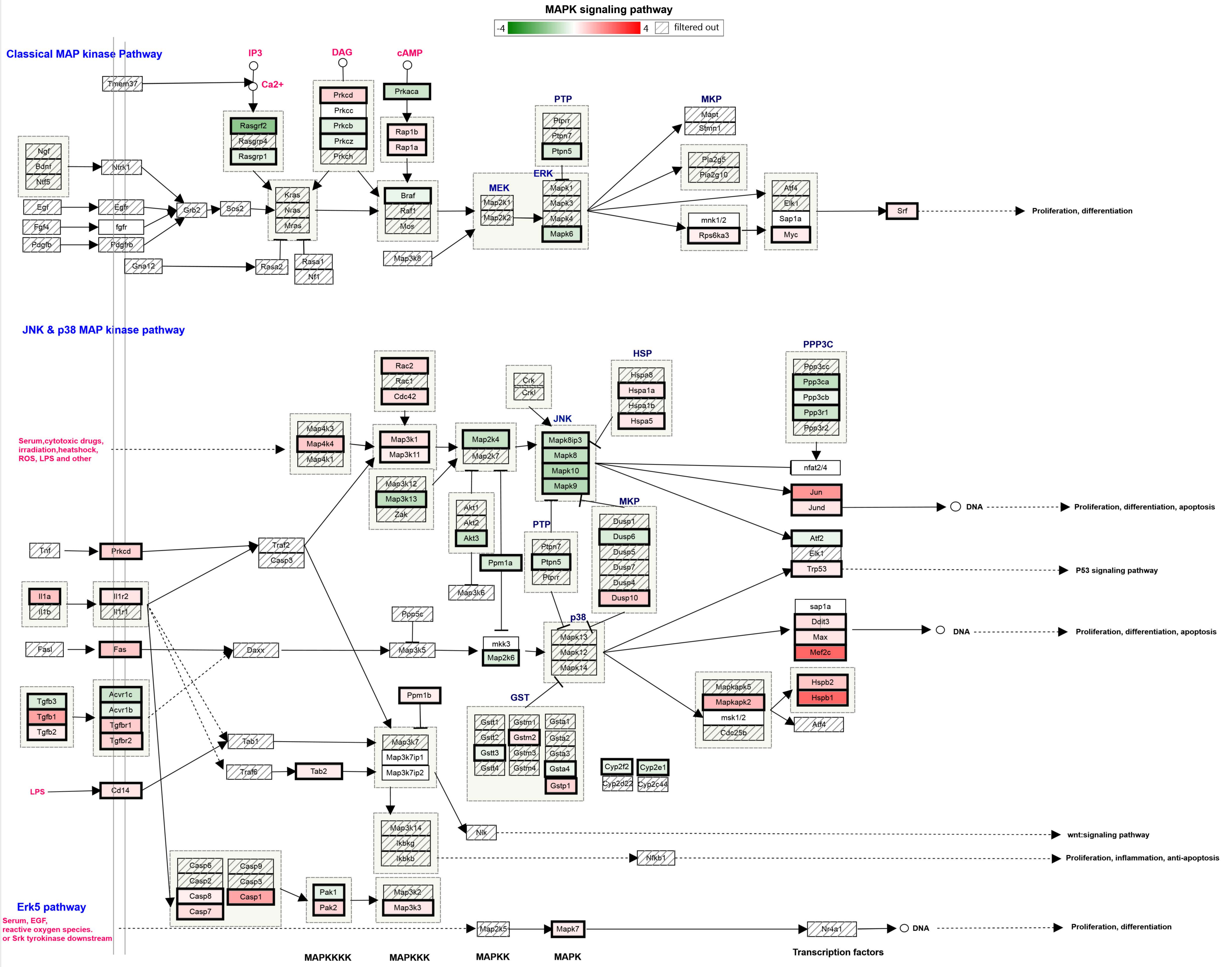

MAPK signaling pathway
-4
4
filtered out
Classical MAP kinase Pathway
IP3
DAG
cAMP
Ca2+
Tmem37
Prkcd
Prkcc
Prkcb
Prkcz
Prkch
Prkaca
Rasgrf2
Rasgrp4
Rasgrp1
Rap1b
Rap1a
Ngf
Bdnf
Ntf5
Ntrk1
Egf
Egfr
Fgf4
fgfr
Pdgfb
Pdgfrb
Grb2
Sos2
Kras
Nras
Mras
Braf
Raf1
Mos
Gna12
Rasa2
Rasa1
Nf1
Map3k8
MEK
Map2k1
Map2k2
PTP
Ptprr
Ptpn7
Ptpn5
ERK
Mapk1
Mapk3
Mapk4
Mapk6
MKP
Mapt
Stmn1
Pla2g5
Pla2g10
mnk1/2
Rps6ka3
Atf4
Elk1
Sap1a
Myc
Srf
Proliferation, differentiation
JNK & p38 MAP kinase pathway
Serum,cytotoxic drugs, irradiation,heatshock, ROS, LPS and other
Rac2
Rac1
Cdc42
Map4k3
Map4k4
Map4k1
Map3k1
Map3k11
Map3k12
Map3k13
Zak
Map2k4
Map2k7
Crk
Crkl
JNK
Mapk8ip3
Mapk8
Mapk10
Mapk9
HSP
Hspa8
Hspa1a
Hspa1b
Hspa5
PPP3C
Ppp3cc
Ppp3ca
Ppp3cb
Ppp3r1
Ppp3r2
nfat2/4
Jun
Jund
DNA
Proliferation, differentiation, apoptosis
Akt1
Akt2
Akt3
Ppm1a
Map3k6
PTP
Ptpn7
Ptpn5
Ptprr
MKP
Dusp1
Dusp6
Dusp5
Dusp7
Dusp4
Dusp10
Atf2
Elk1
Trp53
P53 signaling pathway
Tnf
Prkcd
Traf2
Casp3
Il1a
Il1b
Il1r2
Il1r1
Fasl
Fas
Daxx
Ppp5c
Map3k5
mkk3
Map2k6
p38
Mapk13
Mapk12
Mapk14
sap1a
Ddit3
Max
Mef2c
DNA
Proliferation, differentiation, apoptosis
Tgfb3
Tgfb1
Tgfb2
Acvr1c
Acvr1b
Tgfbr1
Tgfbr2
Ppm1b
GST
Gstt1
Gstt2
Gstt3
Gstt4
Gstm1
Gstm2
Gstm3
Gstm4
Gsta1
Gsta2
Gsta3
Gsta4
Gstp1
Cyp2f2
Cyp2e1
Cyp2d22
Cyp2c44
Mapkapk5
Mapkapk2
msk1/2
Cdc25b
Hspb2
Hspb1
Atf4
Tab1
Map3k7
Map3k7ip1
Map3k7ip2
Traf6
Tab2
LPS
Cd14
Nlk
wnt:signaling pathway
Map3k14
Ikbkg
Ikbkb
Nfkb1
Proliferation, inflammation, anti-apoptosis
Casp6
Casp2
Casp8
Casp7
Casp9
Casp3
Casp1
Pak1
Pak2
Map3k2
Map3k3
Erk5 pathway
Serum, EGF, reactive oxygen species. or Srk tyrokinase downstream
Map2k5
Mapk7
Nr4a1
DNA
Proliferation, differentiation
MAPKKKK
MAPKKK
MAPKK
MAPK
Transcription factors

### Insulin Signaling

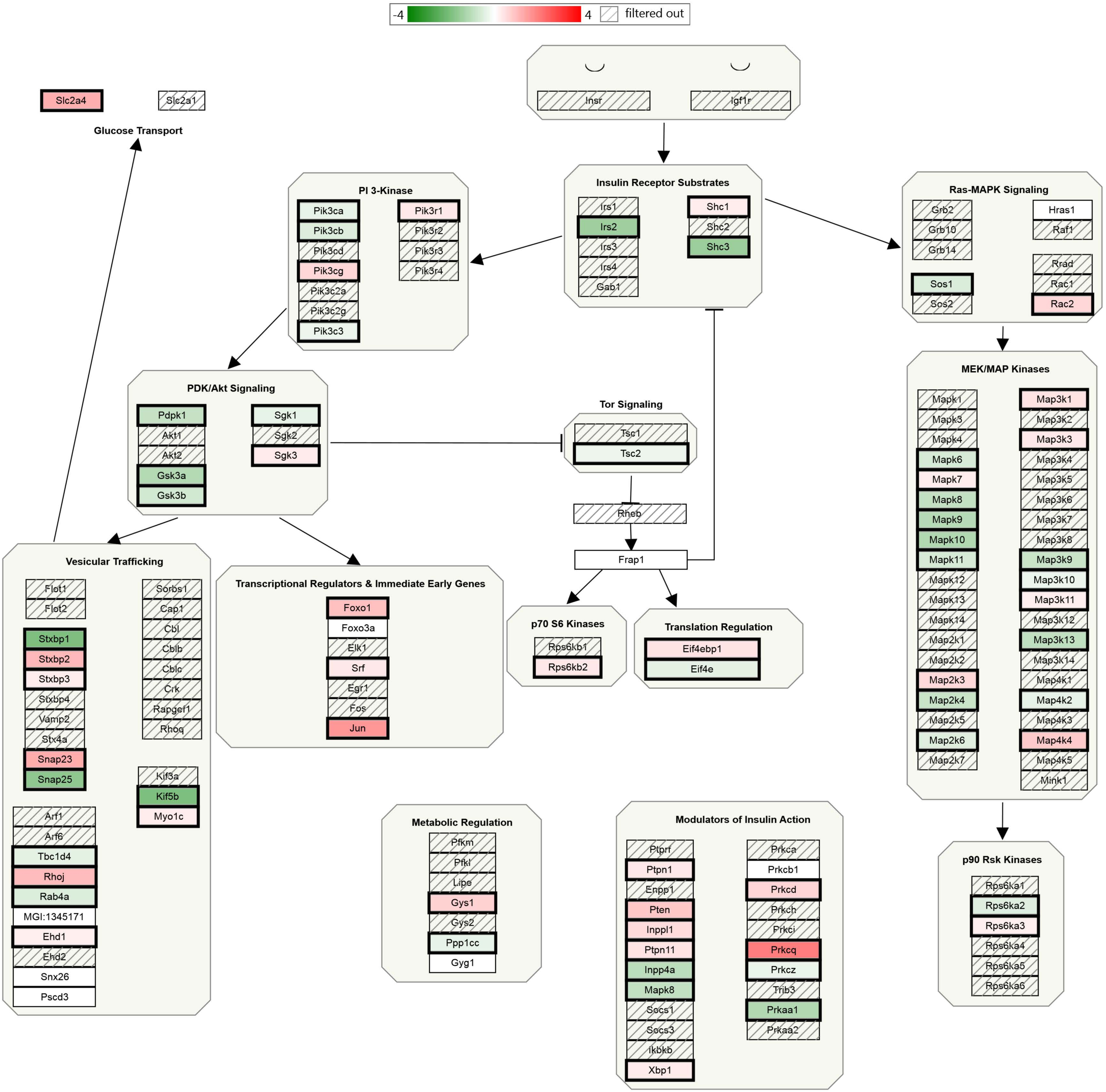

Focal Adhesion-PI3K-Akt-mTOR-signaling pathway

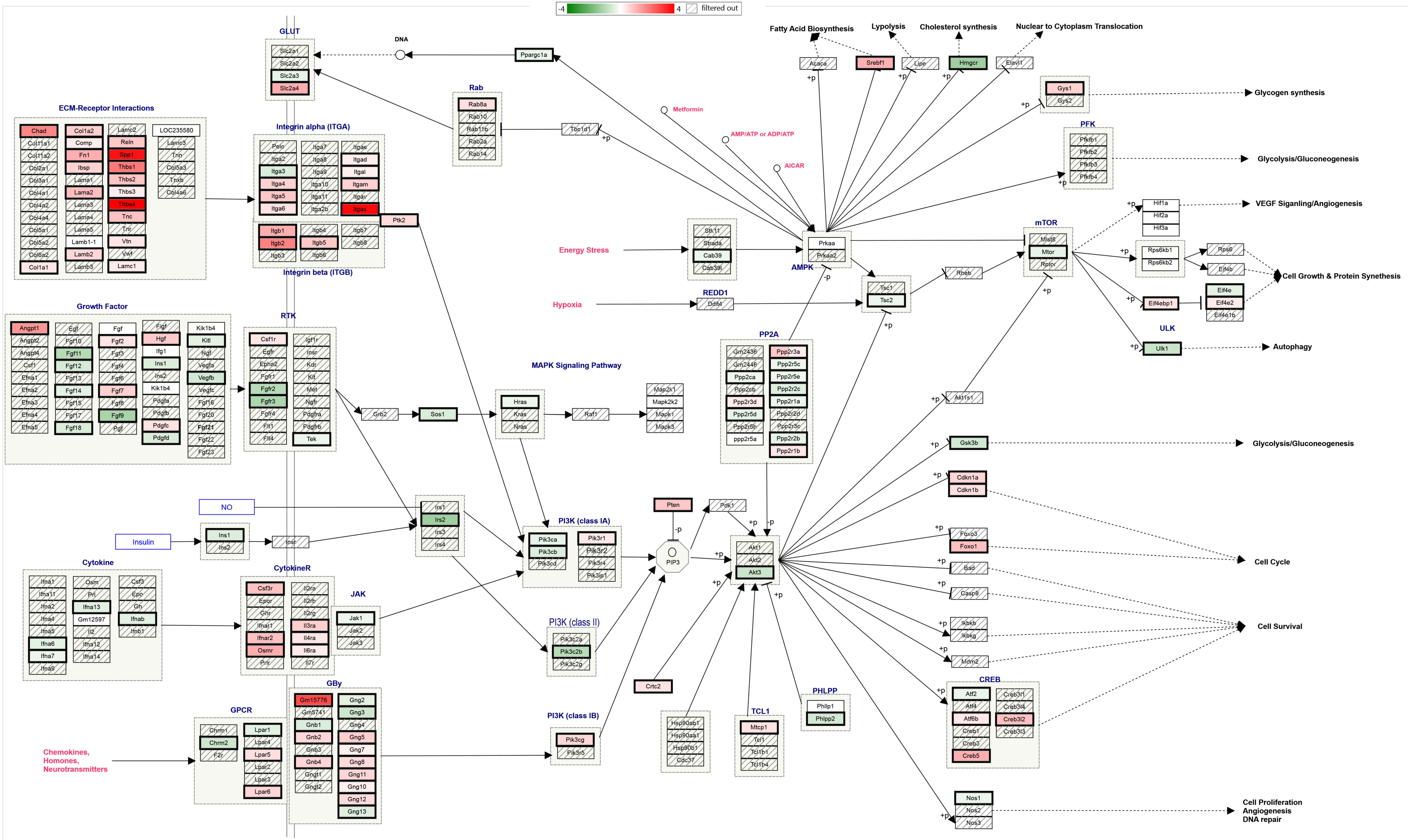

### EGFR1 Signaling Pathway

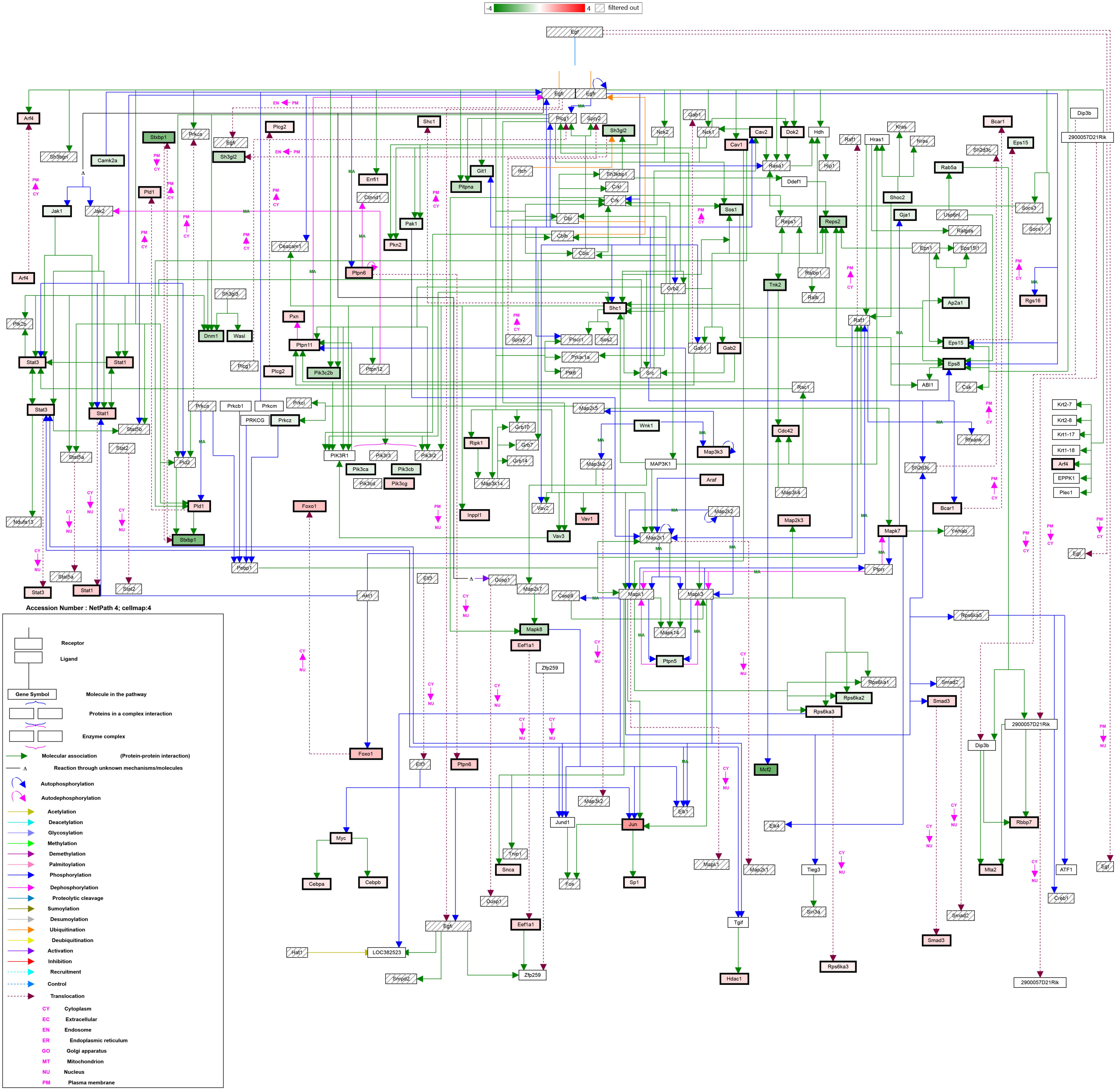

### mRNA processing

-4 4 filtered out

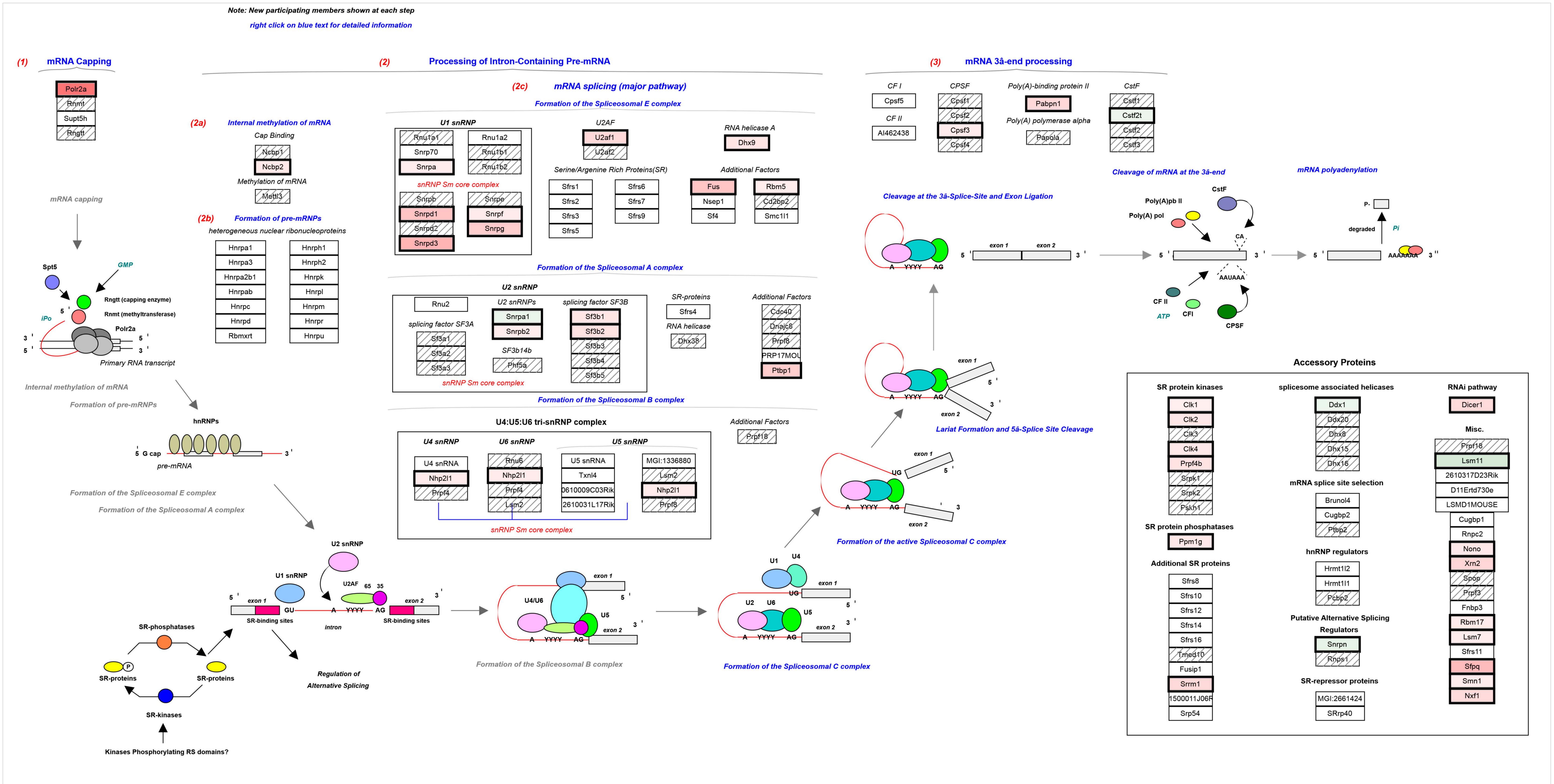


**InterPro Domain Containing:** **RNA-binding region RNP-1 (RNA recognition motif)**

| | | | | | | | | |
|---|---|---|---|---|---|---|---|---|
| 0610009D07Rik | 2410104I19Rik | 3830425H19Rik | A130095G20Rik | AW049977 | Cyp4f18 | Matr3-ps1 | Rbms2 | U2af1-rs2 |
| 0610009J05Rik | 2410112D09Rik | 4632417O19Rik | AA407033 | AW107884 | Cpeb1 | Matr3-ps2 | Rbmx | V1rc17 |
| 0610033I05Rik | 2600011C06Rik | Hnrpf | AA522010 | AW260507 | Cpeb3 | Mki67ip | Rbmxrt | Wdr9 |
| 0710005M24Rik | Rbm13 | 4921506I22Rik | AA589495 | AW553050 | Cpeb4 | Spen | Rbmy1a1 | Wbscr1 |
| Ilkap | 2600016C11Rik | Nol8 | AI043106 | BB131279 | Cova1 | Msi1h | Rbpms | |
| Rnpepl1 | 2600016F06Rik | 4922503N01Rik | AI194270 | BC013481 | D1Pas1 | Msi2h | Rdbp | |
| 1110034O07Rik | Htatsf1 | 4930422M05Rik | Safb2 | Dnd1 | Dazap1 | Myef2 | Thoc4 | |
| 1110037F02Rik | 2610015J01Rik | Lrrn6a | AI255183 | B230118G17Rik | Dazl | Syncrip | Refbp2 | |
| Rbm19 | Sfrs11 | 4930506D01Rik | AI255207 | B230333C21Rik | MGI:2387367 | Nsap1l-pending | Rod1 | |
| 1200009B18Rik | 2610020N02Rik | 4930554P13Rik | AI255212 | B930007L02Rik | D15Ertd119e | Zfml | Rnpc1 | |
| 1300006N24Rik | 2610023M21Rik | 4930562C03Rik | AI255215 | C230097K14Rik | D5Ertd700e | Ncl | Rnpep | |
| Rbm7 | 2610024A01Rik | 4930565A21Rik | AI255219 | Rbed1 | D8Ertd233e | Pspc1 | Rnu12 | |
| 1500012C14Rik | 2610101N10Rik | Boll | AI255230 | C330027G06Rik | Elavl1 | Ppil4 | Rnu108 | |
| 1600023H17Rik | Bclaf1 | 4932422E22Rik | D9Mgi24 | C430048L16Rik | Elavl2 | Ppie | Rnu11 | |
| 1700003D06Rik | 2610102M01Rik | 4932422M17Rik | AI255750 | C79783 | Elavl3 | Ppargc1a | Rnu1a1 | |
| 1700009P03Rik | 2610209F03Rik | 4932702K14Rik | AI255840 | C80672 | Elavl4 | Ppargc1b | Rnu1b1 | |
| 1700012H05Rik | Hnrpa3 | Rbm31y | AI255847 | C85084 | Eral1 | Pabpc1 | Rnu1b2 | |
| 1700025B16Rik | Rbm17 | 5031411E02Rik | AI256346 | Zfp622 | Eif3s4 | Pabpc2 | Rnu1b6 | |
| 1700047G05Rik | 2700028P07Rik | 5330419I02Rik | AI256354 | D19Bwg1357e | Eif3s9 | Pabpc4 | Sfrs10 | |
| 1700095D18Rik | Rbm21 | 5730453I16Rik | AI314192 | D230005D02Rik | Eif4g2 | Pabpc5 | Ssb | |
| Rbm15b | 2700077B20Rik | Tdrd7 | AI447568 | D330023I21Rik | Ewsr1 | Poldip3 | Sbno1 | |
| 1810035L17Rik | 2700088M22Rik | 5730555F13Rik | Sfrs12 | D830027H13Rik | G3bp2-pending | Pcolce | Slc25a4 | |
| 1810036A22Rik | Hnrpll | 5730557B15Rik | AI462171 | G430095G15Rik | MGI:1351465 | Ppp1r14b | Slc6a8 | |
| 1810036E18Rik | 2810036M01Rik | 5730557H03Rik | AI467299 | Adam5 | Ggcx | Rbm3 | Sart3 | |
| Rbm18 | Rbmx2 | 6330548G22Rik | AI503051 | Afg3l1 | Grsf1 | Rbm4 | Synj2 | |
| 2010300K22Rik | 2810411C16Rik | Brunol6 | AI504630 | Acin1 | Hdp | Rbm6 | Taf15 | |
| 2010319N22Rik | 2810441O16Rik | Rbms3 | AI642080 | MGI:1926224 | Hnrpdl | Rbm8a | Tardbp | |
| 2210008M09Rik | 2810442I21Rik | 6720478K01Rik | AI843301 | Abcb4 | H2-Bf | Rbm9 | Tia1 | |
| Afg3l2 | Rbm28 | 8030431D03Rik | AI844780 | Cnot4 | Raly | Rbm10 | Tiaf1 | |
| 2310046H11Rik | 3000004N20Rik | Rbm22 | AL022933 | Cstf2t | Htf9c | Rbm11 | Tial1 | |
| Rbpms2 | Hnrpa0 | 9130414A06Rik | D12Mgi43 | Cirbp | Igf2bp1 | Rbm12 | Traf6 | |
| Rad52b | 3100004P22Rik | 9430070C08Rik | D12Mgi40 | Cdk9 | Igf2bp3 | Rbm14 | Tlr5 | |
| 2410013L13Rik | 3110061K15Rik | 9530027K23Rik | AU023006 | Col4a3 | Uhmk1 | Rbm16 | Tnrc4 | |
| Wdr55 | 3200001K10Rik | Tnrc6c | AU045931 | Csad | Matr3 | Rbms1 | U2af1-rs1 | |

**Gene Ontology: RNA Binding**

| | | | | |
|---|---|---|---|---|
| Rnaset2 | Adar | Ilf3 | Rpl22 | Tert |
| Rpl38 | Adarb2 | Lgtn | Rpl26 | Trim21 |
| 1110017C15Rik | Adat1 | Lsm4 | Rpl32 | Tsn |
| 1110033J19Rik | Akap1 | Mbnl1 | Rpl37 | Wbp4 |
| 1190002L16Rik | Auh | Mrps28 | Rpl39 | Zbp1 |
| 1190005F20Rik | Bard1 | Mrpl11 | Rpl7 | Zfp346 |
| Eif4e3 | Bat1a | Mrpl23 | Rpl8 | Zfp385 |
| Eif4g3 | Bicc1 | Npm1 | Rpl9 | |
| Lsm8 | Brca1 | Nufip1 | Rps11 | |
| 2010003O18Rik | Ciz1 | Nxf7 | Rps13 | |
| Rpl11 | D1Pas1-rs1 | Oas1a | Rps14 | |
| Exosc8 | Imp4 | Oas1b | Rps20 | |
| 2400002F11Rik | D3Jfr1 | Oas1g | Rps23 | |
| Rnaseh2a | Exosc5 | Oas2 | Rps24 | |
| 2410003C20Rik | Ddx19a | Oas3 | Rps27 | |
| Exosc7 | Ddx21 | Oasl2 | Rps28 | |
| Srp68 | Ddx24 | Papolb | Rps29 | |
| 2610027L16Rik | Ddx25 | Park7 | Rps4x | |
| Zmat2 | Ddx39 | Pcbp1 | Rps6 | |
| Rpp14 | Ddx3x | Pcbp3 | Rps7 | |
| Ttc14 | Ddx4 | Pcbp4 | Exosc4 | |
| Secisbp2 | Ddx41 | Exosc9 | Sca1 | |
| Lsm1 | Ddx48 | Polr2g | Scye1 | |
| 2810028N01Rik | Ddx5 | Prkr | Slbp | |
| Ddx19b | Ddx56 | 2810055E05Rik | Snrp1c | |
| Rps9 | Ddx6 | Pum1 | Son | |
| Rnmtl1 | Dkc1 | Pum2 | Srp14 | |
| 4930429A22Rik | Eif4a2 | Qk | Srp19 | |
| 4930517K11Rik | Eif4e | Rad21 | Srp9 | |
| 4930534J06Rik | Eif4e2 | Rae1 | Ssa2 | |
| Ppp1r8 | Fbl | Rnasel | Nhp2l1 | |
| Papolg | Fmr1 | Rnu2 | Stau1 | |
| 9630048M01Rik | Fxr1h | Rpl12 | Tarbp2 | |
| Aco1 | Fxr2h | Rpl19 | Tenr | |

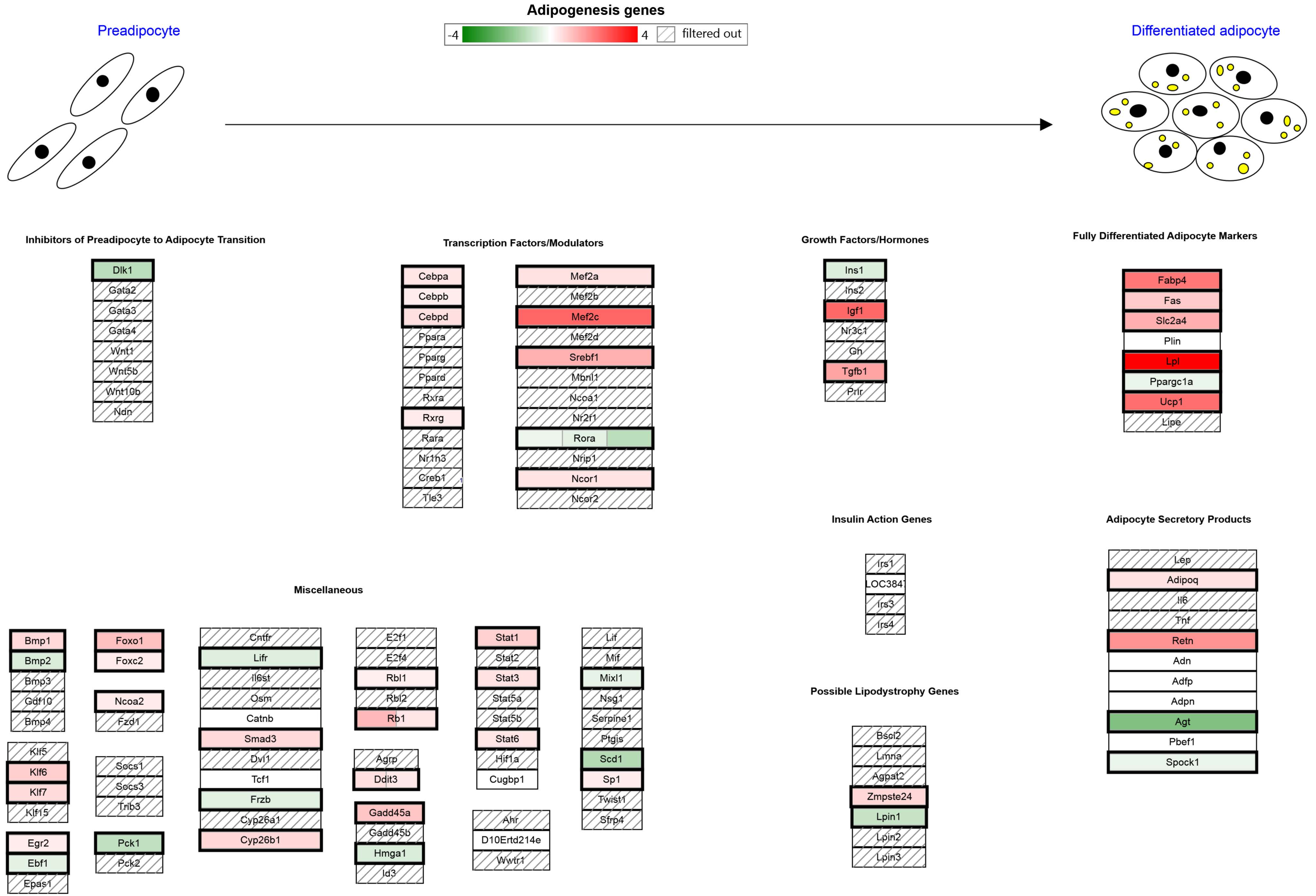
Adipogenesis genes
-4
4
filtered out
Preadipocyte
Differentiated adipocyte
Inhibitors of Preadipocyte to Adipocyte Transition
Dlk1
Gata2
Gata3
Gata4
Wnt1
Wnt5b
Wnt10b
Ndn
Transcription Factors/Modulators
Cebpa
Cebpb
Cebpd
Ppara
Pparg
Ppard
Rxra
Rxrg
Rara
Nr1h3
Creb1
Tle3
Mef2a
Mef2b
Mef2c
Mef2d
Srebf1
Mbnl1
Ncoa1
Nr2f1
Rora
Nrip1
Ncor1
Ncor2
Growth Factors/Hormones
Ins1
Ins2
Igf1
Nr3c1
Gh
Tgfb1
Prlr
Fully Differentiated Adipocyte Markers
Fabp4
Fas
Slc2a4
Plin
Lpl
Ppargc1a
Ucp1
Lipe
Insulin Action Genes
Irs1
LOC3847
Irs3
Irs4
Adipocyte Secretory Products
Lep
Adipoq
Il6
Tnf
Retn
Adn
Adfp
Adpn
Agt
Pbef1
Spock1
Possible Lipodystrophy Genes
Bscl2
Lmna
Agpat2
Zmpste24
Lpin1
Lpin2
Lpin3
Miscellaneous
Bmp1
Bmp2
Bmp3
Gdf10
Bmp4
Klf5
Klf6
Klf7
Klf15
Egr2
Ebf1
Epas1
Foxo1
Foxc2
Ncoa2
Fzd1
Socs1
Socs3
Trib3
Pck1
Pck2
Cntfr
Lifr
Il6st
Osm
Catnb
Smad3
Dvl1
Tcf1
Frzb
Cyp26a1
Cyp26b1
E2f1
E2f4
Rbl1
Rbl2
Rb1
Agrp
Ddit3
Gadd45a
Gadd45b
Hmga1
Id3
Stat1
Stat2
Stat3
Stat5a
Stat5b
Stat6
Hif1a
Cugbp1
Ahr
D10Ertd214e
Wwtr1
Lif
Mif
Mixl1
Nsg1
Serpine1
Ptgis
Scd1
Sp1
Twist1
Sfrp4

XPodNet - protein-protein interactions in the podocyte expanded by STRI...
PodNet: protein-protein interactions in the podocyte
Focal Adhesion-PI3K-Akt-mTOR-signaling pathway
mRNA processing
Osteoclast
PluriNetWork
Chemokine signaling pathway
Focal Adhesion
B Cell Receptor Signaling Pathway
Non-odorant GPCRs
Calcium Regulation in the Cardiac Cell
Myometrial Relaxation and Contraction Pathways
MAPK signaling pathway
MAPK signaling pathway
EGFR1 Signaling Pathway
Insulin Signaling
Regulation of Actin Cytoskeleton
GPCRs, Class A Rhodopsin-like
TGF-beta Receptor Signaling Pathway
TNF-alpha NF-kB Signaling Pathway
Adipogenesis genes
Purine metabolism
Odorant GPCRs
Integrin-mediated Cell Adhesion
T Cell Receptor Signaling Pathway
G Protein Signaling Pathways
IL-6 signaling Pathway
**Spinal Cord Injury**
Cytoplasmic Ribosomal Proteins
IL-3 Signaling Pathway
TYROBP Causal Network
Alzheimers Disease
Toll-like receptor signaling pathway
Metapathway biotransformation
GPCRs, Other
Androgen Receptor Signaling Pathway
Primary Focal Segmental Glomerulosclerosis FSGS
Wnt Signaling Pathway and Pluripotency
ESC Pluripotency Pathways
Cholesterol metabolism (includes both Bloch and Kandutsch-Russell pathw...
Striated Muscle Contraction
IL-2 Signaling Pathway
Endochondral Ossification

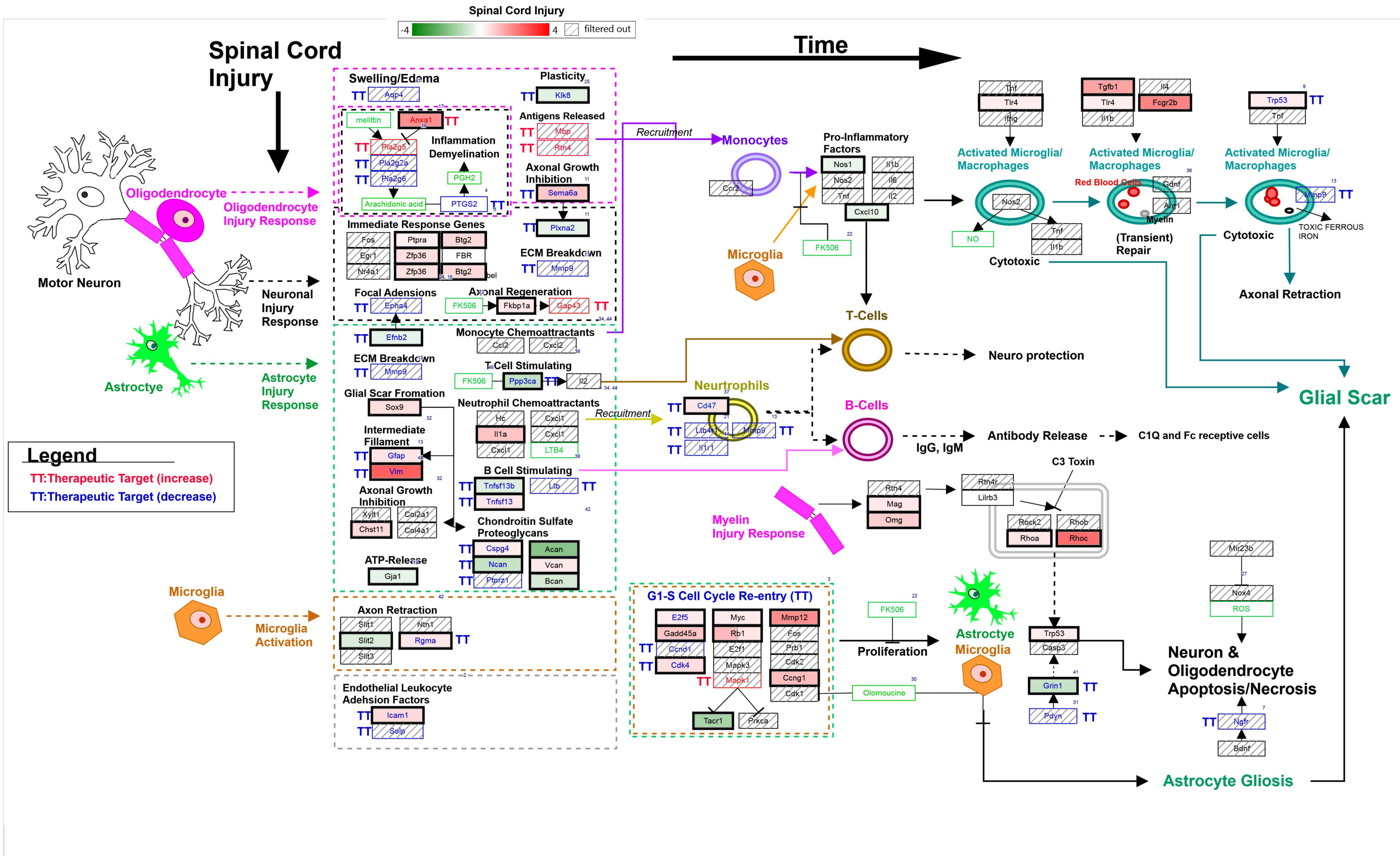
